# CRISPR/Cas13 effectors have differing extents of off-target effects that limit their utility in eukaryotic cells

**DOI:** 10.1101/2021.11.04.467323

**Authors:** Yuxi Ai, Dongming Liang, Jeremy E. Wilusz

**Affiliations:** Biochemistry and Molecular Biophysics Graduate Group, University of Pennsylvania Perelman School of Medicine, Philadelphia, PA 19104; Department of Biochemistry and Biophysics, University of Pennsylvania Perelman School of Medicine, Philadelphia, PA 19104

**Keywords:** circRNA, collateral damage, LwaCas13a, PspCas13b, RxCas13d, RNA degradation

## Abstract

CRISPR/Cas13 effectors have garnered increasing attention as easily customizable tools for detecting and depleting RNAs of interest. Near perfect complementarity between a target RNA and the Cas13-associated guide RNA is required for activation of Cas13 ribonuclease activity. Nonetheless, the specificity of Cas13 effectors in eukaryotic cells has been debated as the Cas13 nuclease domains can be exposed on the enzyme surface, providing the potential for promiscuous cleavage of nearby RNAs (so-called collateral damage). Here, using co-transfection assays in *Drosophila* and human cells, we found that the off-target effects of RxCas13d, a commonly used Cas13 effector, can be as strong as the level of on-target RNA knockdown. The extent of off-target effects is positively correlated with target RNA expression levels, and collateral damage can be observed even after reducing RxCas13d/guide RNA levels. The PspCas13b effector showed improved specificity and, unlike RxCas13d, can be used to deplete a *Drosophila* circular RNA without affecting the expression of the associated linear RNA. PspCas13b nonetheless still can have off-target effects and we notably found that the extent of off-target effects for Cas13 effectors differs depending on the cell type and target RNA examined. In total, these results highlight the need for caution when designing and interpreting Cas13-based knockdown experiments.

## INTRODUCTION

The ability to edit eukaryotic genomes and alter gene expression patterns has revolutionized biology in recent years, opening up many new opportunities for defining gene functions and developing new therapeutic approaches. In particular, CRISPR (Clustered regularly interspaced short palindromic repeats)-Cas systems, which function as defense mechanisms in prokaryotic cells against invading viruses and infectious plasmids (1,2), have been adapted as easily programmable nucleases that can be directed to specific nucleic acid sequences via complementary guide RNAs (3–5). Many CRISPR-Cas systems target DNA (e.g., the commonly used Cas9 and Cas12 systems), but Cas13 effectors target RNA and thus have the potential to manipulate the levels of RNAs of interest in cells (6–12). For the last 20 years, knockdown of cellular RNAs has generally been accomplished using RNA interference (RNAi), an approach that takes advantage of small interfering RNAs (siRNAs) or short hairpin RNAs (shRNAs) that direct an Argonaute family nuclease to catalyze cleavage of the target RNA (13,14). Unfortunately, si/shRNAs can result in depletion of additional cellular RNAs that have partial sequence complementarity, especially when the transcript is able to base pair to the seed sequence (nucleotides 2–7) of the si/shRNA (15–18). There has thus been an ongoing desire to develop novel RNA knockdown approaches that have higher specificity in cells.

Cas13 effectors associate with a guide RNA that base pairs to a target RNA via a spacer sequence that is often 22-30 nucleotides (or even longer) in length (6–9). Notably, near perfect complementarity between the guide RNA spacer and its target RNA is required for the nuclease activity of the Cas13 effector to become activated (7,19). Single point mutations in the target sequence can be sufficient to prevent knockdown by Cas13, and guide RNAs with <20 nucleotides of complementarity to their targets are generally inactive (6,8). These observations suggest that Cas13 effectors may have higher specificity than RNAi-based approaches. However, unlike Argonaute proteins that have their nuclease domain located immediately next to where the target RNA is bound (20), the HEPN (Higher Eukaryotic and Prokaryotic Nucleotide-binding) nuclease domains of Cas13 effectors are located away from the guide RNA:target RNA binding pocket (21–23). In fact, the HEPN domains of activated (target bound) Cas13 effectors can be exposed on the surface of the complex (21–24), thereby potentially allowing RNAs in the immediate vicinity to be indiscriminately cleaved. Such collateral cleavage events of non-target RNAs have been observed in bacteria (25) and *in vitro* (9,26), where they have been exploited to enable the development of viral RNA detection tools (27,28). There is nonetheless conflicting evidence as to whether Cas13 effectors do (29–31) or do not (6–8,32–36) have significant off-target effects (via collateral cleavage events) when expressed in eukaryotic cells. This debate has made it unclear how phenotypes observed with Cas13 should be interpreted and whether Cas13 effectors indeed have biotechnological potential for use as specific RNA knockdown tools in eukaryotic cells.

We thus aimed to determine the efficiency and specificity of several prominent Cas13 effectors, including RxCas13d (also known as CasRx or RfxCas13) (8) and PspCas13b (7), using simple co-transfection assays in *Drosophila* and human cells. We find that the off-target effects of RxCas13d can be as strong as the level of on-target knockdown observed in cells. The extent of off-target effects is positively correlated with the target RNA expression levels, and collateral damage can be observed even after reducing RxCas13d/guide RNA levels. In contrast, we find that PspCas13b shows improved specificity and, unlike RxCas13d, can be used to deplete a *Drosophila* circular RNA without affecting the expression of the associated linear RNA. Notably, we find that the extent of Cas13 off-target effects can differ between RNA targets and between human cell lines, which helps explain why there have been conflicting reports regarding Cas13 specificity. In total, our results underscore the need for caution when designing and interpreting Cas13 experiments, while also suggesting that certain Cas13 effectors, such as PspCas13b, may be more appropriate for use in cells.

## MATERIALS AND METHODS

### Cell culture

*Drosophila* DL1 and S2 cells were cultured at 25°C in Schneider’s *Drosophila* medium (Thermo Fisher Scientific 21720024), supplemented with 10% (v/v) fetal bovine serum (HyClone SH30396.03), 1% (v/v) penicillin-streptomycin (Thermo Fisher Scientific 15140122), and 1% (v/v) L-glutamine (Thermo Fisher Scientific 35050061).

HeLa and HEK293T cells were cultured at 37°C and 5% CO_2_ in Dulbecco’s modified Eagle’s medium (DMEM) containing high glucose (Thermo Fisher Scientific 11995065) supplemented with 10% (v/v) fetal bovine serum (HyClone SH30396.03) and 1% (v/v) penicillin-streptomycin (Thermo Fisher Scientific 15140122).

### *Drosophila* expression plasmids, transfections, and RNA isolation

To generate *Drosophila* plasmids that co-express a Cas13 effector along with its associated guide RNA, site directed mutagenesis was first used to remove the existing BsmBI site from the previously described pUb 3xFLAG MCS plasmid backbone (37). A control guide RNA sequence containing BsmBI sites was then inserted downstream of the *Drosophila* snRNA:U6:96Ab promoter using the NdeI restriction site. HA-tagged versions of RxCas13d (8), PspCas13b (7), or LwaCas13a (6) (or their associated catalytic dead versions) were then cloned downstream of the constitutive *Drosophila* Ubi-p63e promoter. Lastly, each guide RNA of interest was cloned into the BsmBI sites. Full details of the cloning, including full plasmid sequences, are described in detail in the **Supplementary Material**. The sequences of all guide RNAs used in *Drosophila* are listed in **Supplementary Table S1**.

*Drosophila* reporter plasmids expressing eGFP (Hy_pMT eGFP SV40; Addgene #69911), *Laccase2* Exons 1-3 (Hy_pMT Laccase2 Exons 1-3; Addgene #91799), and nLuc (Hy_pMtnA nLuc SV40; Addgene #132654) under the control of the inducible Metallothionein A promoter were described previously (38-40). Reporter plasmids expressing mCherry (Hy_pMtnA mCherry SV40; Addgene #176302) and FFLuc (Hy_pMtnA FFLuc SV40; Addgene #176299) under the control of the Metallothionein A promoter were generated by replacing the eGFP ORF of Hy_pMT eGFP SV40 with the indicated ORF. Full details of the cloning, including full plasmid sequences, are described in detail in the **Supplementary Material**. *Drosophila* reporter plasmids expressing eGFP (Hy_pUbi-p63e eGFP SV40; Addgene #132650) under the control of the Ubi-p63e promoter was described previously (40). The associated mCherry plasmid (Hy_pUbi-p63e mCherry SV40; Addgene #176300) was generated by replacing the eGFP ORF with mCherry.

DL1 cells were seeded in 12-well plates (5 × 10^5^ cells per well) in complete Schneider’s *Drosophila* medium and cultured overnight. On the following day, 500 ng of plasmid DNA was transfected into each well using Effectene (4 μL of enhancer and 5 μL of Effectene reagent; Qiagen 301427). Unless otherwise noted, 50 ng of Cas13 expression plasmid was transfected along with 225 ng of each of the on-/off-target expression plasmids. pUb-3xFLAG-MCS (No BsmBI) was used as the empty vector plasmid. Total RNA was isolated ~40 hr later using TRIzol (Thermo Fisher Scientific 15596018) according to the manufacturer’s instructions. When examining reporters driven by the Metallothionein A promoter, a final concentration of 500 μM copper sulfate (Fisher BioReagents BP346-500) was added to cells for the last 14 hr prior to RNA isolation.

S2 cells were seeded in 6-well plates (4 × 10^6^ cells per well) in complete Schneider’s *Drosophila* medium and cultured overnight. On the following day, 2000 ng of plasmid DNA was transfected into each well using Effectene (16 μL of enhancer and 30 μL of Effectene reagent; Qiagen 301427). As above, total RNA was isolated ~40 hr later using TRIzol (Thermo Fisher Scientific 15596018) according to the manufacturer’s instructions.

### Human expression plasmids, transfections, and RNA isolation

The RxCas13d-2A-eGFP expression plasmid (pXR001; Addgene #109049) and the corresponding guide RNA expression vector (pXR003; Addgene #109053) were described previously (8). The PspCas13b-2A-eGFP expression vector was a gift from Eric Wang (University of Florida) and made by replacing the RxCas13d ORF in pXR001 with PspCas13b. The corresponding PspCas13b guide RNA expression vector was described previously (pC0043; Addgene #103854) (7). All guide RNAs were cloned into the BbsI sites and their sequences are listed in **Supplementary Table S2**.

pcDNA3.1(+)/AUG-nLuc-3XFLAG reporter (Addgene #127299) was previously described (41) and the FFLuc expression plasmid (pGL4.13) was obtained from Promega. The pcDNA3.1(+) mCherry plasmid (Addgene #176301) was generated by replacing the eGFP ORF in pcDNA3.1(+) eGFP (Addgene #109020) with mCherry. Humanized Renilla luciferase reporter (p CIneo-RL; Addgene #115366) was described previously (42).

HeLa cells (3 × 10^5^ cells per well) or HEK293T cells (5.5 × 10^5^ cells per well) were seeded in 6-well plates in complete DMEM medium and cultured overnight. On the following day, 1000 ng of plasmid DNA was transfected into each well using Lipofectamine 2000 (Thermo Fisher Scientific 11668019) according to the manufacturer’s instructions. Unless otherwise noted, 300 ng of Cas13 expression plasmid and 200 ng of guide RNA expression plasmid were transfected along with 250 ng of each of the on-/off-target expression plasmids. pBEVY-L was used as the empty vector plasmid (43). Total RNA was isolated 40-48 hr later, as indicated, using TRIzol (Thermo Fisher Scientific 15596018) according to the manufacturer’s instructions.

### Northern blotting

Northern blots using NorthernMax reagents (Thermo Fisher Scientific) and oligonucleotide probes were performed as previously described (44). Oligonucleotide probe sequences are provided in **Supplementary Table S3**. Blots were viewed with the Typhoon 9500 scanner (GE Healthcare) and quantified using ImageQuant (GE Healthcare). Representative blots are shown.

### Western blotting

Cells were lysed 40 hr after transfection using RIPA buffer (150 mM NaCl, 1% Triton X-100, 50 mM Tris pH 7.5, 0.1% SDS, 0.5% sodium deoxycholate, and protease inhibitors [Roche 11836170001]) for 20 min on ice. Lysates were then spun at 12,000 × *g* for 15 min and the supernatants were moved to new tubes. Samples were resolved on NuPAGE 4-12% Bis-Tris gels (Thermo Fisher Scientific NP0323) and transferred to PVDF membranes (Bio-Rad 1620177). Membranes were blocked with 5% nonfat milk for 1 hr before incubation in primary antibody (diluted in 1x TBST) overnight at 4°C. Membranes were then washed with 1x TBST (3 x 5 min) followed by incubation in secondary antibody at room temperature for 1 hr. The following antibodies were used: rabbit anti-HA (1:1,000, Abcam ab9110), mouse anti-α-tubulin (1:10,000, Sigma T6074), rabbit anti-GAPDH (1:10,000, Proteintech 10494-1-AP), sheep anti-mouse IgG/HRP (1:10,000, Cytiva NA931), and donkey anti-rabbit IgG/HRP (1:10,000, Cytiva NA934). Membranes were processed using SuperSignal West Pico PLUS Chemiluminescent Substrate (Thermo Fisher Scientific 34580) according to the manufacturer’s protocol.

### RT-qPCR

9 μg of total RNA was digested with TURBO DNase (Thermo Fisher Scientific AM2238) in a 50 μL reaction following the manufacturer’s protocol. 1 μg of the digested RNA was then reverse transcribed to cDNA in a 20 μL reaction using SuperScript III (Thermo Fisher Scientific 18080051) with random hexamers following the manufacturer’s protocol. cDNA was diluted 10-fold with DEPC-treated water and RT-qPCR was performed using Power SYBR Green PCR Master Mix (Thermo Fisher Scientific 4368708). RT-qPCR reactions were performed in 15 μL reactions that contained 1.5 μL of diluted cDNA, 7.5 μL 2x Power SYBR Green PCR Master Mix, and 6 μL 1.5 μM gene-specific primer pairs. Primer sequences are provided in **Supplementary Table S4**.

Using the LightCycler 96 Real-Time PCR System (Roche), the following cycling conditions were used: 95°C for 10 min, 40 amplification cycles of 95°C for 15 s followed by 60°C for 1 min, and a final melting cycle of 95°C for 10 s, 65°C for 1 min, and 97°C for 1 s. Relative transcript levels were calculated using the 2^−ΔΔCT^ method. RT-qPCR reactions were performed using three independent biological replicates, with each replicate having two technical replicates.

### Quantification and statistical analysis

For Northern blots and RT-qPCR data, statistical significance for comparisons of means was assessed by one-way ANOVA. Statistical details and error bars are defined in each figure legend.

## RESULTS

### Strong off-target effects can be observed when RxCas13d is used to deplete an RNA of interest in *Drosophila* cells

To characterize the efficiency and specificity of different classes of Cas13 effectors in *Drosophila* cells, HA-tagged versions of RxCas13d (8), PspCas13b (7), or LwaCas13a (6) were inserted into expression plasmids downstream of the constitutively active Ubi-p63e promoter **(Figure 1A)**. A guide RNA expression cassette driven by the U6 promoter was also inserted, thereby allowing both the Cas13 effector protein **(Supplementary Figure S1A)** and its associated guide RNA **(Supplementary Figure S1B, C)** to be expressed from a single plasmid. We then took advantage of simple co-transfection assays using fluorescent protein reporter genes to characterize the on- and off-target effects of each Cas13 effector, starting with RxCas13d (from *Ruminococcus flavefaciens*) because it is among the smallest and most commonly used Cas13 effectors (8,29,33,34,36,45–48). As diagrammed in **Figure 1A**, *Drosophila* DL1 cells were transiently transfected with the RxCas13d/guide RNA (24-nucleotide spacer length) expression plasmid along with reporter gene plasmids that express eGFP and mCherry from the copper-inducible Metallothionein A (MtnA) promoter. 24 hr after transfection, copper sulfate was added to induce transcription of eGFP and mCherry. Total RNA was subsequently isolated after an additional 14 hr. Northern blots revealed, as expected, that expression of RxCas13d and a random guide RNA sequence had no effect on expression of the fluorescent reporter genes (**Figure 1B)**. Guide RNAs complementary to eGFP **(Figure 1B, left)** or mCherry **(Figure 1B, right)** resulted in 60-75% depletion of the target transcript, indicative of strong on-target knockdown by RxCas13d. However, for all of the eGFP and mCherry guide RNAs tested, we notably detected equally strong depletion (60-75%) of the non-target reporter mRNA as well as the mRNA encoding RxCas13d itself **(Figure 1B)**.

**Figure 1.**
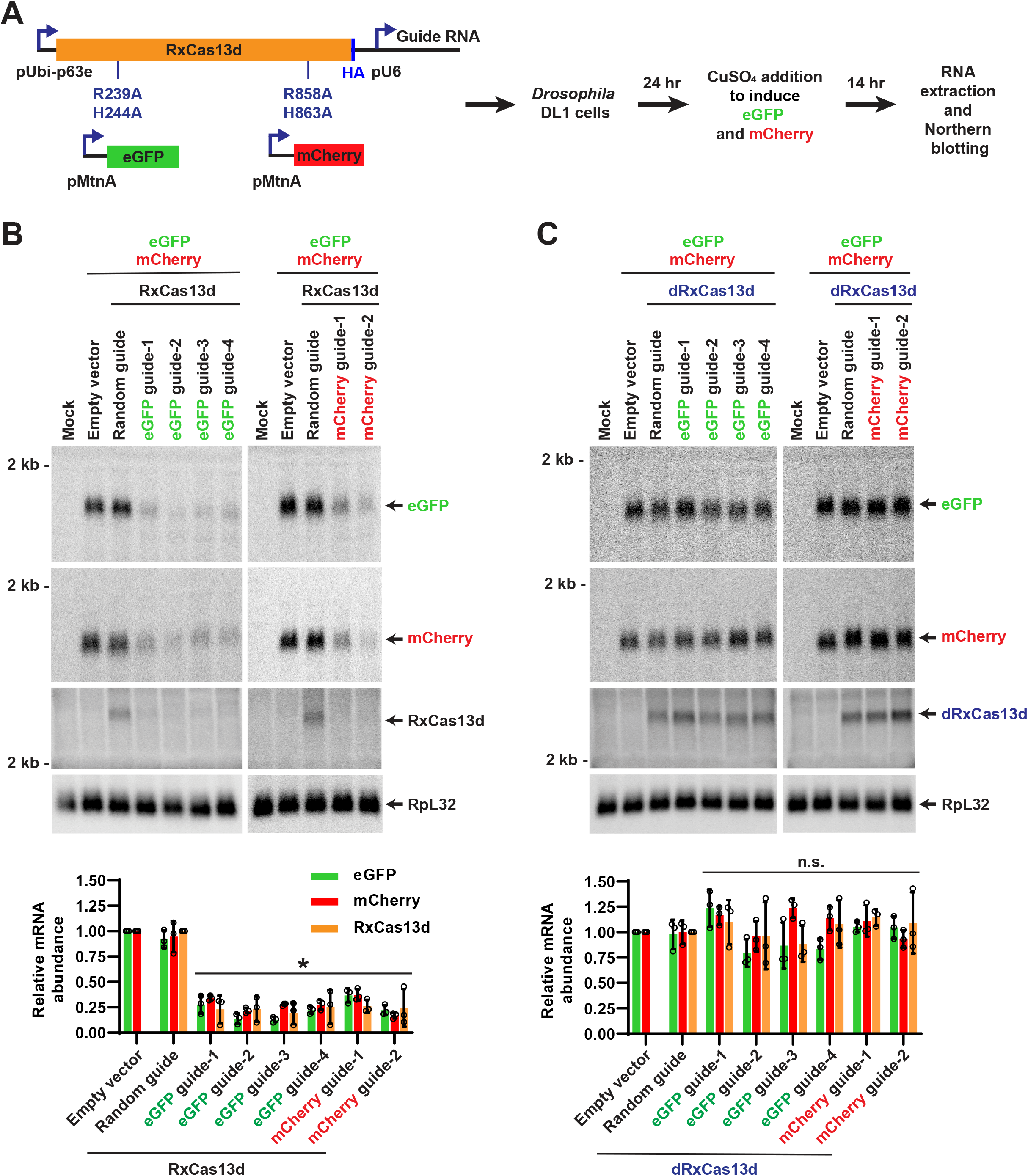
Co-transfection assays revealed that RxCas13d has significant off-target effects in *Drosophila* cells. **(A)** *Drosophila* DL1 cells were co-transfected with (i) 50 ng of plasmid that constitutively expresses a guide RNA from the U6 promoter as well as HA-tagged catalytically active or dead (R239A, H244A, R858A, and H863A mutations) RxCas13d from the Ubi-p63e promoter, (ii) 225 ng of plasmid that expresses eGFP from the copper-inducible MtnA promoter, and (iii) 225 ng of plasmid that expresses mCherry from the MtnA promoter. 24 hr after transfection, CuSO_4_ was added and total RNA was isolated after an additional 14 hr. Northern blots were then performed. **(B)** Plasmids expressing active RxCas13d and a guide RNA complementary to eGFP (left) or mCherry (right) were employed in the co-transfection assay. Representative Northern blots (20 μg of total RNA/lane) are shown. ImageQuant was used to quantify the relative expression levels of eGFP, mCherry, and RxCas13d mRNAs from three independent experiments. eGFP and mCherry mRNA expression was normalized to the empty vector samples, while RxCas13d mRNA expression was normalized to the random guide RNA samples. RpL32 mRNA served as an endogenous loading control. Data are shown as mean ± SD. For statistical comparisons, data were compared to the random guide RNA samples. (*) *P* < 0.05. **(C)** Same as **B** except that plasmids expressing catalytic dead RxCas13d (dRxCas13d) were used. n.s., not significant.

Given that no off-target effects were observed with the random guide RNA **(Figure 1B)**, these results suggested that recognition of the target mRNA by the RxCas13d/guide RNA complex may trigger significant non-specific degradation of RNAs in cells. Indeed, collateral degradation of bystander RNAs has been observed with a number of Cas13 effectors *in vitro* (9,26-28), but several recent studies have suggested Cas13 family members have limited or no off-target effects in cells (6-8,32-36). Because *Drosophila* DL1 cells have low transfection efficiency (~10%), it is difficult to see degradation of endogenous transcripts (e.g. RpL32, **Figure 1B**) using our co-transfection assay. Nonetheless, the fluorescent reporter genes make clear that the off-target effects of RxCas13d can be as strong as on-target knockdown in cells.

### Once activated, the nuclease activity of RxCas13d can be inherently non-specific in *Drosophila* cells

We aimed to explore the underlying cause of the RxCas13d off-target effects and determine if cellular conditions could be identified in which bystander RNA degradation is minimized. First, the co-transfection assays were repeated using catalytic dead RxCas13d (denoted dRxCas13d) that harbors quadruple mutations (R239A/H244A/R858A/H863A) in the HEPN (Higher Eukaryotic and Prokaryotic Nucleotide-binding) nuclease domains **(Figure 1A)**. These mutations abolish the nuclease activity of RxCas13d but do not affect its RNA binding activity (8,21). No significant changes in eGFP or mCherry mRNA levels were observed when dRxCas13d was employed **(Figure 1C)**. This indicates that the HEPN nuclease domains are critical for both the on- and off-target effects of RxCas13d in cells.

We next examined whether the off-target effects are dependent on the presence of the target RNA **(Supplementary Figure S2)**. Similar to the assay diagrammed in **Figure 1A**, DL1 cells were co-transfected with a plasmid expressing RxCas13d/a guide RNA complementary to eGFP and a plasmid expressing eGFP (target RNA) driven by the inducible MtnA promoter. In this case, the co-transfected mCherry (off-target RNA) reporter was driven by the Ubi-p63e promoter, thereby allowing mCherry to be constitutively expressed. Off-target depletion of mCherry by RxCas13d was only observed when expression of the eGFP target mRNA had been induced **(Supplementary Figure S2)**. Collectively, these data confirm that the off-target effects of RxCas13d in cells are due to the RxCas13d/guide RNA complex recognizing its target mRNA and then activating the HEPN nuclease activity **(Figure 2A)**.

**Figure 2.**
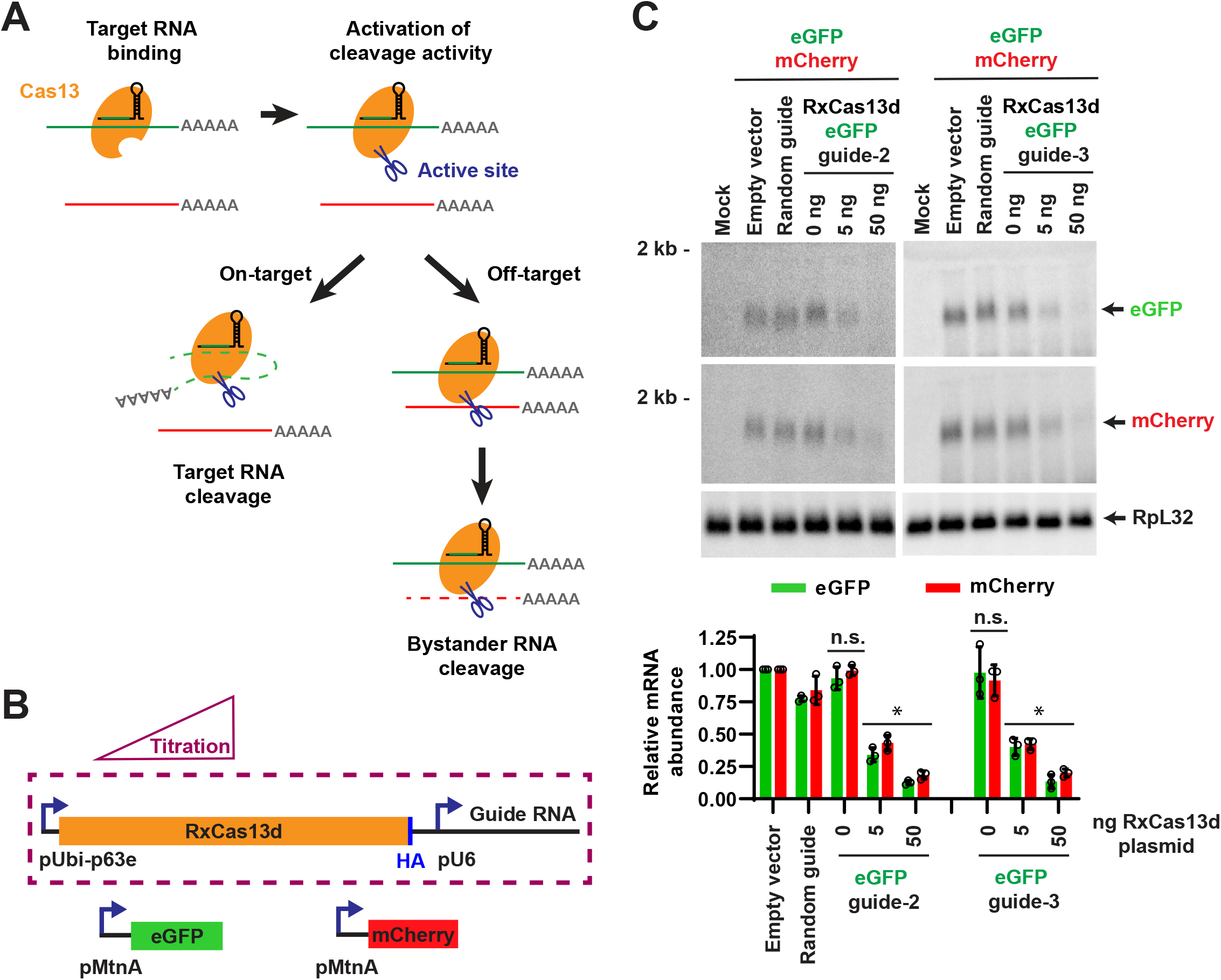
Reducing the expression level of RxCas13d does not diminish the off-target effects. **(A)** Proposed model of Cas13 effector activation. Base pairing between the guide RNA and its target mRNA (green) leads to a conformational change in Cas13 (orange) that activates the endonuclease activity on the surface of the protein. This can result in cleavage/degradation of the target RNA (left) but also cleavage of bystander RNAs (red), resulting in off-target effects (right). This model predicts that increased levels of target mRNA should result in increased off-target effects due to increased numbers of active Cas13 protein in cells. **(B)** *Drosophila* DL1 cells were transfected with decreasing amounts of RxCas13d/guide RNA expression plasmid, while the amounts of the eGFP (225 ng) and mCherry (225 ng) plasmids transfected were kept constant. Empty vector (pUb-3xFLAG MCS (No BsmBI) plasmid) was added as needed so that 500 ng DNA was transfected in all samples. 24 hr after transfection, CuSO_4_ was added and total RNA was isolated after an additional 14 hr. **(C)** Northern blots were used to quantify the relative expression levels of eGFP and mCherry mRNA. Data are shown as mean ± SD, N=3. For statistical comparisons, data were compared to the random guide RNA samples. (*) *P* < 0.05. n.s., not significant.

To test whether RxCas13d generally has significant bystander effects in *Drosophila* cells, we systematically altered each of the components of the co-transfection assay. Neither extending the RxCas13d guide RNA spacer length from 24 to 30 nucleotides **(Supplementary Figure S3A)** nor changing the promoter **(Supplementary Figure S3B)** or the open reading frame **(Supplementary Figure S3C)** of the on-/off-target reporter mRNAs diminished the off-target effects observed with RxCas13d. In all cases, the extent of off-target effects remained as strong as the level of on-target knockdown observed **(Supplementary Figure S3A-C)**. Similar results were observed in *Drosophila* S2 cells **(Supplementary Figure S3D)**. It thus appears that the nuclease activity of activated (target bound) RxCas13d may be inherently non-specific in *Drosophila* cells **(Figure 2A)**.

### Off-target effects of RxCas13d are positively correlated with the expression level of the target mRNA

Although the nuclease activity of activated RxCas13d may lack strong specificity, we reasoned that limiting the amounts of RxCas13d/guide RNA and/or the activating target mRNA in cells may maximize on-target effects and diminish off-target RNA degradation. To first test the effect of reducing RxCas13d/guide RNA levels, the amount of RxCas13d/eGFP guide RNA plasmid transfected was reduced by 10-fold (to 5 ng from 50 ng, which was the amount transfected in **Figure 1B**), while keeping the amounts of eGFP (225 ng) and mCherry (225 ng) expression plasmids transfected constant **(Figure 2B)**. Less efficient on-target eGFP mRNA depletion was observed under these conditions **(Figure 2C)**, confirming that RxCas13d/eGFP guide RNA were at limiting levels. However, the degree of off-target degradation of mCherry mRNA still mirrored the efficiency of on-target knockdown **(Figure 2C)**. We thus conclude that reducing the level of RxCas13d/guide RNA was insufficient to enable specific on-target knockdown in the co-transfection assay. This is perhaps because the target RNA is expressed at levels high enough that the RxCas13d/guide RNA complexes are constantly bound by target RNA, thereby activating the non-specific RxCas13d nuclease activity.

To directly test the importance of target RNA levels, we decreased the expression of the eGFP target mRNA and examined on-/off-target effects **(Figure 3)**. Previously, in **Figures 1B and 2C**, 225 ng of target mRNA expression plasmid was used, so the amount of eGFP plasmid used here was progressively decreased to as low as 2 ng while keeping the amounts of RxCas13d/guide RNA (50 ng) and mCherry (off-target) expression plasmids (225 ng) constant **(Figure 3A)**. RT-qPCR confirmed that reducing the amount of eGFP plasmid transfected resulted in the expected gradient of eGFP mRNA levels **(Supplementary Figure S4A)** with no effect on mCherry mRNA levels **(Supplementary Figure S4B)**. Compared to a control random guide RNA, guide RNAs complementary to eGFP enabled RxCas13d to deplete eGFP mRNA levels by >75% regardless of the amount of eGFP plasmid transfected **(Figure 3B, Supplementary Figure S4C)**. At the highest eGFP expression levels (50 ng eGFP plasmid transfected), the mCherry and RxCas13d mRNAs were also depleted by RxCas13d by ~75%, thereby mirroring the degree of on-target knockdown **(Figure 3C)**. In contrast, as the eGFP mRNA levels were progressively decreased, the amounts of off-target effects also decreased and ultimately went away. For example, when 2 ng of eGFP plasmid was transfected, eGFP guide-2 resulted in >75% depletion of eGFP mRNA **(Figure 3B, Supplementary Figure S4C)** but no significant change in mCherry or RxCas13d mRNA levels **(Figure 3C)**. This suggests that when RxCas13d is used to target lowly expressed RNAs, off-target effects may be minimal as the RxCas13d/guide RNA complex is rarely in an activated state. Furthermore, this observation regarding target RNA abundance likely helps explain in part why some groups have suggested RxCas13d has good specificity in cells (8,33–36), while others (including earlier in this manuscript) have come to the opposite conclusion (29,31).

**Figure 3.**
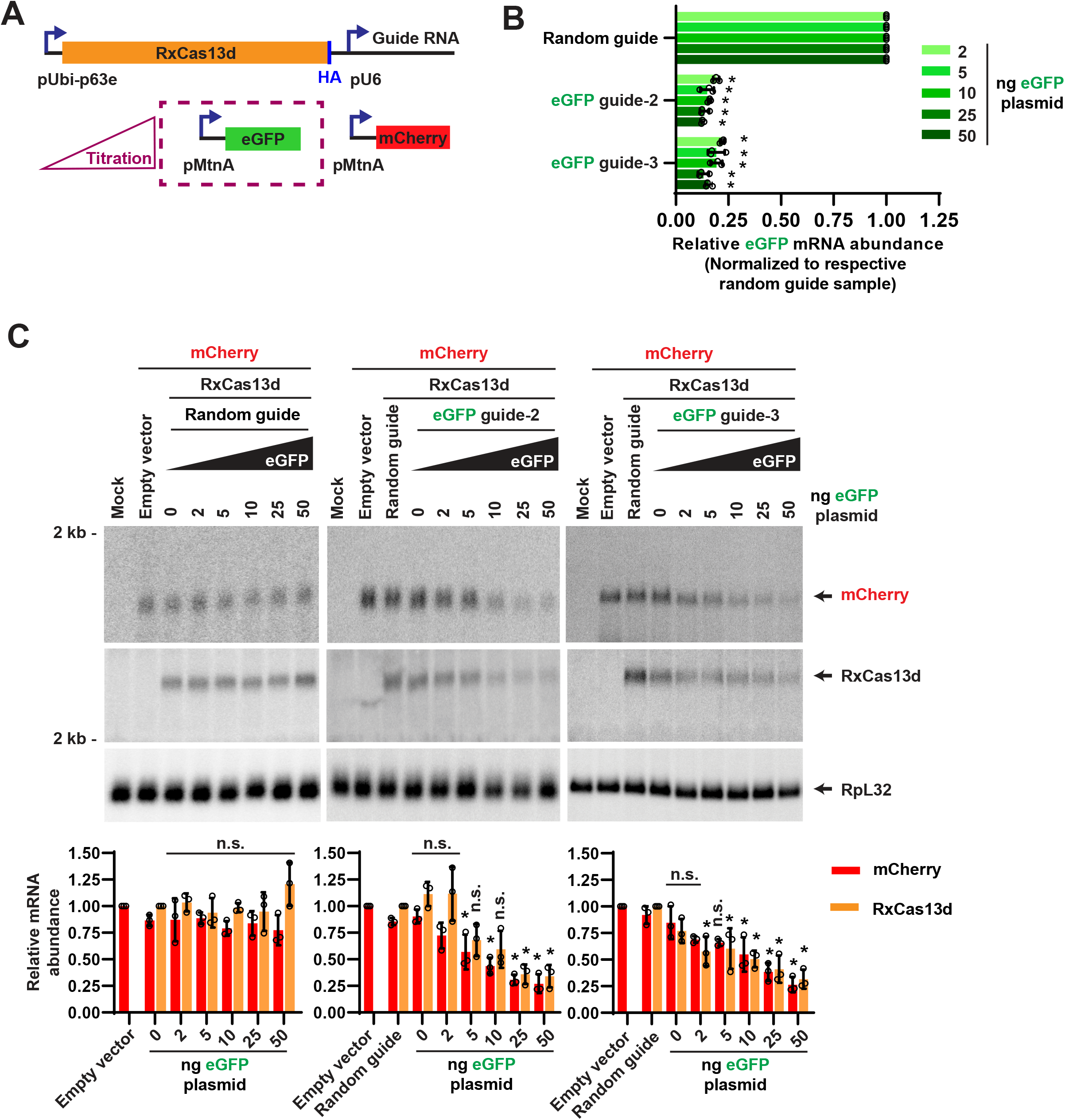
A positive correlation was observed between target mRNA level and RxCas13d off-target effects. **(A)** *Drosophila* DL1 cells were co-transfected with a constant amount of RxCas13d/guide RNA (50 ng) and mCherry (225 ng) expression plasmids, but variable amounts of eGFP expression plasmid (2, 5, 10, 25, or 50 ng). Empty vector (pUb-3xFLAG MCS (No BsmBI) plasmid) was added as needed so that 500 ng DNA was transfected in all samples. 24 hr after transfection, CuSO_4_ was added and total RNA was isolated after an additional 14 hr. RNA expression levels were then analyzed by RT-qPCR **(B)** or Northern blotting **(C)**. **(B)** RT-qPCR was used to quantify the expression of eGFP mRNA in cells transfected with the RxCas13d plasmid expressing a random guide RNA or a guide RNA complementary to eGFP. For each amount of eGFP plasmid transfected, the relative abundance of eGFP mRNA was normalized to the respective random guide RNA samples. Data are shown as mean ± SD, N=3. (*) *P* < 0.05. **(C)**Northern blots were used to quantify the relative expression levels of mCherry and RxCas13d mRNAs. Data are shown as mean ± SD, N=3. For statistical comparisons, data were compared to the random guide RNA samples. (*) *P* < 0.05. n.s., not significant. A complete table of P-values for all comparisons is provided in **Supplementary Table S5**.

### PspCas13b has significantly improved specificity in *Drosophila* cells compared to RxCas13d

The potential for significant off-target effects with RxCas13d severely limits its ability to catalyze specific depletion of RNAs of interest in *Drosophila* cells. We thus next examined other commonly used Cas13 effectors, especially those in other Cas13 subtypes **(Supplementary Figure S1A-B)**. Cas13 effectors all possess nucleic acid recognition and RNA cleavage activities, but they are divided into 6 subtypes that have low sequence similarity beyond the active sites for the two HEPN domains (10). In fact, Cas13 subtypes vary greatly in size and domain organization, including the locations of the HEPN domains. We thus reasoned that the on-/off-target effects of different Cas13 effectors could be highly divergent in *Drosophila* cells.

LwaCas13a (isolated from *Leptotrichia wadei*) was originally shown in 2017 to be able to elicit knockdown of target RNAs in *E. coli* and human HEK293FT cells, with an efficiency similar to that of short hairpin RNAs (shRNAs) in human cells (6). We found that the LwaCas13a effector protein and its associated guide RNA (30-nt spacer length) are able to be expressed in *Drosophila* DL1 cells **(Supplementary Figure S1A-B)**. However, no depletion of target mRNAs could be observed when LwaCas13a was employed in the same co-transfection assay that had been used in **Figure 1** to characterize RxCas13d **(Supplementary Figure S5)**. This negative result is consistent with a recent report that also failed to observe target mRNA knockdown by LwaCas13a in human cells (34).

We instead obtained much more promising results when PspCas13b (from *Prevotella sp. P5-*125) (7) was used in the co-transfection assays **(Figure 4A)**. Guide RNAs (30-nt spacer length) complementary to eGFP **(Figure 4B, left)** or mCherry **(Figure 4B, right)** resulted in 50-75% depletion of the target transcript, indicative of strong on-target knockdown by PspCas13b. As expected, these on-target effects were lost with catalytic dead PspCas13b (denoted dPspCas13b), which harbors mutations (H133A/H1098A) in the HEPN nuclease domains (7) **(Figure 4C)**. In stark contrast to what we observed with RxCas13d **(Figure 1B)**, all of the eGFP and mCherry guide RNAs tested with PspCas13b resulted in no significant change in expression of the non-target reporter mRNA or the mRNA encoding the Cas13 effector itself **(Figure 4B)**. This suggests that PspCas13b has better specificity than RxCas13d in *Drosophila* cells.

**Figure 4.**
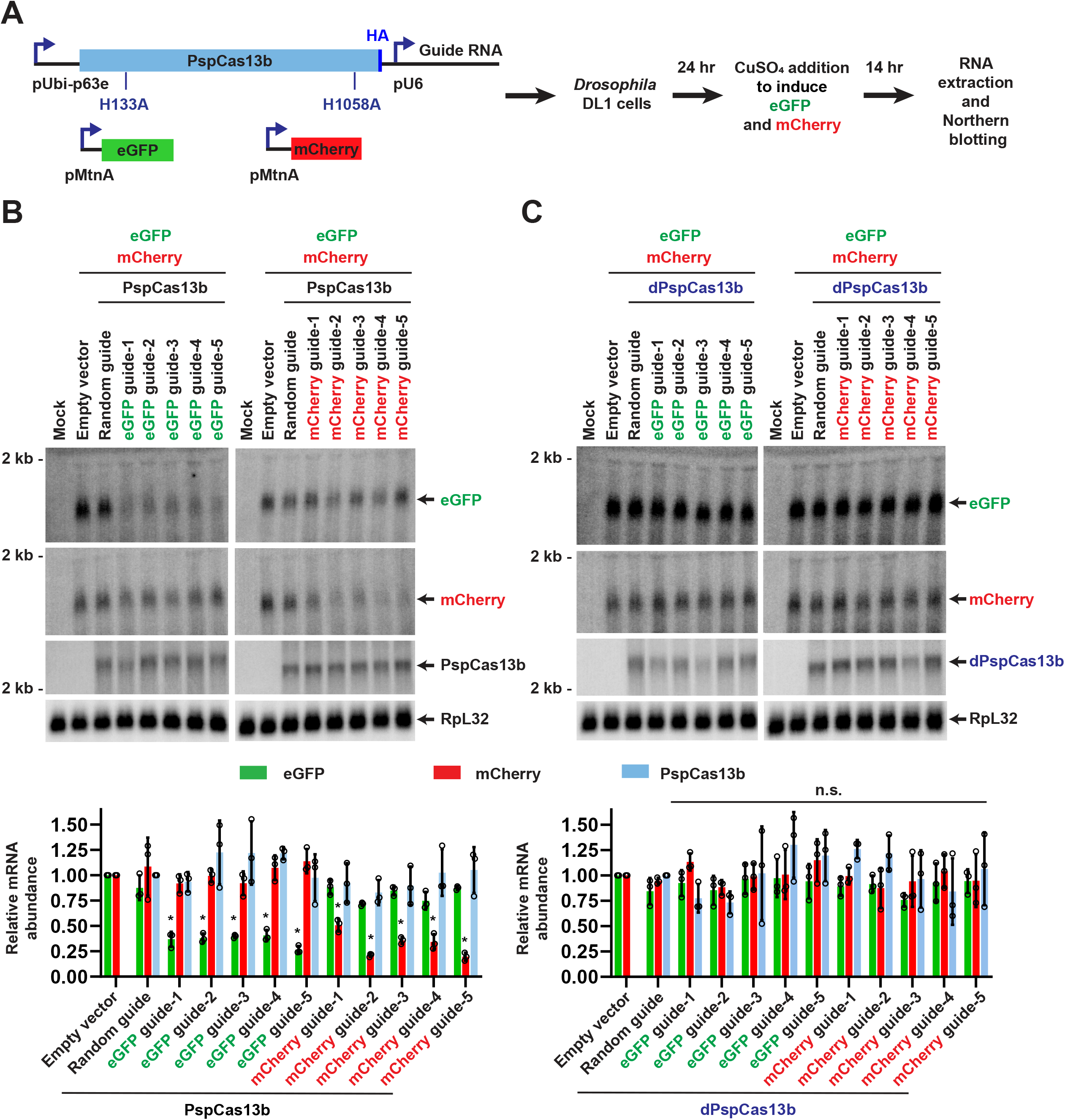
Co-transfection assays revealed PspCas13b has better specificity in *Drosophila* cells. **(A)** *Drosophila* DL1 cells were co-transfected with (i) 50 ng of plasmid that constitutively expresses a guide RNA from the U6 promoter as well as HA-tagged catalytically active or dead (H133A and H1058A mutations) PspCas13b from the Ubi-p63e promoter, (ii) 225 ng of plasmid that expresses eGFP from the copper-inducible MtnA promoter, and (iii) 225 ng of plasmid that expresses mCherry from the MtnA promoter. 24 hr after transfection, CuSO_4_ was added and total RNA was isolated after an additional 14 hr. Northern blots were then performed. **(B)** Plasmids expressing active PspCas13b and a guide RNA complementary to eGFP (left) or mCherry (right) were employed in the co-transfection assay. Representative Northern blots (20 μg of total RNA/lane) are shown. ImageQuant was used to quantify the relative expression levels of eGFP, mCherry, and PspCas13b mRNAs from three independent experiments. eGFP and mCherry mRNA expression was normalized to the empty vector samples, while PspCas13b mRNA expression was normalized to the random guide RNA samples. RpL32 mRNA served as an endogenous loading control. Data are shown as mean ± SD. For statistical comparisons, data were compared to the random guide RNA samples. (*) *P* < 0.05. No significant changes in expression of the mRNAs encoding PspCas13b or the off-target fluorescent protein were found. **(C)** Same as **B** except that plasmids expressing catalytic dead PspCas13b (dPspCas13b) were used. n.s., not significant.

To test the robustness of PspCas13b, we systematically altered each of the components of the co-transfection assay in a manner analogous to how RxCas13d was characterized in **Supplementary Figure S3**. The PspCas13b guide RNA spacer length was decreased from 30 to 24 nucleotides **(Supplementary Figure 6A)**, the promoter **(Supplementary Figure S6B)**or open reading frame **(Supplementary Figure S6C)** of the on-/off-target reporter mRNAs was changed, and the assays were performed in *Drosophila* S2 cells rather than DL1 cells **(Supplementary Figure S6D)**. In all cases, we found that PspCas13b was able to catalyze on-target knockdown with no significant change in expression of the off-target reporter.

### PspCas13b, but not RxCas13d, can be used to specifically deplete a circular RNA in *Drosophila* cells

Recent work has revealed that many eukaryotic protein-coding genes can be alternatively spliced to yield linear RNAs as well as covalently closed circular RNAs (49–52) **(Figure 5A, left)**. Because the exonic sequence(s) present in a mature circular RNA are almost always also present in its cognate linear mRNA, approaches for depleting circular RNAs have focused on the backsplicing junction (BSJ) as this is the only sequence unique to the circular RNA. Indeed, two recent reports have suggested that targeting RxCas13d to a circular RNA using a guide RNA complementary to its BSJ can be used to specifically deplete the transcript without affecting cognate linear mRNA levels (34,36). Given our data from the co-transfection assays in *Drosophila* cells, we wanted to revisit this approach and determine whether RxCas13d or PspCas13b could indeed be used to specifically deplete a circular RNA in *Drosophila* cells.

**Figure 5.**
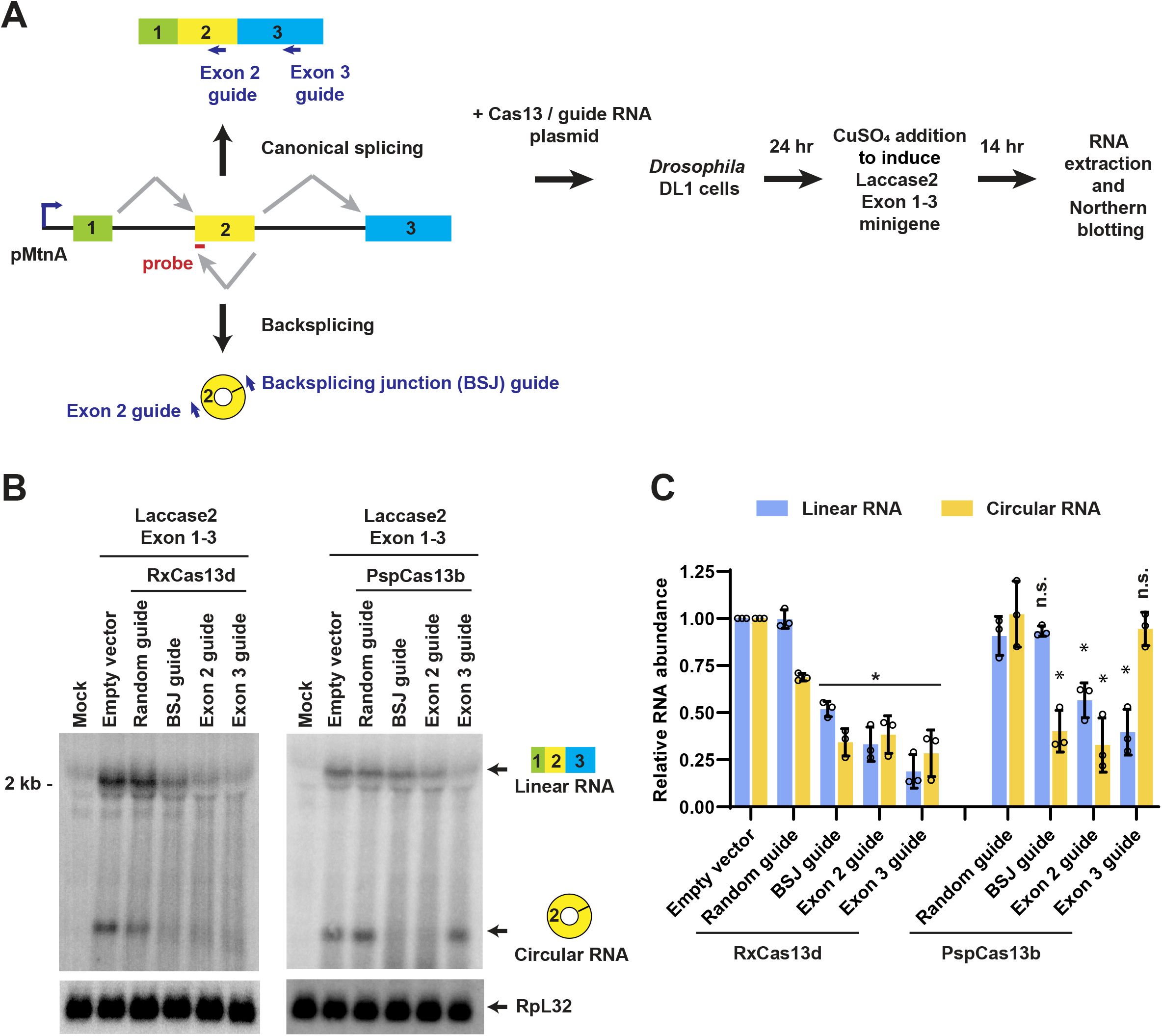
PspCas13b, but not RxCas13d, can be used to specifically deplete a circular RNA in *Drosophila* cells. **(A)** A three-exon *Laccase2* minigene driven by the copper-inducible MtnA promoter can be alternatively spliced to yield a linear mRNA or a circular RNA derived from exon 2. To test the ability of Cas13 effectors to catalyze isoform-specific depletion, guide RNAs were designed that should deplete only the linear RNA (Exon 3 guide), only the circular RNA (BSJ guide), or both the linear and circular RNAs (Exon 2 guide). *Drosophila* DL1 cells were co-transfected with (i) 50 ng of plasmid that constitutively expresses a guide RNA as well as RxCas13d or PspCas13b effector and (ii) 450 ng of the *Laccase2* Exon 1-3 minigene expression plasmid. 24 hr after transfection, CuSO_4_ was added and total RNA was isolated after an additional 14 hr. **(B)** Representative Northern blots (20 μg of total RNA/lane) using an oligonucleotide probe complementary to exon 2 of the *Laccase2* minigene. **(C)** ImageQuant was used to quantify the expression levels of linear and circular RNA normalized to the empty vector samples. RpL32 mRNA served as an endogenous loading control. Data are shown as mean ± SD, N=3. For statistical comparisons, data were compared to the random guide RNA samples. (*) *P* < 0.05.

We focused on the *Drosophila laccase2* (*straw*) gene, which can be backspliced to generate a 490-nt circular RNA from exon 2 that accumulates to high levels *in vivo* (38,53). In prior work, we characterized the *cis*-acting sequences required for *laccase2* backsplicing and generated a copper-inducible three-exon minigene plasmid (Hy_pMT Laccase2 Exons 1-3) that efficiently produces a three-exon linear RNA as well as a circular RNA from exon 2 (38,39) **(Figure 5A, left)**. To test the ability of Cas13 effectors to modulate the output of this minigene, guide RNAs were designed that are complementary to (i) only the circular RNA (BSJ guide), (ii) only the linear RNA (Exon 3 guide), or (iii) both the linear and circular RNAs (Exon 2 guide). Co-transfection assays were then performed in *Drosophila* DL1 cells and Northern blots used to quantify the levels of linear and circular RNA derived from the minigene **(Figure 5A)**. When RxCas13d was employed, the knockdown effects lacked specificity: the BSJ guide RNA (which should be specific for the circular RNA) also significantly depleted the linear mRNA and the Exon 3 guide RNA (which should be specific for the linear RNA) also strongly depleted the circular RNA **(Figures 5B-C, left)**. In contrast, PspCas13b was able to deplete only the transcript of interest **(Figures 5B-C, right)**. The PspCas13b BSJ guide RNA depleted the target circular RNA by 60% +/− 11% while having no significant effect on linear RNA levels. Similarly, the PspCas13b Exon 3 guide RNA depleted the target linear RNA by 60% +/− 12% while having no significant effect on circular RNA levels. These observations suggest that PspCas13b, but not RxCas13d, can be used to specifically knock down individual spliced isoforms, including circular RNAs, in *Drosophila* cells.

### RxCas13d can have strong off-target effects in human cells

Having defined the efficiency and specificity of several Cas13 effectors in *Drosophila* cells, we next aimed to perform similar co-transfection analyses in human cells, focusing first on RxCas13d. HeLa cells were transiently transfected with plasmids that constitutively express (i) HA-tagged RxCas13d (with nuclear localization sequences) followed by a 2A peptide and eGFP (8) **(Supplementary Figure S7A)**, (ii) a guide RNA **(Supplementary Figure S1C)**, (iii) 3xFLAG-tagged firefly luciferase (FFLuc), and (iv) 3xFLAG-tagged nanoLuciferase (nLuc) **(Figure 6A)**. Guide RNAs complementary to FFLuc **(Figure 6B, left)** or nLuc **(Supplementary Figure S7B)** enabled RxCas13d to deplete the target transcripts by 50-75%. Nonetheless, similar to the results in *Drosophila* cells **(Figure 1B)**, activation of RxCas13d in HeLa cells resulted in significant degradation of the non-target reporter mRNA and Cas13-eGFP transcripts **(Figure 6B, Supplementary Figure S7B)**. Northern blots notably revealed no change in expression of endogenous GAPDH transcripts in these experiments. However, we noted that significantly less total RNA could be isolated from the RxCas13d co-transfection assays when the FFLuc **(Figure 6C)** or nLuc **(Supplementary Figure S7C)** guide RNAs were used. This suggests that the presence of active RxCas13d may cause cells to die or be subjected to a growth disadvantage, which is consistent with several recent reports that indicate RxCas13 can be toxic in *Drosophila*, human U87 glioblastoma cells, and mouse ES cells (29,31). We propose that this defect in transfected cells causes the signal from non-transfected cells to dominate the Northern blots, thereby making it difficult to see changes in expression of endogenous transcripts. Regardless, the reporter mRNAs make it clear that bystander cleavage of RNAs can be an issue for RxCas13d in human HeLa cells **(Figure 6B, Supplementary Figure S7B)**. We indeed also observed significant off-target effects when examining the effect of RxCas13d on other combinations of reporter genes, including Renilla luciferase (RLuc)/mCherry **(Supplementary Figure S8A-C)** and nLuc/RLuc **(Supplementary Figure S9A-C)**.

**Figure 6.**
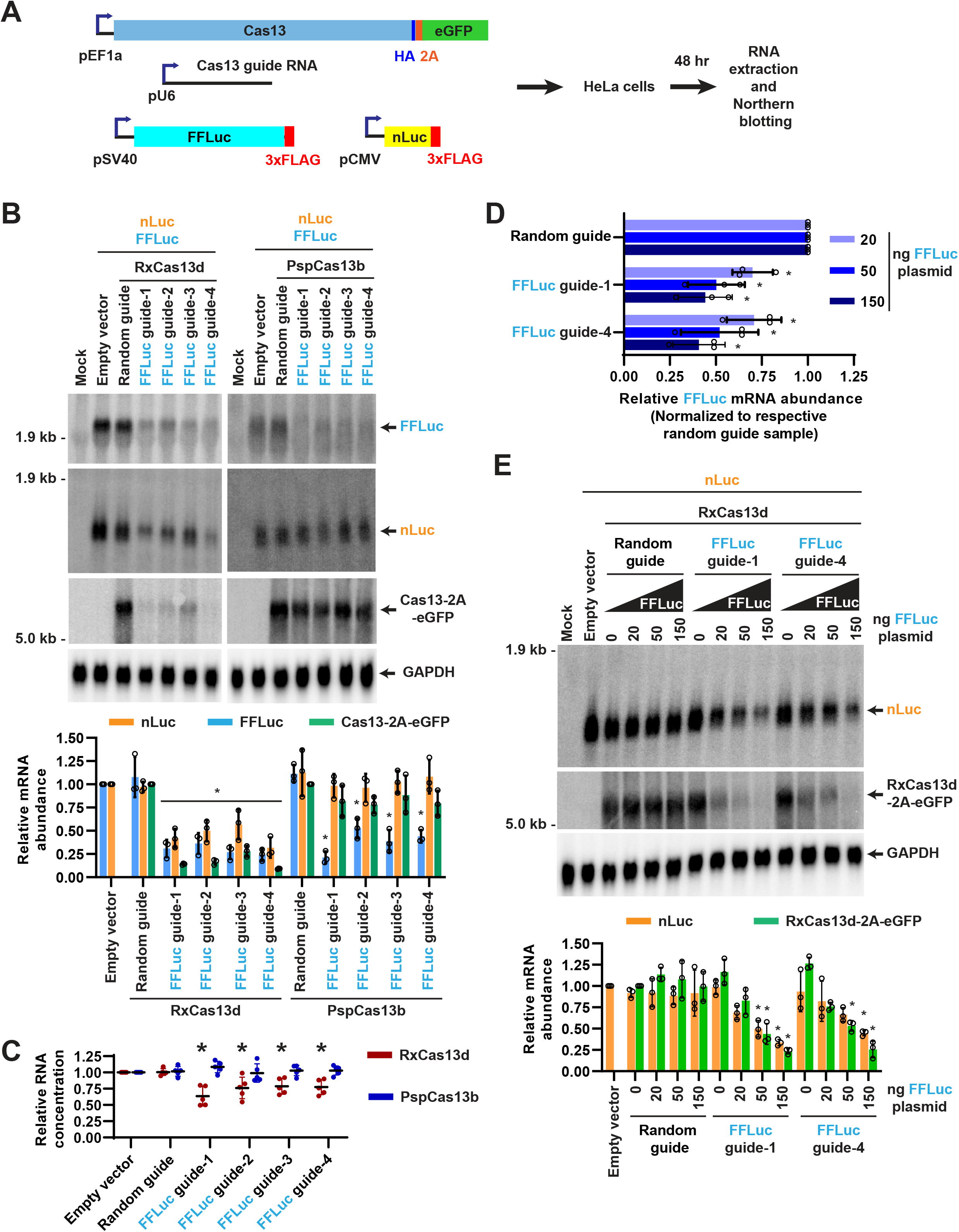
Quantification of on- and off-target effects of RxCas13d and PspCas13b in human HeLa cells. **(A)** HeLa cells were co-transfected with (i) 300 ng of plasmid that constitutively expresses HA-tagged Cas13 protein followed by a 2A peptide and eGFP, (ii) 200 ng of plasmid that expresses a guide RNA, (iii) 250 ng of plasmid that expresses nanoLuciferase (nLuc), and (iv) 250 ng of plasmid that expresses firefly luciferase (FFLuc). 48 hr after transfection, total RNA was isolated and Northern blots performed. **(B-C)** Guide RNAs complementary to FFLuc were employed in the co-transfection assay. **(B)** Representative Northern blots (20 μg of total RNA/lane) are shown. ImageQuant was used to quantify the relative expression level of FFLuc, nLuc, and Cas13-2A-eGFP mRNAs. nLuc and FFLuc mRNA expression was normalized to the empty vector (pBEVY-L) samples, while Cas13-2A-eGFP mRNA expression was normalized to the random guide RNA samples. GAPDH mRNA served as an endogenous loading control. Data are shown as mean ± SD, N=3. For statistical comparisons, data were compared to the random guide RNA samples. (*) *P* < 0.05. **(C)** Relative RNA concentrations obtained from the co-transfection assays when the RxCas13d or PspCas13b expression plasmids were used. Data are normalized to the empty vector samples and shown as mean ± SD, N=5. (*) *P* < 0.05. **(D-E)** HeLa cells were co-transfected with constant amounts of RxCas13d, guide RNA, and nLuc expression plasmids, but variable amounts of FFLuc expression plasmid (20, 50, or 150 ng). **(D)** RT-qPCR was used to quantify depletion of FFLuc mRNA. For each amount of FFLuc plasmid transfected, the relative abundance of FFLuc mRNA was normalized to the respective random guide RNA samples. Data are shown as mean ± SD, N=3. (*) *P* < 0.05. **(E)** Northern blots were used to quantify the relative expression levels of nLuc and RxCas13d-2A-eGFP mRNAs. Data are shown as mean ± SD, N=3. For statistical comparisons, data were compared to the random guide RNA samples. (*) *P* < 0.05. A complete table of P-values for all comparisons in **D-E** is provided in **Supplementary Table S5**.

To determine if the off-target effects of RxCas13d in HeLa cells are diminished when the target mRNA level is reduced, we performed a titration assay analogous to that performed previously in *Drosophila* cells **(Figure 3)**. The amounts of the RxCas13d-2A-eGFP, guide RNA, and nLuc (off-target mRNA) expression plasmids transfected were kept the same as in **Figure 6B**, but the amount of the FFLuc (target mRNA) expression plasmid was reduced from 250 ng to as low as 25 ng **(Figure 6D-E)**. At the highest FFLuc expression level (150 ng FFLuc plasmid transfected), the nLuc and RxCas13d-2A-eGFP transcripts were depleted by 50-75% **(Figure 6E)** which mirrors the amount of on-target nLuc depletion observed **(Figure 6D)**. Meanwhile, when 20 ng FFLuc plasmid was transfected, RxCas13d was able to deplete FFLuc mRNA **(Figure 6D)** with no significant change in nLuc or RxCas13d-2A-eGFP transcript levels **(Figure 6E)**. This suggests RxCas13d may be able to specifically deplete low abundance transcripts in HeLa cells, but these assays need to be interpreted cautiously and one must be aware of the potential for off-target effects.

### The extent of RxCas13d off-target effects differs between human cell lines

Given that several prior reports have indicated that RxCas13d lacks off-target effects in human cells (8,33-36), we next wanted to further explore the underlying reasons for the conflicting observations. A prominent difference between studies is often the cell type examined, so we wanted to perform the same co-transfection experiments side-by-side in HeLa and human embryonic kidney HEK293T cells **(Supplementary Figure S8, S9)**. When guide RNAs complementary to RLuc were employed in HeLa cells, significant depletion of the off-target mCherry (**Supplementary Figure S8B)** or nLuc transcripts **(Supplementary Figure S9B)** was observed and less total RNA could be isolated from the co-transfection assays **(Supplementary Figure S8C, S9C)**. In contrast, when these same experiments were performed in HEK293T cells, guide RNAs complementary to RLuc did not affect expression of the off-target reporter (**Supplementary Figure S8D, S9D)**or the total RNA yield (**Supplementary Figure S8E, S9E)**. This indicates that the extent of off-target effects can differ significantly depending on cell type. Interestingly, the RxCas13d-2A-eGFP transcript was depleted in both HeLa **(Supplementary Figure S8B)**and HEK293T cells **(Supplementary Figure S9B)** when RxCas13d was activated by RLuc guide RNAs. This indicates that there is, in fact, some degree of off-target effects present in both cell types. It remains unclear why some transcripts are more sensitive than others to collateral damage by active RxCas13d, but these results highlight how very different conclusions can be drawn depending on how one measures off-target effects.

### PspCas13b has better specificity than RxCas13d, but still can have some off-target effects in human cells

As PspCas13b behaved significantly better in *Drosophila* cells compared to RxCas13d **(Figures 4–5)**, the co-transfection assays in HeLa cells were repeated using the PspCas13b effector **(Figure 6A-B)**. Guide RNAs complementary to FFLuc **(Figure 6B, right)** or nLuc **(Supplementary Figure S7B)** enabled PspCas13b to deplete the target transcripts by 50-75%, a similar efficiency to that obtained with RxCas13d. The FFLuc guide RNAs resulted in no significant change in nLuc (off-target) or PspCas13b-2A-eGFP mRNA levels **(Figure 6B, right)**, suggesting that PspCas13b may have better specificity than RxCas13d in HeLa cells. Indeed, unlike what was observed for RxCas13d, the FFLuc guide RNAs for PspCas13b had no effect on the amount of total RNA that could be isolated from the co-transfection assays **(Figure 6C)**. Notably, the nLuc guide RNAs did not affect FFLuc (off-target) levels but did, however, reduce PspCas13b-2A-eGFP mRNA levels by ~50% **(Supplementary Figure S7B)**. It remains unclear why the PspCas13b off-target effects were somewhat selective with the nLuc guide RNAs, but we speculate that more bystander cleavage events may be observed with the nLuc vs. the FFLuc guides as the nLuc target mRNA is expressed 2.9 +/− 0.6-fold higher than FFLuc mRNA **(Supplementary Figure S7D)**. Higher mRNA target levels mean that more PspCas13b effectors become activated, thereby giving more opportunity for off-target effects **(Figure 2A)**. Similar selective off-target effects were observed when PspCas13b was used in co-transfection assays using the Renilla luciferase and mCherry reporter genes **(Figure S8)**. We thus conclude that PspCas13b has better specificity than RxCas13d in HeLa cells, but bystander cleavage events still need to be kept in mind.

## DISCUSSION

Recent work has suggested that Cas13 effectors may hold significant promise as tools for knocking down RNAs of interest in eukaryotic cells. There nonetheless have been conflicting reports on the specificity and efficiency of these approaches. On the one hand, there is strong agreement that near perfect complementarity between the guide RNA spacer and its target RNA is required for activating Cas13 nuclease activity (7,19). This makes the process of Cas13 activation highly specific. However, the activated Cas13 HEPN nuclease domains are located distally from the guide RNA:target RNA binding pocket and crystallization/cryo-EM efforts have revealed that, at least for some Cas13 effectors, the catalytic residues can be exposed on the enzyme surface (21–24). This structural arrangement is distinct from that observed in Argonaute or Cas enzymes with high specificity, such as Cas9 that cleaves the target DNA sequence at defined positions within the guide RNA-DNA heteroduplex (3,54). Cas13 effectors instead cleave target RNAs in exposed single-stranded regions outside the guide RNA binding site (26), and this mode of action provides the potential for promiscuous cleavage of RNAs beyond the intended target RNA (so-called collateral damage). Indeed, in the initial report characterizing the activity of a Cas13a effector (C2c2), prominent collateral damage effects were noted in bacteria (25) and similar effects have been noted in human glioblastoma cells (30). It was thus highly notable when two other Cas13 effectors, RxCas13d and PspCas13b, were reported to not have off-target effects (as determined by RNA-seq) when used to deplete endogenous RNAs in human cells (7,8). There is significant evolutionary divergence among Cas13 family members, but these latter results are still surprising considering that Cas13 effectors have been developed as *in vitro* tools for viral detection, including SARS-CoV-2, by exploiting their collateral damage activities (55,56). In this study, we thus aimed to re-examine the specificity of prominent Cas13 effectors, including RxCas13d and PspCas13b, in both *Drosophila* and human cells to determine their applicability for knockdown studies.

Using a series of co-transfection assays, we showed that the off-target effects of RxCas13d can be as strong as the level of on-target knockdown achieved **(Figure 1 and Figure 6)**. In particular, as the expression level of the target RNA was increased, the off-target effects concomitantly increased **(Figure 3)**. We propose this is due to increased numbers of activated (target bound) RxCas13d complexes being present in cells. In human HeLa cells, we were able to achieve good transfection efficiency and further found that the presence of active RxCas13d was associated with reduced RNA yield, suggestive of cellular toxicity **(Figure 6, Supplementary Figure S7-S9)**. These results are consistent with recent *in vivo* work in flies that showed on-target RNA knockdown by RxCas13d can be associated with lethality, even when guide RNAs complementary to non-essential genes were used (29). Similar toxicity has now also been observed when RxCas13d is used in human U87 glioblastoma cells and mouse embryonic stem cells (31). Nonetheless, other reports have suggested there are minimal or no off-targets when RxCas13d is used in zebrafish embryos (33) and human HEK293FT cells (8,34,36).

The reason for these contradictory reports has been unclear, but our work helps explain at least part of the confusion. We now have revealed that the degree of Cas13 off-target effects can differ depending on which cell type is examined. When the exact same reporter genes were transfected into HeLa and HEK293T cells, we found that expression of the off-target reporter gene was affected to a significantly greater degree by RxCas13d in HeLa cells **(Supplementary Figure S8-S9)**. The underlying cause of this cell type specific difference remains to be determined, but it could be due to differences in target RNA expression levels or cell-type specific modification of RxCas13d, the guide RNA, or its binding partners. We further notably found that some transcripts, e.g. the mRNA encoding the Cas13 effector itself, are more sensitive than others to off-target effects, perhaps due to their transcript lengths **(Figure 6, Supplementary Figure S7)**. This means that any conclusion regarding Cas13 specificity needs to be made with caution, especially when examining the expression of a single (or small number) of off-target RNAs.

Compared to RxCas13d, we found that the PspCas13b effector may have better specificity, especially in *Drosophila* cells where it was able to deplete a circular RNA without altering the expression of the associated linear RNA **(Figures 5)**. This may be because the catalytic residues of the PspCas13b HEPN domains are less solvent exposed, but unfortunately no structural details of PspCas13b are yet available to address this point. Regardless, it is highly unlikely that PspCas13b functions without off-target effects in cells, as we were able to observe some collateral damage cleavage events when PspCas13b was used in human cells **(Supplementary Figure S7B, S8B)**. Consistent with these data, expression of PspCas13b has been reported to cause abnormal development of zebrafish (33) and to inhibit mouse cerebellar Purkinje cell dendritic growth (57).

In total, our data lead to the obvious question of whether Cas13 effectors can or should be used in eukaryotic cells to knock down a specific RNA of interest. We conclude that caution is needed and there is unfortunately not yet a straightforward answer. We observed prominent roles for target RNA expression and cell type in dictating the degree of Cas13-mediated off-target effects. When trying to deplete a lowly abundant RNA, it seems like it may be possible to achieve specific knockdown (especially with PspCas13b), but it is critical that care is taken to ensure that a minimal number of activated Cas13 proteins are present in cells. This task becomes increasingly more difficult as target RNA abundance increases. There are nonetheless situations when Cas13 collateral damage activity in eukaryotic cells may be useful, e.g. to kill cancer cells that are expressing a tumor-specific antigen or a mutated oncogene (30) or to impair/kill cells that are infected with a pathogen. In the future, further mechanistic studies will hopefully reveal why Cas13 effectors have inherently different propensities to cleave bystander RNAs as well as how these propensities are affected by cellular conditions. This should aid in the development of novel RNA knockdown approaches with both higher efficiency and specificity.

## ACKNOWLEDGEMENTS

We thank all the members of the Wilusz lab, Junwei Shi (University of Pennsylvania), as well as Chase Kelley and Eric T. Wang (University of Florida) for discussions and advice.

## AUTHOR CONTRIBUTIONS

Y.A. and J.E.W. conceived and designed the project. Y.A. and D.L. performed experiments and analyzed data. Y.A. and J.E.W. wrote the manuscript with input from all authors.

