## Supplementary Material for "CRISPR/Cas13 effectors have differing extents of off-target effects that limit their utility in eukaryotic cells"

#### SUPPLEMENTARY FIGURES

Supplementary Figure S1

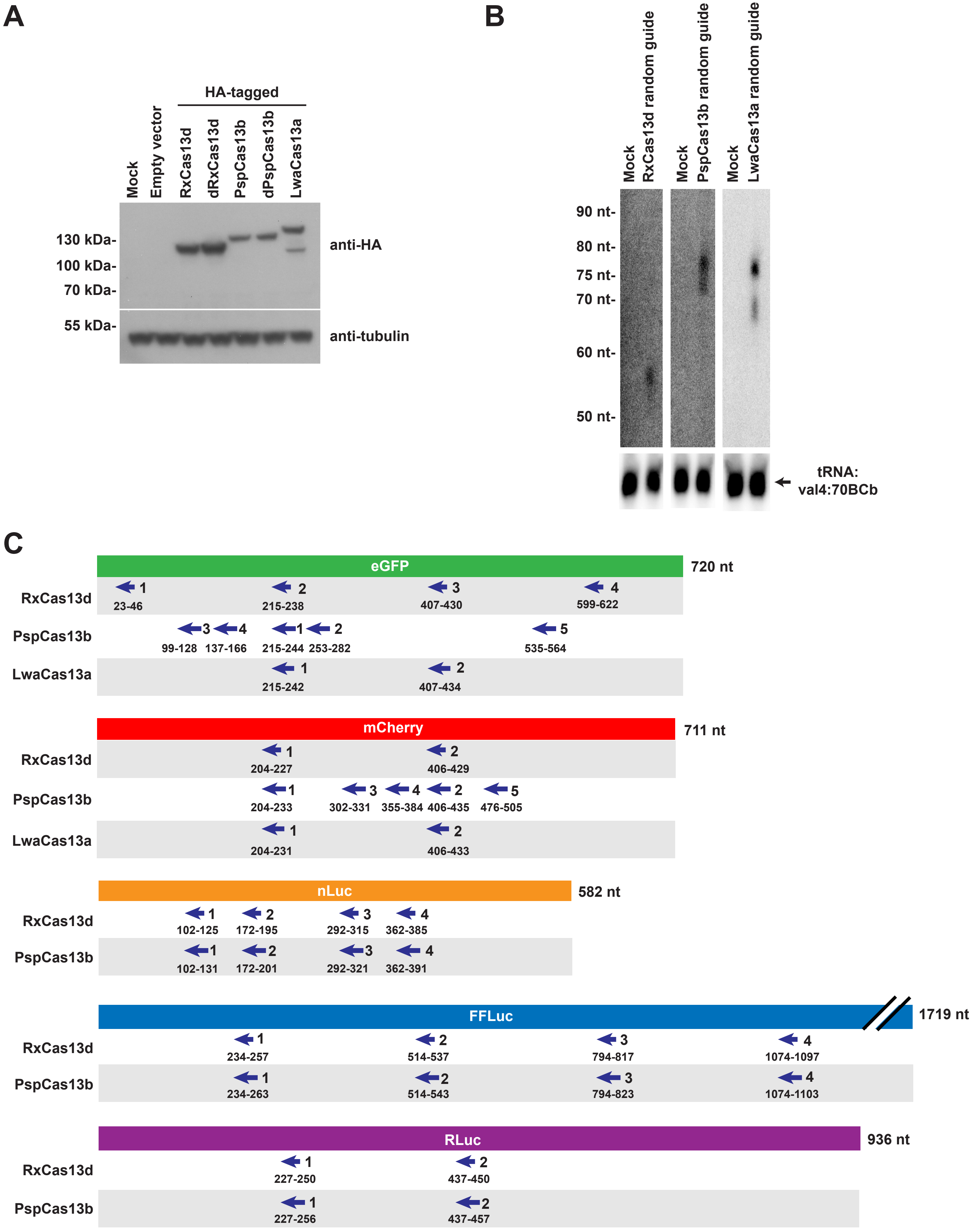

**Supplementary Figure S1.** ***Drosophila*** **Cas13 and guide RNA expression plasmids. (A-B)** Analogous to Figure 1A, *Drosophila* DL1 cells were co-transfected with (i) 50 ng of plasmid that expresses the indicated HA-tagged Cas13 effector as well as a random guide RNA, (ii) 225 ng of plasmid that expresses eGFP from the copper-inducible MtnA promoter, and (iii) 225 ng of plasmid that expresses mCherry from the MtnA promoter. 24 hr after transfection, $CuSO_4$ was added for 14 hr and total protein **(A)** or RNA **(B)** was then isolated. **(A)** Representative Western blot using an $\alpha$-HA antibody to confirm Cas13 effector protein expression. $\alpha$-Tubulin was used as a loading control. **(B)** Northern blots using 8% polyacrylamide gels were performed to detect guide RNA expression. Expected guide RNA lengths: RxCas13d (59 nt), PspCas13b (75 nt), LwaCas13a (71 nt). tRNA:val4:70BCb was used as a loading control. **(C)** The guide RNAs complementary to eGFP, mCherry, nanoluciferase (nLuc), firefly luciferase (FFLuc), and Renilla luciferase (RLuc) that were used in this study.

Supplementary Figure S2

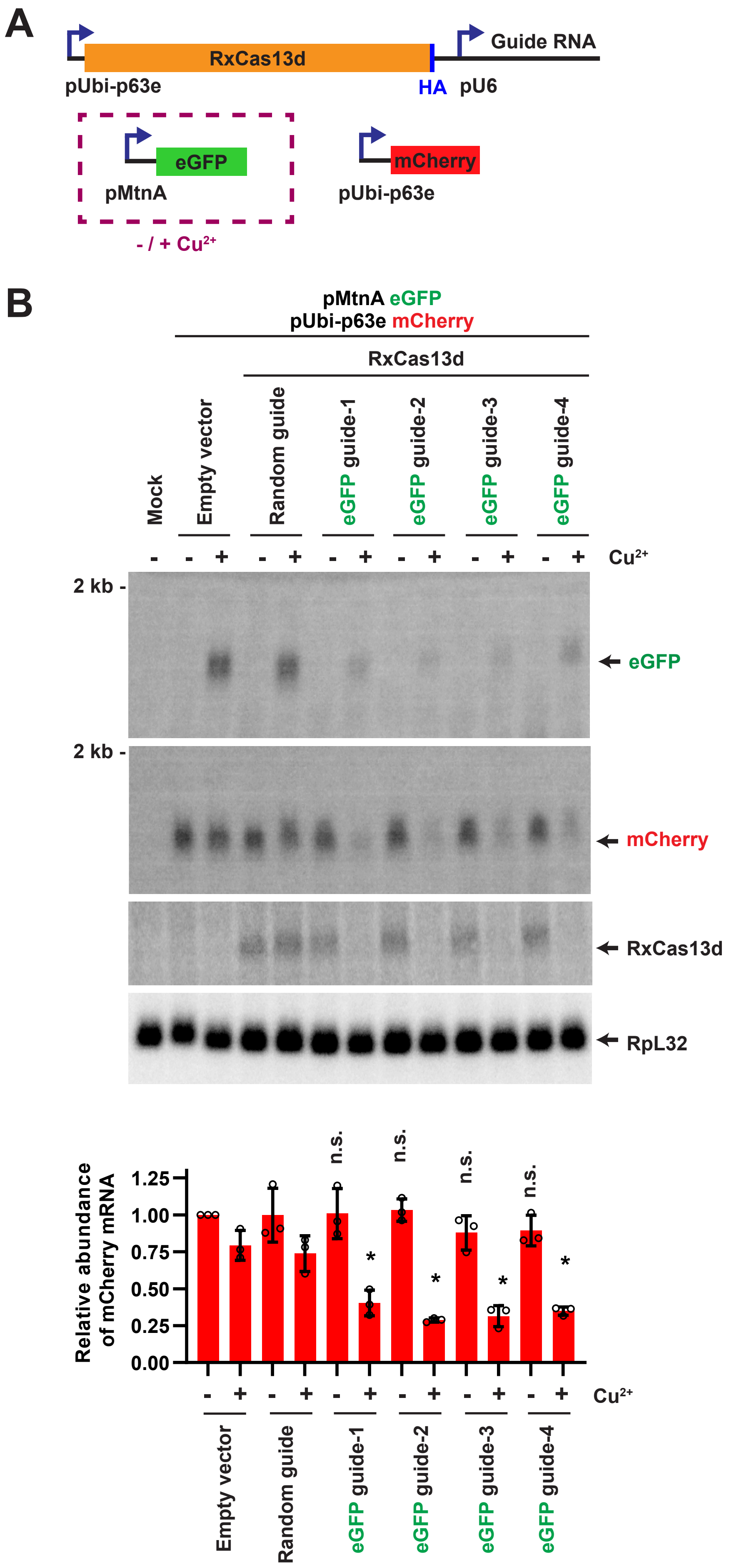

**Supplementary Figure S2. RxCas13d off-target effects are dependent on transcription of the target mRNA in *Drosophila* cells. (A)** *Drosophila* DL1 cells were co-transfected with (i) 50 ng of plasmid that constitutively expresses a guide RNA from the U6 promoter as well as catalytically active RxCas13d from the Ubi-p63e promoter, (ii) 225 ng of plasmid that expresses eGFP from the copper-inducible MtnA promoter, and (iii) 225 ng of plasmid that expresses mCherry from the Ubi-p63e promoter. After 24 hr, expression of eGFP was (+) or was not (-) induced by adding $CuSO_4$ for 14 hr, as indicated. **(B)** Total RNA was then isolated and Northern blots used to quantify the relative expression levels of eGFP, mCherry, and RxCas13d mRNAs. Representative blots are shown. mCherry mRNA expression data are shown as mean ± SD, N=3. For statistical comparisons, data were compared to the random guide RNA samples. (∗) $P < 0.05$. n.s., not significant.

**Supplementary Figure S3**

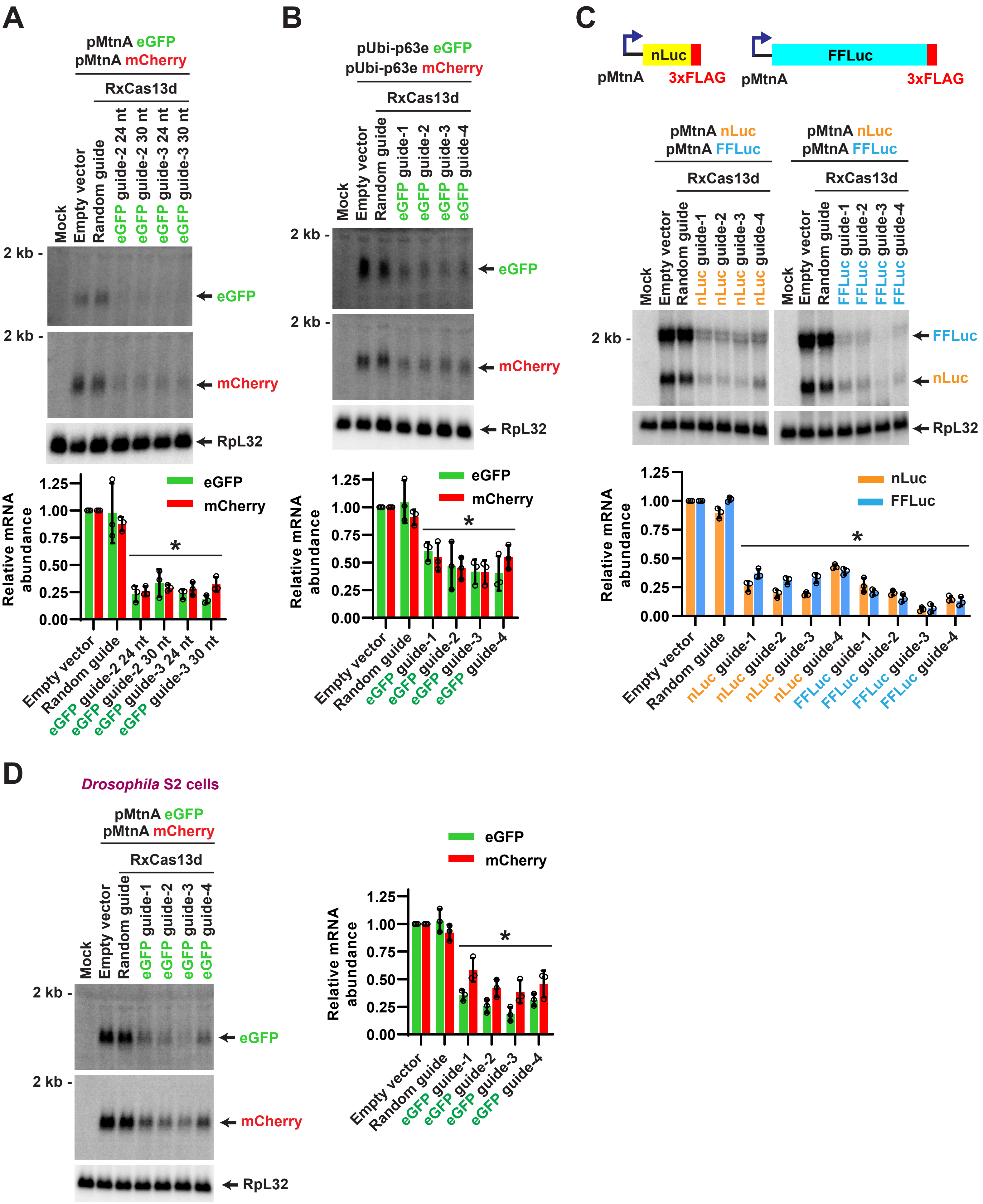

**Supplementary Figure S3. Strong off-target effects of RxCas13d are observed under multiple conditions in *Drosophila* cells. (A)** *Drosophila* DL1 cells were co-transfected with RxCas13d-guide RNA (50 ng), eGFP (225 ng), and mCherry (225 ng) expression plasmids as described in Figure 1A except that two different lengths (24 and 30 nt) of guide RNA spacers were tested. 24 hr after transfection, $CuSO_4$ was added and total RNA was isolated after an additional 14 hr. Northern blots were then performed to quantify expression of eGFP and mCherry mRNAs. **(B)** *Drosophila* DL1 cells were co-transfected as in Figure 1A except that eGFP and mCherry were expressed from the constitutive Ubi-p63e promoter (not the copper-inducible MtnA promoter). 40 hr after transfection, total RNA was isolated and Northern blots performed. **(C)** *Drosophila* DL1 cells were co-transfected with (i) 50 ng of plasmid expressing RxCas13d and a guide RNA complementary to nLuc (left) or FFLuc (right), (ii) 225 ng of plasmid that expresses 3xFLAG tagged nLuc from the copper-inducible MtnA promoter, and (iii) 225 ng of plasmid that expresses 3xFLAG tagged FFLuc from the MtnA promoter. 24 hr after transfection, $CuSO_4$ was added and total RNA was isolated after an additional 14 hr. Northern blotting using a probe complementary to the 3xFLAG tag was then performed. **(D)** *Drosophila* S2 cells were co-transfected with (i) 200 ng of plasmid that expresses RxCas13d and a guide RNA complementary to eGFP, (ii) 900 ng of plasmid that expresses eGFP from the copper-inducible MtnA promoter, and (iii) 900 ng of plasmid that expresses mCherry from the MtnA promoter. 24 hr after transfection, $CuSO_4$ was added and total RNA was isolated after an additional 14 hr. Northern blotting was then performed. For all panels **(A-D)**, representative Northern blots (20 $\mu$g of total RNA/lane) are shown. ImageQuant was used to quantify the relative expression levels of the indicated mRNAs (normalized to the empty vector samples) from three independent experiments. RpL32 mRNA served as an endogenous loading control. Data are shown as mean ± SD. For statistical comparisons, data were compared to the random guide RNA samples. ($*$) $P < 0.05$.

Supplementary Figure S4

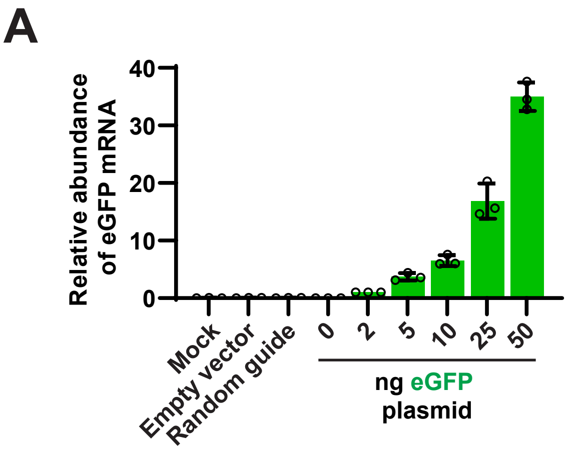

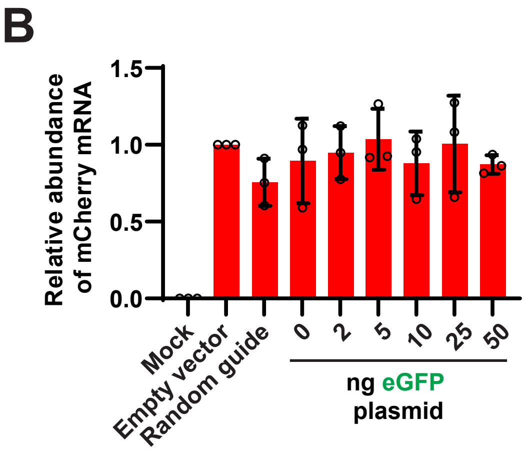

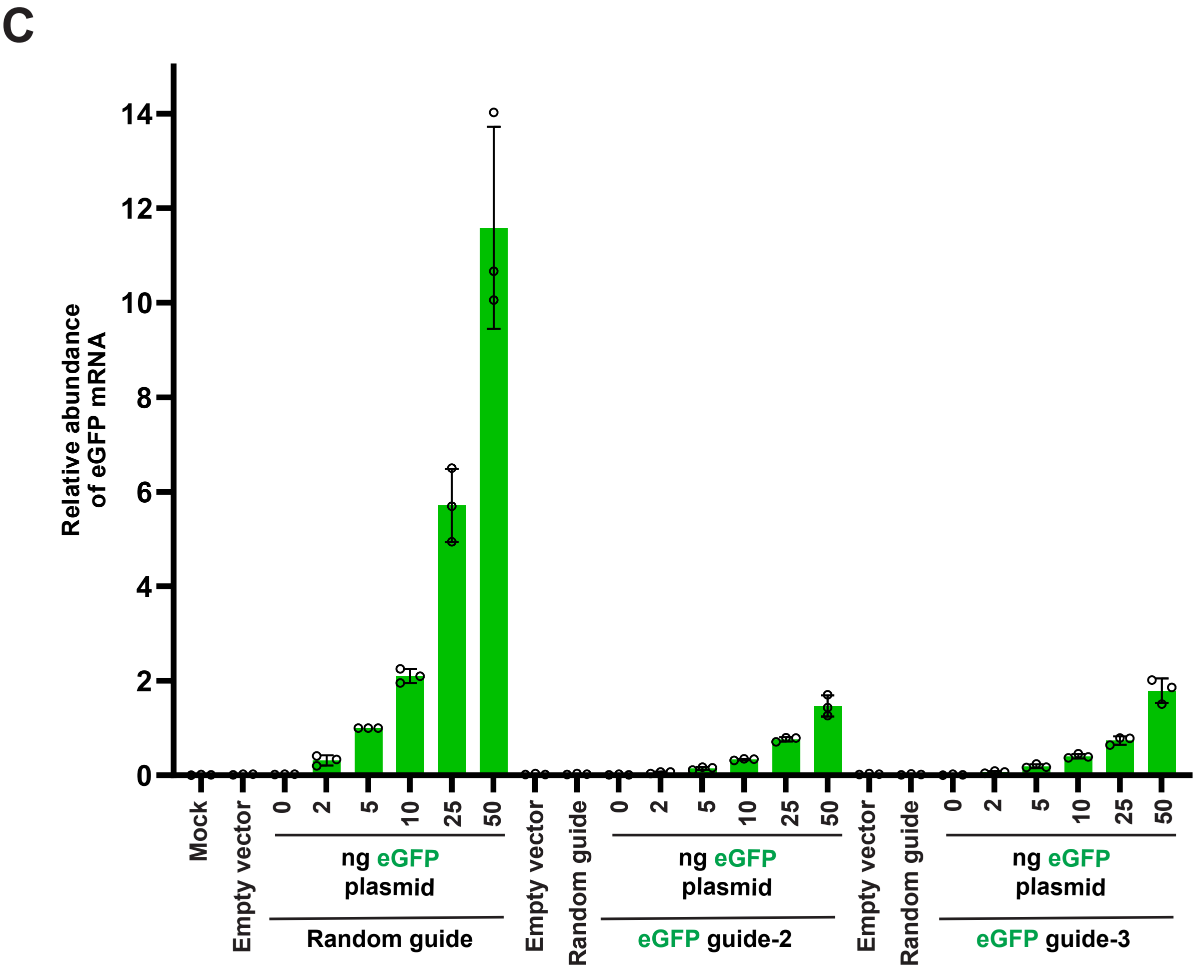

**Supplementary Figure S4. Titration of eGFP target mRNA expression to characterize dose-dependent off-target effects of RxCas13d in *Drosophila* cells. (A-B)** *Drosophila* DL1 cells were transfected with increasing amounts of copper-inducible eGFP expression plasmid (0, 2, 5, 10, 25, or 50 ng) along with a constant amount (225 ng) of copper-inducible mCherry expression plasmid. Empty vector (pUb-3xFLAG MCS (No BsmBI) plasmid) was added as needed so that 500 ng DNA was transfected in all samples. 24 hr after transfection, $CuSO_4$ was added for 14 hr and total RNA then isolated. RT-qPCR was used to quantify relative expression of eGFP **(A)** and mCherry **(B)** mRNAs. eGFP data were normalized to the 'eGFP 2 ng' sample, while mCherry data were normalized to the 'empty vector' samples. Data are shown as mean ± SD, N=3. **(C)** Same RT-qPCR data as in Figure 3B, but with all eGFP expression levels normalized to the 'eGFP 5 ng - Random guide RNA' sample. Data are shown as mean ± SD, N=3.

Supplementary Figure S5

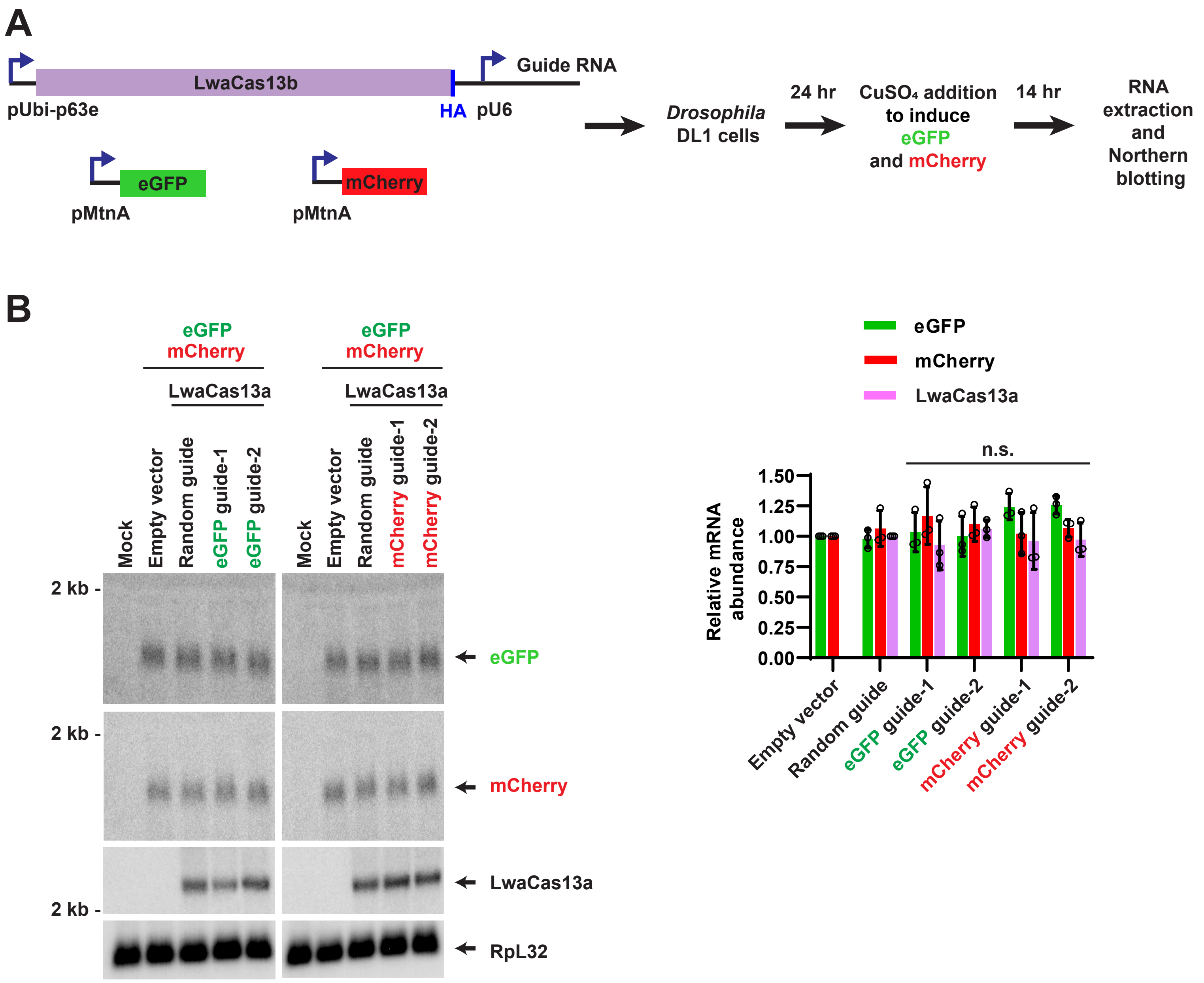

**Supplementary Figure S5. LwaCas13a failed to deplete target RNAs in *Drosophila* cells.** **(A)** Co-transfection assay analogous to Figure 1A except that LwaCas13a rather than RxCas13d was examined. *Drosophila* DL1 cells were co-transfected with (i) 50 ng of plasmid that constitutively expresses a guide RNA from the U6 promoter as well as HA-tagged LwaCas13a from the Ubi-p63e promoter, (ii) 225 ng of plasmid that expresses eGFP from the copper-inducible MtnA promoter, and (iii) 225 ng of plasmid that expresses mCherry from the MtnA promoter. 24 hr after transfection, $CuSO_4$ was added and total RNA was isolated after an additional 14 hr. Northern blots were then performed. **(B)** Representative Northern blots (20 μg of total RNA/lane) are shown from experiments in which guide RNAs complementary to eGFP (left) or mCherry (right) were used. ImageQuant was used to quantify the relative expression levels of eGFP, mCherry, and LwaCas13a mRNAs from three independent experiments. eGFP and mCherry mRNA expression was normalized to the empty vector samples, while LwaCas13a mRNA expression was normalized to the random guide RNA samples. Data are shown as mean ± SD. n.s., not significant.

Supplementary Figure S6

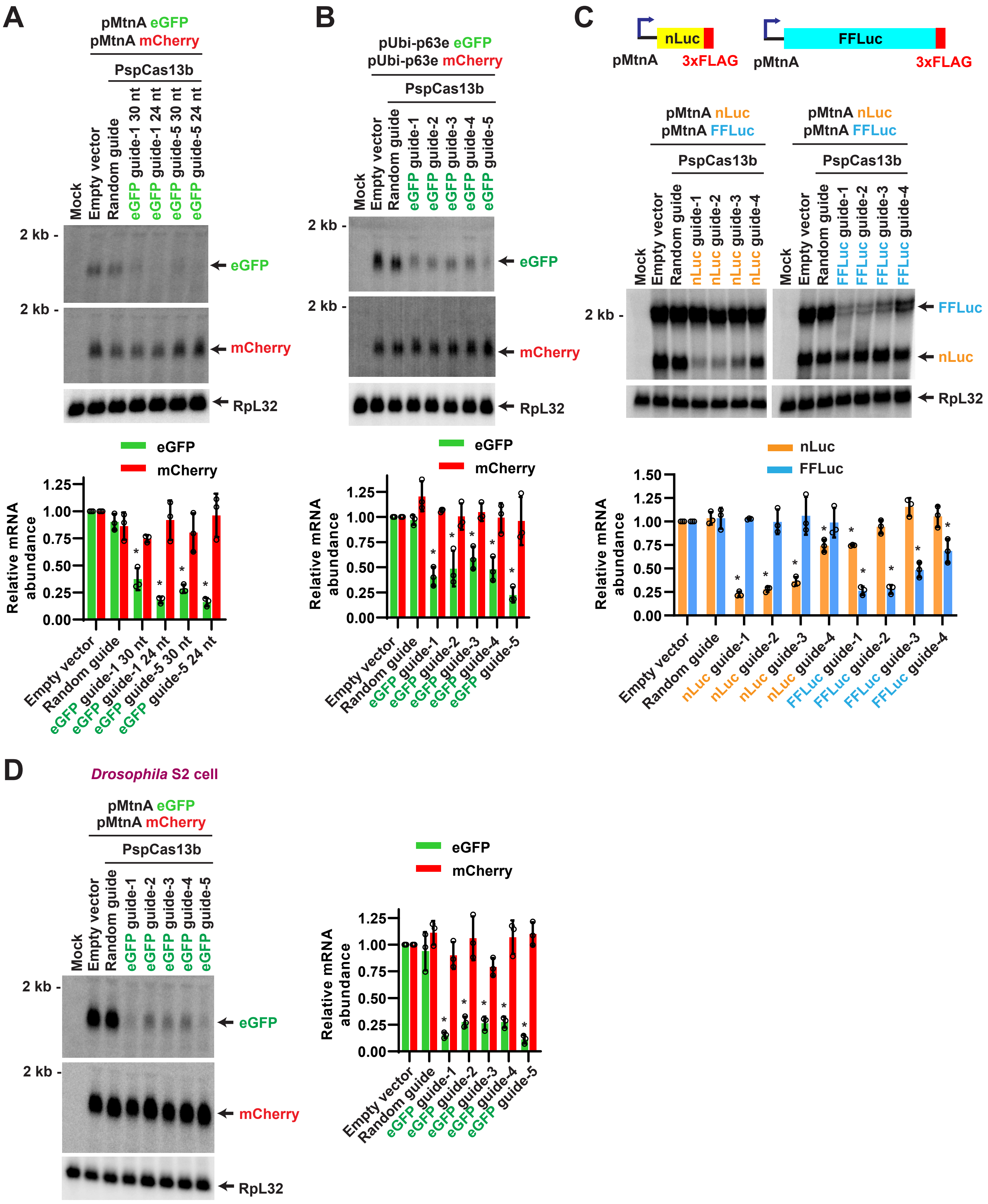

**Supplementary Figure S6. PspCas13b has good specificity under multiple conditions in *Drosophila* cells.** These experiments are the same as those performed in Supplementary Figure S3 except that PspCas13b rather than RxCas13d was examined. **(A)** *Drosophila* DL1 cells were co-transfected with PspCas13b-guide RNA (50 ng), eGFP (225 ng), and mCherry (225 ng) expression plasmids. Two different lengths (24 and 30 nt) of guide RNA spacers were tested. 24 hr after transfection, $CuSO_4$ was added and total RNA was isolated after an additional 14 hr. Northern blots were then performed to quantify expression of eGFP and mCherry mRNAs. **(B)** *Drosophila* DL1 cells were co-transfected as in Figure 4A except that the eGFP and mCherry expression plasmids were driven by the constitutive Ubi-p63e promoter (not the copper-inducible MtnA promoter). 40 hr after transfection, total RNA was isolated and Northern blots performed. **(C)** *Drosophila* DL1 cells were co-transfected with (i) 50 ng of plasmid expressing PspCas13b and a guide RNA complementary to nLuc (left) or FFLuc (right), (ii) 225 ng of plasmid that expresses 3xFLAG tagged nLuc from the copper-inducible MtnA promoter, and (iii) 225 ng of plasmid that expresses 3xFLAG tagged FFLuc from the MtnA promoter. 24 hr after transfection, $CuSO_4$ was added and total RNA was isolated after an additional 14 hr. Northern blotting using a probe complementary to the 3xFLAG tag was then performed. **(D)** *Drosophila* S2 cells were co-transfected with (i) 500 ng of plasmid that expresses PspCas13b and a guide RNA complementary to eGFP, (ii) 750 ng of plasmid that expresses eGFP from the copper-inducible MtnA promoter, and (iii) 750 ng of plasmid that expresses mCherry from the MtnA promoter. 24 hr after transfection, $CuSO_4$ was added and total RNA was isolated after an additional 14 hr. Northern blotting was then performed. For all panels **(A-D)**, representative Northern blots (20 $\mu$g of total RNA/lane) are shown. ImageQuant was used to quantify the relative expression levels of the indicated mRNAs (normalized to the empty vector samples) from three independent experiments. RpL32 mRNA served as an endogenous loading control. Data are shown as mean ± SD. For statistical comparisons, data were compared to the random guide RNA samples. (∗) $P < 0.05$.

Supplementary Figure S7

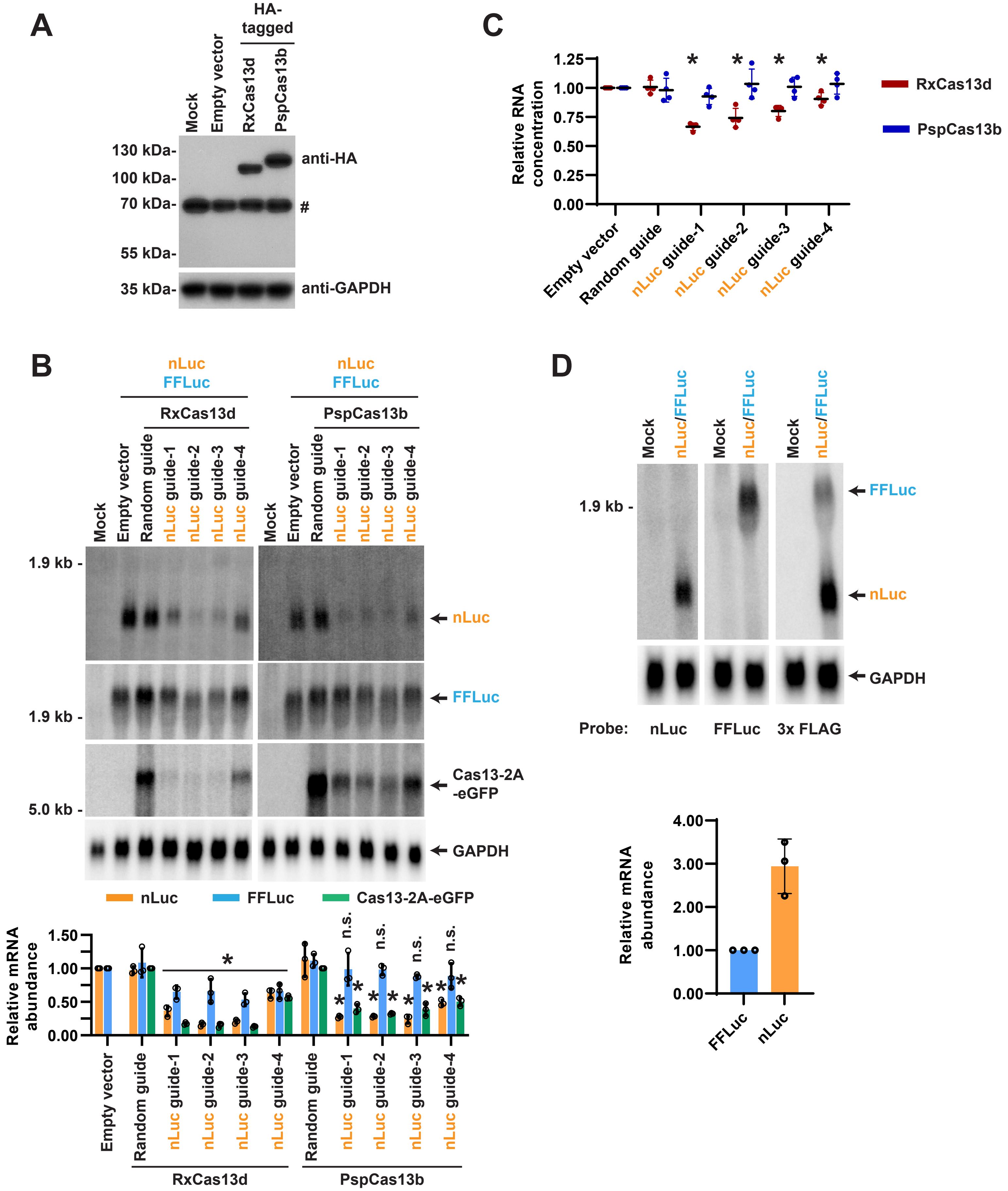

**Supplementary Figure S7. Specificity of RxCas13d and PspCas13b in HeLa cells when nLuc and FFLuc were co-transfected. (A-C)** Analogous to Figure 6A, HeLa cells were co-transfected with (i) 300 ng of plasmid that expresses HA-tagged Cas13 protein followed by a 2A peptide and eGFP, (ii) 200 ng of plasmid that expresses a guide RNA, (iii) 250 ng of plasmid that expresses 3xFLAG-tagged nLuc, and (iv) 250 ng of plasmid that expresses 3xFLAG-tagged FFLuc. 48 hr after transfection, total protein **(A)** or RNA **(B-C)** was isolated. **(A)** A representative Western blot using an $\alpha$-HA antibody to confirm Cas13 protein expression (Note: random guide RNA plasmid was co-transfected). $\alpha$-GAPDH was used as a loading control. # denotes non-specific band. **(B)** Representative Northern blots (20 $\mu$g of total RNA/lane) are shown. ImageQuant was used to quantify the relative expression levels of nLuc, FFLuc, and Cas13-2A-eGFP mRNAs. nLuc and FFLuc mRNA expression was normalized to the empty vector (pBEVY-L) samples, while Cas13-2A-eGFP mRNA expression was normalized to the random guide RNA samples. GAPDH mRNA served as an endogenous loading control. Data are shown as mean ± SD, N=3. For statistical comparisons, data were compared to the random guide RNA samples. (∗) $P < 0.05$. **(C)** Relative RNA concentrations obtained from the co-transfection assays when the RxCas13d or PspCas13b expression plasmids were used. Data are normalized to the empty vector samples and shown as mean ± SD, N=4. (∗) $P < 0.05$. **(D)** HeLa cells were co-transfected with (i) 250 ng of plasmid that expresses 3xFLAG-tagged nLuc, (ii) 250 ng of plasmid that expresses 3xFLAG-tagged FFLuc, and (iii) 500 ng of empty vector (pBEVY-L). 48 hr after transfection, total RNA was isolated. Northern blots were then performed using probes complementary to nLuc, FFLuc, or the 3xFLAG that detects both nLuc and FFLuc mRNAs. ImageQuant was used to quantify the relative expression levels of nLuc and FFLuc mRNAs that were detected using the 3x FLAG probe. nLuc expression was normalized to FFLuc expression. Data are shown as mean ± SD, N=3.

#### Supplementary Figure S8

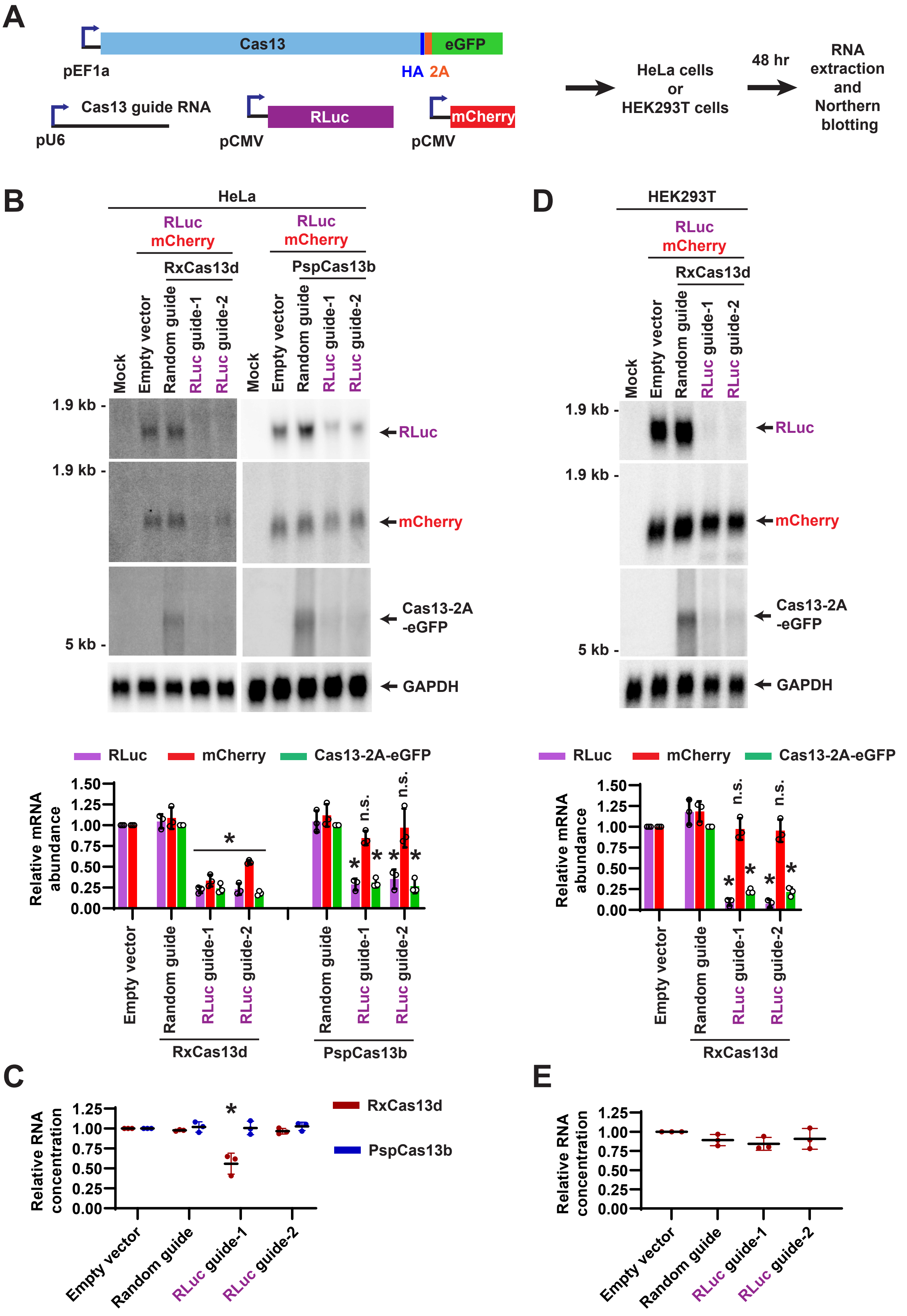

**Supplementary Figure S8. Specificity of Cas13 effectors when RLuc and mCherry were co-transfected into different human cell lines. (A)** HeLa or HEK293T cells were co-transfected with (i) 300 ng of plasmid that constitutively expresses HA-tagged Cas13 protein followed by a 2A peptide and eGFP, (ii) 200 ng of plasmid that expresses a guide RNA, (iii) 250 ng of plasmid that expresses Renilla luciferase (RLuc), and (iv) 250 ng of plasmid that expresses mCherry. 48 hr after transfection, total RNA was isolated and Northern blots performed. **(B-C)** Guide RNAs complementary to RLuc were employed in the co-transfection assay in HeLa cells. **(B)** Representative Northern blots (20 μg of total RNA/lane) are shown. ImageQuant was used to quantify the relative expression levels of RLuc, mCherry, and Cas13-2A-eGFP mRNAs. RLuc and mCherry mRNA expression was normalized to the empty vector (pBEVY-L) samples, while Cas13-2A-eGFP mRNA expression was normalized to the random guide RNA samples. GAPDH mRNA served as an endogenous loading control. Data are shown as mean ± SD, N=3. For statistical comparisons, data were compared to the random guide RNA samples. (*) $P < 0.05$. **(C)** Relative RNA concentrations obtained from the HeLa co-transfection assays when the RxCas13d or PspCas13b expression plasmids were used. Data are normalized to the empty vector samples and shown as mean ± SD, N=3. (*) $P < 0.05$. **(D-E)** RxCas13d and guide RNAs complementary to RLuc were employed in the co-transfection assay in HEK293T cells as diagrammed in A. **(D)** Representative Northern blots (20 μg of total RNA/lane) are shown. ImageQuant was used to quantify the relative expression levels of RLuc, mCherry, and Cas13-2A-eGFP mRNAs. RLuc and mCherry mRNA expression was normalized to the empty vector (pBEVY-L) samples, while Cas13-2A-eGFP mRNA expression was normalized to the random guide RNA samples. GAPDH mRNA served as an endogenous loading control. Data are shown as mean ± SD, N=3. For statistical comparisons, data were compared to the random guide RNA samples. (*) $P < 0.05$. **(E)** Relative RNA concentrations obtained from the HEK293T co-transfection assays. Data are normalized to the empty vector samples and shown as mean ± SD, N=3. No significant changes were observed.

#### Supplementary Figure S9

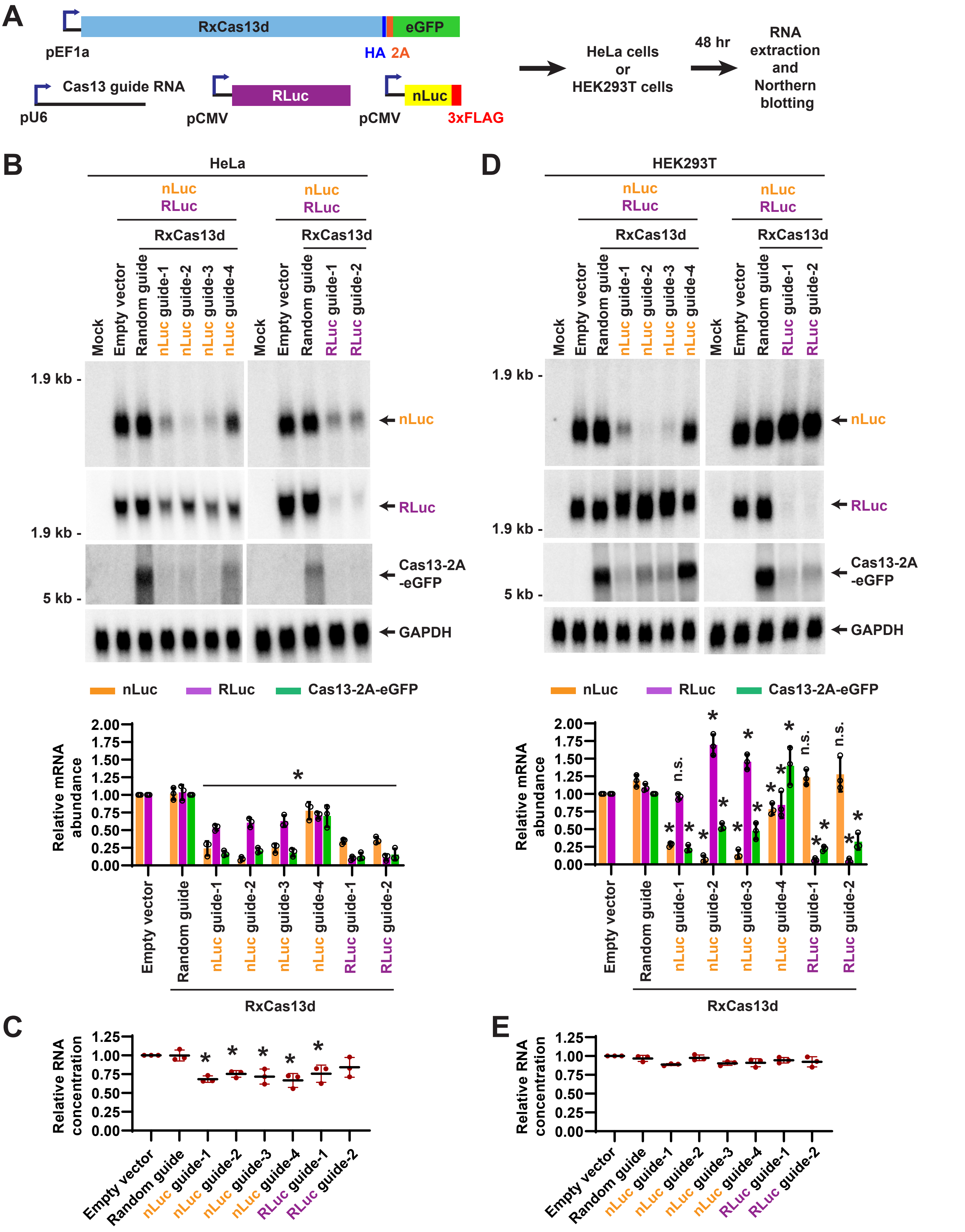

**Supplementary Figure S9. Specificity of RxCas13d when RLuc and nLuc were co-transfected into HeLa or HEK293T cells. (A)** HeLa or HEK293T cells were co-transfected with (i) 300 ng of plasmid that constitutively expresses HA-tagged Cas13 protein followed by a 2A peptide and eGFP, (ii) 200 ng of plasmid that expresses a guide RNA, (iii) 250 ng of plasmid that expresses Renilla luciferase (RLuc), and (iv) 250 ng of plasmid that expresses 3xFLAG-tagged nLuc. 48 hr after transfection, total RNA was isolated and Northern blots performed. **(B-E)** Guide RNAs complementary to nLuc or RLuc were employed in the co-transfection assay in HeLa cells **(B-C)** or HEK293T cells **(D-E)**. **(B, D)** Representative Northern blots (20 μg of total RNA/lane) are shown. ImageQuant was used to quantify the relative expression level of RLuc, nLuc, and Cas13-2A-eGFP mRNAs. RLuc and nLuc mRNA expression was normalized to the empty vector (pBEVY-L) samples, while Cas13-2A-eGFP mRNA expression was normalized to the random guide RNA samples. GAPDH mRNA served as an endogenous loading control. Data are shown as mean ± SD, N=3. For statistical comparisons, data were compared to the random guide RNA samples. (∗) $P < 0.05$. **(C, E)** Relative RNA concentrations obtained from the co-transfection assays. Data are normalized to the empty vector samples and shown as mean ± SD, N=3. (∗) $P < 0.05$. n.s., not significant.

#### SUPPLEMENTARY TABLES

### Supplementary Table S1

#### Guide RNAs used in *Drosophila* cells

| Cas protein | Guide RNA | Forward primer | Reverse primer |
|---|---|---|---|
| RxCas13d | Random guide (DL1) | gaaacCGAGGGCGACTTAACCTTAGGTt | aaaaaACCTAAGGTTAAGTCGCCCTCGg |
| RxCas13d | Random guide (S2) | gaaacGCAGGGTTTTCCCAGTCACGACGTt | aaaaaACGTCGTGACTGGGAAAACCCTGCg |
| RxCas13d | eGFP guide-1 | GAAACGGATGGGCACCACCCCGGTGAACAT | AAAAATGTTCACCGGGGTGGTGCCCATCCG |
| RxCas13d | eGFP guide-2 | GAAACTCATGTGGTCGGGGTAGCGGCTGAT | AAAAATCAGCCGCTACCCCGACCACATGAG |
| RxCas13d | eGFP guide-3 | GAAACACTCCAGCTTGTGCCCCAGGATGTT | AAAAAACATCCTGGGGCACAAGCTGGAGTG |
| RxCas13d | eGFP guide-4 | GAAACGGGCGGACTGGGTGCTCAGGTAGTT | AAAAAACTACCTGAGCACCCAGTCCGCCCG |
| RxCas13d | mCherry guide-1 | gaaacgccttggagccgtacatgaactgat | aaaaatcagttcatgtacggctccaaggcg |
| RxCas13d | mCherry guide-2 | gaaaccttctgcattacggggccgtcggat | aaaaatccgacggccccgtaatgcagaagg |
| RxCas13d | Laccase2 BSJ guide | GAAACGGTAACCGCTTCCAGCGGTTAGGGT | AAAAAGGGGAGCTCAGCAAAGGTACCAAAG |
| RxCas13d | Laccase2 Exon 2 guide | GAAACGGTAACCGCTTCCAGCGGTTAGGGT | AAAAACCCTAACCGCTGGAAGCGGTTACCG |
| RxCas13d | Laccase2 Exon 3 guide | GAAACGTCCATCTAGTTTCTGTAGACCGGT | AAAAACCGGTCTACAGAAACTAGATGGACG |
| RxCas13d | nLuc guide-1 | GAAACGGAGTTACGGACACCCCGAGATTCT | AAAAAGAATCTCGGGGTGTCCGTAACTCCG |
| RxCas13d | nLuc guide-2 | GAAACTTCATACGGGATGATGACATGGATT | AAAAAATCCATGTCATCATCCCGTATGAAG |
| RxCas13d | nLuc guide-3 | GAAACCGTAACCCCGTCGATTACCAGTGTT | AAAAAACACTGGTAATCGACGGGGTTACGG |
| RxCas13d | nLuc guide-4 | GAAACCAGTGATCTTTTTGCCGTCGAACAT | AAAAATGTTCGACGGCAAAAAGATCACTGG |
| RxCas13d | FFLuc guide-1 | GAAACaagctattctcgctgcacaccacgT | AAAAAcgtggtgtgcagcgagaatagcttG |
| RxCas13d | FFLuc guide-2 | GAAACgtactcgttgaagccgggtggcaaT | AAAAAttgccacccggcttcaacgagtacG |
| RxCas13d | FFLuc guide-3 | GAAACatagctcctcctcgaagcggtacaT | AAAAAtgtaccgcttcgaggaggagctatG |
| RxCas13d | FFLuc guide-4 | GAAACaccaccttgcctactgcgccaggcT | AAAAAgcctggcgcagtaggcaaggtggtG |
| RxCas13d | eGFP guide-2 30 nt | GAAACGCTGCTTCATGTGGTCGGGGTAGCGGCTGAT | AAAAATCAGCCGCTACCCCGACCACATGAAGCAGCG |
| RxCas13d | eGFP guide-3 30 nt | GAAACAGTTGTACTCCAGCTTGTGCCCCAGGATGTT | AAAAAACATCCTGGGGCACAAGCTGGAGTACAACTG |
| PspCas13b | Random guide (DL1) | cttcggagacggtCGAGGGCGACTTAACCTTAGGT | caacACCTAAGGTTAAGTCGCCCTCGaccgtctcc |
| PspCas13b | Random guide (S2) | cttcgGCAGGGTTTTCCCAGTCACGACGTTGTAAA | caacTTTACAACGTCGTGACTGGGAAAACCCTGCc |
| PspCas13b | eGFP guide-1 | cttcgGCTGCTTCATGTGGTCGGGGTAGCGGCTGA | caacTCAGCCGCTACCCCGACCACATGAAGCAGCc |
| PspCas13b | eGFP guide-2 | cttcgGACGTAGCCTTCGGGCATGGCGGACTTGAA | caacTTCAAGTCCGCCATGCCCGAAGGCTACGTCc |
| PspCas13b | eGFP guide-3 | cttcgAGCTTGCCGTAGGTGGCATCGCCCTCGCCC | caacGGGCGAGGGCGATGCCACCTACGGCAAGCTc |
| PspCas13b | eGFP guide-4 | cttcgCGGGCAGCTTGCCGGTGGTGCAGATGAACT | caacAGTTCATCTGCACCACCGGCAAGCTGCCCGc |
| PspCas13b | eGFP guide-5 | cttcgGGGGGTGTTCTGCTGGTAGTGGTCGGCGAG | caacCTCGCCGACCACTACCAGCAGAACACCCCCc |
| PspCas13b | mCherry guide-1 | cttcgacgtaggccttggagccgtacatgaactga | caactcagttcatgtacggctccaaggcctacgtc |
| PspCas13b | mCherry guide-2 | cttcgggtcttcttctgcattacggggccgtcgga | caactccgacggccccgtaatgcagaagaagaccc |
| PspCas13b | mCherry guide-3 | cttcgtcaccacgccgccgtcctcgaagttcatca | caactgatgaacttcgaggacggcggcgtggtgac |
| PspCas13b | mCherry guide-4 | cttcgcttcaccttgtagatgaactcgccgtcctg | caaccaggacggcgagttcatctacaaggtgaagc |
| PspCas13b | mCherry guide-5 | cttcgtctgcttgatctcgcccttcagggcgccgt | caacacggcgccctgaagggcgagatcaagcagac |
| PspCas13b | Laccase2 BSJ guide | cttcgGATTTTGGTACCTTTGCTGAGCTCCCCGGG | caacCCCGGGGAGCTCAGCAAAGGTACCAAAATCc |
| PspCas13b | Laccase2 Exon 2 guide | cttcgTTAGGTAACCGCTTCCAGCGGTTAGGGCTG | caacCAGCCCTAACCGCTGGAAGCGGTTACCTAAc |
| PspCas13b | Laccase2 Exon 3 guide | cttcgAGAGTCCATCTAGTTTCTGTAGACCGGTAT | caacATACCGGTCTACAGAAACTAGATGGACTCTc |
| PspCas13b | nLuc guide-1 | cttcgTGGATCGGAGTTACGGACACCCCGAGATTC | caacGAATCTCGGGGTGTCCGTAACTCCGATCCAc |
| PspCas13b | nLuc guide-2 | cttcgCAGACCTTCATACGGGATGATGACATGGAT | CaacATCCATGTCATCATCCCGTATGAAGGTCTGc |
| PspCas13b | nLuc guide-3 | cttcgGTTCGGCGTAACCCCGTCGATTACCAGTGT | caacACACTGGTAATCGACGGGGTTACGCCGAACc |
| PspCas13b | nLuc guide-4 | cttcgCTGTTACAGTGATCTTTTTGCCGTCGAACA | caacTGTTCGACGGCAAAAAGATCACTGTAACAGc |
| PspCas13b | FFLuc guide-1 | cttcgaactgcaagctattctcgctgcacaccacg | caaccgtggtgtgcagcgagaatagcttgcagttc |
| PspCas13b | FFLuc guide-2 | cttcggaagtcgtactcgttgaagccgggtggcaa | Caacttgccacccggcttcaacgagtacgacttcc |
| PspCas13b | FFLuc guide-3 | cttcggcaagaatagctcctcctcgaagcggtaca | caactgtaccgcttcgaggaggagctattcttgcc |
| PspCas13b | FFLuc guide-4 | cttcgaagggcaccaccttgcctactgcgccaggc | caacgcctggcgcagtaggcaaggtggtgcccttc |
| PspCas13b | eGFP guide-1 24 nt | cttcgTCATGTGGTCGGGGTAGCGGCTGA | caacTCAGCCGCTACCCCGACCACATGAc |
| PspCas13b | eGFP guide-5 24 nt | cttcgACTCCAGCTTGTGCCCCAGGATGT | caacACATCCTGGGGCACAAGCTGGAGTc |
| LwaCas13a | eGFP guide-1 | aaaacTGCTTCATGTGGTCGGGGTAGCGGCTGAt | aaaaaTCAGCCGCTACCCCGACCACATGAAGCAg |
| LwaCas13a | eGFP guide-2 | aaaacTTGTACTCCAGCTTGTGCCCCAGGATGTt | aaaaaACATCCTGGGGCACAAGCTGGAGTACAAg |
| LwaCas13a | mCherry guide-1 | aaaacgtaggccttggagccgtacatgaactgat | aaaaatcagttcatgtacggctccaaggcctacg |
| LwaCas13a | mCherry guide-2 | aaaactcttcttctgcattacggggccgtcggat | aaaaatccgacggccccgtaatgcagaagaagag |

### Supplementary Table S2

#### Guide RNAs used in human cells

| Cas protein | Guide RNA | Forward primer | Reverse primer |
|---|---|---|---|
| RxCas13d | Random guide | aaaccGCAGGGTTTTCCCAGTCACGACGTt | aaaaaACGTCGTGACTGGGAAAACCCTGCg |
| RxCas13d | nLuc guide-1 | AAACCGGAGTTACGGACACCCCGAGATTCT | AAAAAGAATCTCGGGGTGTCCGTAACTCCG |
| RxCas13d | nLuc guide-2 | AAACCTTCATACGGGATGATGACATGGATT | AAAAAATCCATGTCATCATCCCGTATGAAG |
| RxCas13d | nLuc guide-3 | AAACCCGTAACCCCGTCGATTACCAGTGTT | AAAAAACACTGGTAATCGACGGGGTTACGG |
| RxCas13d | nLuc guide-4 | AAACCCAGTGATCTTTTTGCCGTCGAACAT | AAAAATGTTCGACGGCAAAAAGATCACTGG |
| RxCas13d | FFLuc guide-1 | AAACCaagctattctcgctgcacaccacgT | AAAAAcgtggtgtgcagcgagaatagcttG |
| RxCas13d | FFLuc guide-2 | AAACCgtactcgttgaagccgggtggcaaT | AAAAAttgccacccggcttcaacgagtacG |
| RxCas13d | FFLuc guide-3 | AAACCatagctcctcctcgaagcggtacaT | AAAAAtgtaccgcttcgaggaggagctatG |
| RxCas13d | FFLuc guide-4 | AAACCaccaccttgcctactgcgccaggcT | AAAAAgcctggcgcagtaggcaaggtggtG |
| RxCas13d | RLuc guide-1 | AAACCacttacccattccgatcagatcagT | AAAAActgatctgatcggaatgggtaagtG |
| RxCas13d | RLuc guide-2 | AAACCcccaggactcgatcacgtccacgaT | AAAAAtcgtggacgtgatcgagtcctgggG |
| PspCas13b | Random guide | caccgGCAGGGTTTTCCCAGTCACGACGTTGTAAA | caacTTTACAACGTCGTGACTGGGAAAACCCTGCc |
| PspCas13b | nLuc guide-1 | caccgTGGATCGGAGTTACGGACACCCCGAGATTC | caacGAATCTCGGGGTGTCCGTAACTCCGATCCAc |
| PspCas13b | nLuc guide-2 | caccgCAGACCTTCATACGGGATGATGACATGGAT | CaacATCCATGTCATCATCCCGTATGAAGGTCTGc |
| PspCas13b | nLuc guide-3 | caccgGTTCGGCGTAACCCCGTCGATTACCAGTGT | caacACACTGGTAATCGACGGGGTTACGCCGAACc |
| PspCas13b | nLuc guide-4 | caccgCTGTTACAGTGATCTTTTTGCCGTCGAACA | caacTGTTCGACGGCAAAAAGATCACTGTAACAGc |
| PspCas13b | FFLuc guide-1 | caccgaactgcaagctattctcgctgcacaccacg | caaccgtggtgtgcagcgagaatagcttgcagttc |
| PspCas13b | FFLuc guide-2 | caccggaagtcgtactcgttgaagccgggtggcaa | Caacttgccacccggcttcaacgagtacgacttcc |
| PspCas13b | FFLuc guide-3 | caccggcaagaatagctcctcctcgaagcggtaca | caactgtaccgcttcgaggaggagctattcttgcc |
| PspCas13b | FFLuc guide-4 | caccgaagggcaccaccttgcctactgcgccaggc | caacgcctggcgcagtaggcaaggtggtgcccttc |
| PspCas13b | RLuc guide-1 | caccgtgccggacttacccattccgatcagatcag | caacctgatctgatcggaatgggtaagtccggcac |
| PspCas13b | RLuc guide-2 | caccgactcgtcccaggactcgatcacgtccacga | caactcgtggacgtgatcgagtcctgggacgagtc |

### Supplementary Table S3
**Northern blot probes**

| Target transcript | Probe sequence | Figure panels |
|---|---|---|
| eGFP | GTCACGAACTCCAGCAGGAC | 1B, 1C, 2C, 4B, 4C, S2B, S3A, S3B, S3D, S5B, S6A, S6B, S6D |
| mCherry | CGCCGGTGGAGTGGCGGCCC | 1B, 1C, 2C, 3C, 4B, 4C, S2B, S3A, S3B, S3D, S5B, S6A, S6B, S6D, S8B, S8D |
| RxCas13d | CTCATTCCTATCAAAGAGGGCCTCAATA | 1B, 1C, 3C, S2B |
| PspCas13b or LwaCas13a (Complementary to HA tag) | TCAGGCACGTCGTAGGGGTA | 4B, 4C, S5B |
| RpL32 | GACGCACTCTGTTGTCGATACC | 1B, 1C, 2C, 3C, 4B, 4C, 5B, S2B, S3A, S3B, S3C, S3D, S5B, S6A, S6B, S6C, S6D |
| RxCas13d guide RNA | CCCCGACCAGTTGGTAGGG | S1B |
| PspCas13b guide RNA | CCTCAAAACTGGACCTTCCACAAC | S1B |
| LwaCas13a guide RNA | CCCCTTCGTTTTTGGGGTAGTCTAA | S1B |
| tRNA:val4:70BCb | GTTTCCGCCCGGGATCGAAC | S1B |
| Laccase2 minigene (Exon 2) | GTGTAAGATCTGGTTGATTTTGGTACCTTT | 5B |
| nLuc & FFLuc (Complementary to 3x FLAG) | CGATGTCATGATCTTTATAATCACCGTCATGG | S3C, S6C, S7D |
| nLuc | CGTAACCCCGTCGATTACCA | 6B, 6E, S7B, S7D, S9B, S9D |
| FFLuc | GCCCTTCTTGGCCTTAATGAGAATCT | 6B, S6B, S7B, S7D |
| Rluc | GCGTCCTCCTGGCTGAAGT | S8B, S8D, S9B, S9D |
| RxCas13d & PspCas13b-HA-eGFP (Complementary to HA tag) | AGCGTAATCTGGAACATCGTATGGGTA | 6B, 6E, S7B, S8B, S8D, S9B, S9D |
| GAPDH | GTTGCTGTAGCCAAATTCGTTGT | 6B, 6E, S7B, S7D, S8B, S8D, S9B, S9D |

### Supplementary Table S4
#### RT-qPCR primers

| Target RNA | Forward primer | Reverse primer |
|---|---|---|
| eGFP | CCTGAAGTTCATCTGCACCA | AAGTCGTGCTGCTTCATGTG |
| mCherry | CCTGTCCCCTCAGTTCATGT | CCGTCCTCGAAGTTCATCAC |
| *Drosophila* Act42a | CTCCGTCCACCATGAAGATT | TTCGAGATCCACATCTGCTG |
| FFLuc | GCAGTACCGGATTGCCCAAG | GTCGGGGATGATCTGGTTGC |
| Human GAPDH | GGTGGTCTCCTCTGACTTCAACA | GTTGCTGTAGCCAAATTCGTTGT |

### Supplementary Table S5

#### Full statistical analyses for titration assays

ns (not significant), * p<0.05; ** p<0.01, *** p<0.001, **** p<0.0001

##### Figure 3C: eGFP mRNA (eGFP guide-2)

| Tukey's multiple comparisons test | Mean Diff. | 95.00% CI of diff. | Summary | Adjusted P Value |
|---|---|---|---|---|
| 50 ng vs. 25 ng | -0.008231 | -0.05932 to 0.04286 | ns | 0.982 |
| 50 ng vs. 10 ng | -0.03191 | -0.08300 to 0.01918 | ns | 0.3086 |
| 50 ng vs. 5 ng | -0.02127 | -0.07236 to 0.02982 | ns | 0.6578 |
| 50 ng vs. 2 ng | -0.06465 | -0.1157 to -0.01356 | * | 0.013 |
| 25 ng vs. 10 ng | -0.02368 | -0.07477 to 0.02741 | ns | 0.5703 |
| 25 ng vs. 5 ng | -0.01304 | -0.06413 to 0.03805 | ns | 0.9118 |
| 25 ng vs. 2 ng | -0.05642 | -0.1075 to -0.005328 | * | 0.0294 |
| 10 ng vs. 5 ng | 0.01064 | -0.04045 to 0.06173 | ns | 0.9553 |
| 10 ng vs. 2 ng | -0.03274 | -0.08383 to 0.01835 | ns | 0.2877 |
| 5 ng vs. 2 ng | -0.04337 | -0.09447 to 0.007715 | ns | 0.1074 |

##### Figure 3C: eGFP mRNA (eGFP guide-3)

| Tukey's multiple comparisons test | Mean Diff. | 95.00% CI of diff. | Summary | Adjusted P Value |
|---|---|---|---|---|
| 50 ng vs. 25 ng | 0.02554 | -0.04430 to 0.09537 | ns | 0.7499 |
| 50 ng vs. 10 ng | -0.03679 | -0.1066 to 0.03305 | ns | 0.4574 |
| 50 ng vs. 5 ng | -0.03601 | -0.1058 to 0.03383 | ns | 0.4767 |
| 50 ng vs. 2 ng | -0.06428 | -0.1341 to 0.005560 | ns | 0.075 |
| 25 ng vs. 10 ng | -0.06233 | -0.1322 to 0.007509 | ns | 0.0864 |
| 25 ng vs. 5 ng | -0.06154 | -0.1314 to 0.008291 | ns | 0.0914 |
| 25 ng vs. 2 ng | -0.08981 | -0.1596 to -0.01998 | * | 0.0117 |
| 10 ng vs. 5 ng | 0.000782 | -0.06905 to 0.07062 | ns | >0.9999 |
| 10 ng vs. 2 ng | -0.02749 | -0.09732 to 0.04235 | ns | 0.6999 |
| 5 ng vs. 2 ng | -0.02827 | -0.09810 to 0.04157 | ns | 0.6793 |

##### Figure 3D: mCherry mRNA (Random guide)

| Tukey's multiple comparisons test | Mean Diff. | 95.00% CI of diff. | Summary | Adjusted P Value |
|---|---|---|---|---|
| Empty vector vs. 0 ng | 0.1339 | -0.1723 to 0.4400 | ns | 0.7446 |
| Empty vector vs. 2 ng | 0.1275 | -0.1786 to 0.4337 | ns | 0.782 |
| Empty vector vs. 5 ng | 0.113 | -0.1932 to 0.4191 | ns | 0.8587 |
| Empty vector vs. 10 ng | 0.2092 | -0.09699 to 0.5153 | ns | 0.2944 |
| Empty vector vs. 25 ng | 0.1615 | -0.1447 to 0.4676 | ns | 0.5678 |
| Empty vector vs. 50 ng | 0.2256 | -0.08056 to 0.5318 | ns | 0.225 |
| 0 ng vs. 2 ng | -0.006333 | -0.3125 to 0.2998 | ns | >0.9999 |
| 0 ng vs. 5 ng | -0.0209 | -0.3271 to 0.2853 | ns | >0.9999 |
| 0 ng vs. 10 ng | 0.0753 | -0.2309 to 0.3815 | ns | 0.976 |
| 0 ng vs. 25 ng | 0.0276 | -0.2786 to 0.3338 | ns | >0.9999 |
| 0 ng vs. 50 ng | 0.09173 | -0.2144 to 0.3979 | ns | 0.94 |
| 2 ng vs. 5 ng | -0.01457 | -0.3207 to 0.2916 | ns | >0.9999 |
| 2 ng vs. 10 ng | 0.08163 | -0.2245 to 0.3878 | ns | 0.9647 |
| 2 ng vs. 25 ng | 0.03393 | -0.2722 to 0.3401 | ns | 0.9997 |

| | | | | |
|---|---|---|---|---|
| 2 ng vs. 50 ng | 0.09807 | -0.2081 to 0.4042 | ns | 0.9199 |
| 5 ng vs. 10 ng | 0.0962 | -0.2100 to 0.4024 | ns | 0.9262 |
| 5 ng vs. 25 ng | 0.0485 | -0.2577 to 0.3547 | ns | 0.9976 |
| 5 ng vs. 50 ng | 0.1126 | -0.1935 to 0.4188 | ns | 0.8603 |
| 10 ng vs. 25 ng | -0.0477 | -0.3539 to 0.2585 | ns | 0.9978 |
| 10 ng vs. 50 ng | 0.01643 | -0.2897 to 0.3226 | ns | >0.9999 |
| 25 ng vs. 50 ng | 0.06413 | -0.2420 to 0.3703 | ns | 0.9893 |

#### Figure 3D: RxCas13d mRNA (Random guide)

| Tukey's multiple comparisons test | Mean Diff. | 95.00% CI of diff. | Summary | Adjusted P Value |
|---|---|---|---|---|
| 0 ng vs. 2 ng | -0.03487 | -0.3933 to 0.3235 | ns | 0.9994 |
| 0 ng vs. 5 ng | 0.06207 | -0.2963 to 0.4205 | ns | 0.9904 |
| 0 ng vs. 10 ng | 0.01463 | -0.3438 to 0.3730 | ns | >0.9999 |
| 0 ng vs. 25 ng | 0.05043 | -0.3080 to 0.4088 | ns | 0.9963 |
| 0 ng vs. 50 ng | -0.207 | -0.5654 to 0.1514 | ns | 0.4257 |
| 2 ng vs. 5 ng | 0.09693 | -0.2615 to 0.4553 | ns | 0.937 |
| 2 ng vs. 10 ng | 0.0495 | -0.3089 to 0.4079 | ns | 0.9966 |
| 2 ng vs. 25 ng | 0.0853 | -0.2731 to 0.4437 | ns | 0.9622 |
| 2 ng vs. 50 ng | -0.1721 | -0.5305 to 0.1863 | ns | 0.606 |
| 5 ng vs. 10 ng | -0.04743 | -0.4058 to 0.3110 | ns | 0.9972 |
| 5 ng vs. 25 ng | -0.01163 | -0.3700 to 0.3468 | ns | >0.9999 |
| 5 ng vs. 50 ng | -0.269 | -0.6274 to 0.08936 | ns | 0.1922 |
| 10 ng vs. 25 ng | 0.0358 | -0.3226 to 0.3942 | ns | 0.9993 |
| 10 ng vs. 50 ng | -0.2216 | -0.5800 to 0.1368 | ns | 0.3588 |
| 25 ng vs. 50 ng | -0.2574 | -0.6158 to 0.1010 | ns | 0.2259 |

#### Figure 3D: mCherry mRNA (eGFP guide-2)

| Tukey's multiple comparisons test | Mean Diff. | 95.00% CI of diff. | Summary | Adjusted P Value |
|---|---|---|---|---|
| random vs. 0 ng | -0.0484 | -0.2995 to 0.2027 | ns | 0.9968 |
| random vs. 2 ng | 0.1311 | -0.1200 to 0.3822 | ns | 0.624 |
| random vs. 5 ng | 0.2857 | 0.03463 to 0.5368 | * | 0.0201 |
| random vs. 10 ng | 0.4173 | 0.1662 to 0.6684 | *** | 0.0006 |
| random vs. 25 ng | 0.547 | 0.2959 to 0.7981 | **** | <0.0001 |
| random vs. 50 ng | 0.5841 | 0.3330 to 0.8352 | **** | <0.0001 |
| 0 ng vs. 2 ng | 0.1795 | -0.07157 to 0.4306 | ns | 0.2722 |
| 0 ng vs. 5 ng | 0.3341 | 0.08303 to 0.5852 | ** | 0.0055 |
| 0 ng vs. 10 ng | 0.4657 | 0.2146 to 0.7168 | *** | 0.0002 |
| 0 ng vs. 25 ng | 0.5954 | 0.3443 to 0.8465 | **** | <0.0001 |
| 0 ng vs. 50 ng | 0.6325 | 0.3814 to 0.8836 | **** | <0.0001 |
| 2 ng vs. 5 ng | 0.1546 | -0.09650 to 0.4057 | ns | 0.4367 |
| 2 ng vs. 10 ng | 0.2862 | 0.03510 to 0.5373 | * | 0.0199 |
| 2 ng vs. 25 ng | 0.4159 | 0.1648 to 0.6670 | *** | 0.0006 |
| 2 ng vs. 50 ng | 0.4529 | 0.2018 to 0.7040 | *** | 0.0002 |
| 5 ng vs. 10 ng | 0.1316 | -0.1195 to 0.3827 | ns | 0.6202 |
| 5 ng vs. 25 ng | 0.2613 | 0.01016 to 0.5124 | * | 0.0384 |
| 5 ng vs. 50 ng | 0.2983 | 0.04723 to 0.5494 | * | 0.0144 |
| 10 ng vs. 25 ng | 0.1297 | -0.1214 to 0.3808 | ns | 0.636 |
| 10 ng vs. 50 ng | 0.1667 | -0.08437 to 0.4178 | ns | 0.3507 |
| 25 ng vs. 50 ng | 0.03707 | -0.2140 to 0.2882 | ns | 0.9994 |

#### Figure 3D: RxCas13d mRNA (eGFP guide-2)

| Tukey's multiple comparisons test | Mean Diff. | 95.00% CI of diff. | Summary | Adjusted P Value |
|---|---|---|---|---|
| random vs. 0 ng | -0.1144 | -0.5221 to 0.2932 | ns | 0.9552 |
| random vs. 2 ng | -0.1176 | -0.5253 to 0.2901 | ns | 0.9493 |
| random vs. 5 ng | 0.3192 | -0.08846 to 0.7269 | ns | 0.1762 |
| random vs. 10 ng | 0.4062 | -0.001491 to 0.8138 | ns | 0.0511 |
| random vs. 25 ng | 0.6398 | 0.2321 to 1.047 | ** | 0.0015 |
| random vs. 50 ng | 0.6586 | 0.2510 to 1.066 | ** | 0.0011 |
| 0 ng vs. 2 ng | -0.003167 | -0.4108 to 0.4045 | ns | >0.9999 |
| 0 ng vs. 5 ng | 0.4336 | 0.02598 to 0.8413 | * | 0.0338 |
| 0 ng vs. 10 ng | 0.5206 | 0.1129 to 0.9283 | ** | 0.009 |
| 0 ng vs. 25 ng | 0.7542 | 0.3465 to 1.162 | *** | 0.0003 |
| 0 ng vs. 50 ng | 0.7731 | 0.3654 to 1.181 | *** | 0.0002 |
| 2 ng vs. 5 ng | 0.4368 | 0.02914 to 0.8445 | * | 0.0323 |
| 2 ng vs. 10 ng | 0.5238 | 0.1161 to 0.9314 | ** | 0.0086 |
| 2 ng vs. 25 ng | 0.7574 | 0.3497 to 1.165 | *** | 0.0003 |
| 2 ng vs. 50 ng | 0.7762 | 0.3686 to 1.184 | *** | 0.0002 |
| 5 ng vs. 10 ng | 0.08697 | -0.3207 to 0.4946 | ns | 0.9882 |
| 5 ng vs. 25 ng | 0.3206 | -0.08709 to 0.7282 | ns | 0.173 |
| 5 ng vs. 50 ng | 0.3394 | -0.06822 to 0.7471 | ns | 0.1339 |
| 10 ng vs. 25 ng | 0.2336 | -0.1741 to 0.6413 | ns | 0.479 |
| 10 ng vs. 50 ng | 0.2525 | -0.1552 to 0.6601 | ns | 0.3952 |
| 25 ng vs. 50 ng | 0.01887 | -0.3888 to 0.4265 | ns | >0.9999 |

#### Figure 3D: mCherry mRNA (eGFP guide-3)

| Tukey's multiple comparisons test | Mean Diff. | 95.00% CI of diff. | Summary | Adjusted P Value |
|---|---|---|---|---|
| random vs. 0 ng | 0.07227 | -0.1940 to 0.3385 | ns | 0.9769 |
| random vs. 2 ng | 0.2298 | -0.03641 to 0.4961 | ns | 0.118 |
| random vs. 5 ng | 0.2594 | -0.006840 to 0.5256 | ns | 0.059 |
| random vs. 10 ng | 0.3708 | 0.1045 to 0.6370 | ** | 0.0036 |
| random vs. 25 ng | 0.5342 | 0.2680 to 0.8004 | **** | <0.0001 |
| random vs. 50 ng | 0.655 | 0.3887 to 0.9212 | **** | <0.0001 |
| 0 ng vs. 2 ng | 0.1576 | -0.1087 to 0.4238 | ns | 0.4826 |
| 0 ng vs. 5 ng | 0.1871 | -0.07911 to 0.4534 | ns | 0.2896 |
| 0 ng vs. 10 ng | 0.2985 | 0.03226 to 0.5647 | * | 0.0225 |
| 0 ng vs. 25 ng | 0.4619 | 0.1957 to 0.7282 | *** | 0.0004 |
| 0 ng vs. 50 ng | 0.5827 | 0.3165 to 0.8489 | **** | <0.0001 |
| 2 ng vs. 5 ng | 0.02957 | -0.2367 to 0.2958 | ns | >0.9999 |
| 2 ng vs. 10 ng | 0.1409 | -0.1253 to 0.4072 | ns | 0.6093 |
| 2 ng vs. 25 ng | 0.3044 | 0.03813 to 0.5706 | * | 0.0194 |
| 2 ng vs. 50 ng | 0.4251 | 0.1589 to 0.6914 | *** | 0.0009 |
| 5 ng vs. 10 ng | 0.1114 | -0.1549 to 0.3776 | ns | 0.8224 |
| 5 ng vs. 25 ng | 0.2748 | 0.008560 to 0.5410 | * | 0.0405 |
| 5 ng vs. 50 ng | 0.3956 | 0.1293 to 0.6618 | ** | 0.0019 |
| 10 ng vs. 25 ng | 0.1634 | -0.1028 to 0.4297 | ns | 0.4401 |
| 10 ng vs. 50 ng | 0.2842 | 0.01796 to 0.5504 | * | 0.0321 |
| 25 ng vs. 50 ng | 0.1208 | -0.1455 to 0.3870 | ns | 0.7602 |

#### Figure 3D: RxCas13d mRNA (eGFP guide-3)

| Tukey's multiple comparisons test | Mean Diff. | 95.00% CI of diff. | Summary | Adjusted P Value |
|---|---|---|---|---|

| | | | | |
|---|---|---|---|---|
| random vs. 0 ng | 0.2312 | -0.1010 to 0.5634 | ns | 0.2765 |
| random vs. 2 ng | 0.4231 | 0.09092 to 0.7553 | ** | 0.0092 |
| random vs. 5 ng | 0.3963 | 0.06406 to 0.7285 | * | 0.0152 |
| random vs. 10 ng | 0.4941 | 0.1619 to 0.8263 | ** | 0.0025 |
| random vs. 25 ng | 0.59 | 0.2578 to 0.9222 | *** | 0.0005 |
| random vs. 50 ng | 0.6841 | 0.3519 to 1.016 | **** | <0.0001 |
| 0 ng vs. 2 ng | 0.1919 | -0.1403 to 0.5241 | ns | 0.4703 |
| 0 ng vs. 5 ng | 0.165 | -0.1672 to 0.4972 | ns | 0.6289 |
| 0 ng vs. 10 ng | 0.2629 | -0.06934 to 0.5951 | ns | 0.1684 |
| 0 ng vs. 25 ng | 0.3587 | 0.02652 to 0.6909 | * | 0.0306 |
| 0 ng vs. 50 ng | 0.4528 | 0.1206 to 0.7850 | ** | 0.0053 |
| 2 ng vs. 5 ng | -0.02687 | -0.3591 to 0.3053 | ns | >0.9999 |
| 2 ng vs. 10 ng | 0.07097 | -0.2612 to 0.4032 | ns | 0.9881 |
| 2 ng vs. 25 ng | 0.1668 | -0.1654 to 0.4990 | ns | 0.6181 |
| 2 ng vs. 50 ng | 0.2609 | -0.07128 to 0.5931 | ns | 0.1738 |
| 5 ng vs. 10 ng | 0.09783 | -0.2344 to 0.4300 | ns | 0.9444 |
| 5 ng vs. 25 ng | 0.1937 | -0.1385 to 0.5259 | ns | 0.4602 |
| 5 ng vs. 50 ng | 0.2878 | -0.04441 to 0.6200 | ns | 0.1105 |
| 10 ng vs. 25 ng | 0.09587 | -0.2363 to 0.4281 | ns | 0.9492 |
| 10 ng vs. 50 ng | 0.19 | -0.1422 to 0.5222 | ns | 0.4813 |
| 25 ng vs. 50 ng | 0.0941 | -0.2381 to 0.4263 | ns | 0.9533 |

#### Figure 6D: FFLuc mRNA (FFLuc guide-1)

| Tukey's multiple comparisons test | Mean Diff. | 95.00% CI of diff. | Summary | Adjusted P Value |
|---|---|---|---|---|
| 150 ng vs. 50 ng | -0.05945 | -0.3725 to 0.2536 | ns | 0.9267 |
| 150 ng vs. 20 ng | -0.2568 | -0.5698 to 0.05633 | ns | 0.1127 |
| 50 ng vs. 20 ng | -0.1973 | -0.5104 to 0.1158 | ns | 0.2579 |

#### Figure 6D: FFLuc mRNA (FFLuc guide-4)

| Tukey's multiple comparisons test | Mean Diff. | 95.00% CI of diff. | Summary | Adjusted P Value |
|---|---|---|---|---|
| 150 ng vs. 50 ng | -0.1122 | -0.4961 to 0.2717 | ns | 0.7874 |
| 150 ng vs. 20 ng | -0.2991 | -0.6830 to 0.08484 | ns | 0.1354 |
| 50 ng vs. 20 ng | -0.1869 | -0.5708 to 0.1970 | ns | 0.4502 |

#### Figure 6E: nLuc mRNA (Random guide)

| Tukey's multiple comparisons test | Mean Diff. | 95.00% CI of diff. | Summary | Adjusted P Value |
|---|---|---|---|---|
| 0 ng vs. 20 ng | -0.08253 | -0.4622 to 0.2971 | ns | 0.9998 |
| 0 ng vs. 50 ng | -0.0099 | -0.3895 to 0.3697 | ns | >0.9999 |
| 0 ng vs. 150 ng | -0.04517 | -0.4248 to 0.3345 | ns | >0.9999 |
| 20 ng vs. 50 ng | 0.07263 | -0.3070 to 0.4523 | ns | >0.9999 |
| 20 ng vs. 150 ng | 0.03737 | -0.3423 to 0.4170 | ns | >0.9999 |
| 50 ng vs. 150 ng | -0.03527 | -0.4149 to 0.3444 | ns | >0.9999 |

#### Figure 6E: RxCas13d mRNA (Random guide)

| Tukey's multiple comparisons test | Mean Diff. | 95.00% CI of diff. | Summary | Adjusted P Value |
|---|---|---|---|---|
| 0 ng vs. 20 ng | -0.1916 | -0.5599 to 0.1767 | ns | 0.7624 |
| 0 ng vs. 50 ng | 0.07003 | -0.2983 to 0.4383 | ns | 0.9999 |
| 0 ng vs. 150 ng | -0.0031 | -0.3714 to 0.3652 | ns | >0.9999 |
| 20 ng vs. 50 ng | 0.2616 | -0.1067 to 0.6299 | ns | 0.3522 |

| | | | | |
|---|---|---|---|---|
| 20 ng vs. 150 ng | 0.1885 | -0.1798 to 0.5568 | ns | 0.7791 |
| 50 ng vs. 150 ng | -0.07313 | -0.4414 to 0.2952 | ns | 0.9998 |

#### Figure 6E: nLuc mRNA (FFLuc guide-1)

| Tukey's multiple comparisons test | Mean Diff. | 95.00% CI of diff. | Summary | Adjusted P Value |
|---|---|---|---|---|
| 0 ng vs. 20 ng | 0.2029 | -0.07829 to 0.4841 | ns | 0.1994 |
| 0 ng vs. 50 ng | 0.4755 | 0.1943 to 0.7567 | ** | 0.0017 |
| 0 ng vs. 150 ng | 0.6615 | 0.3803 to 0.9427 | *** | 0.0001 |
| 20 ng vs. 50 ng | 0.2726 | -0.008560 to 0.5538 | ns | 0.0584 |
| 20 ng vs. 150 ng | 0.4586 | 0.1774 to 0.7398 | ** | 0.0023 |
| 50 ng vs. 150 ng | 0.1859 | -0.09526 to 0.4671 | ns | 0.2629 |

#### Figure 6E: RxCas13d mRNA (FFLuc guide-1)

| Tukey's multiple comparisons test | Mean Diff. | 95.00% CI of diff. | Summary | Adjusted P Value |
|---|---|---|---|---|
| 0 ng vs. 20 ng | 0.143 | -0.1487 to 0.4346 | ns | 0.5217 |
| 0 ng vs. 50 ng | 0.5536 | 0.2619 to 0.8453 | *** | 0.0007 |
| 0 ng vs. 150 ng | 0.7727 | 0.4810 to 1.064 | **** | <0.0001 |
| 20 ng vs. 50 ng | 0.4106 | 0.1190 to 0.7023 | ** | 0.0065 |
| 20 ng vs. 150 ng | 0.6297 | 0.3381 to 0.9214 | *** | 0.0002 |
| 50 ng vs. 150 ng | 0.2191 | -0.07258 to 0.5108 | ns | 0.1733 |

#### Figure 6E: nLuc mRNA (FFLuc guide-4)

| Tukey's multiple comparisons test | Mean Diff. | 95.00% CI of diff. | Summary | Adjusted P Value |
|---|---|---|---|---|
| 0 ng vs. 20 ng | 0.09387 | -0.2573 to 0.4450 | ns | 0.898 |
| 0 ng vs. 50 ng | 0.3258 | -0.02533 to 0.6769 | ns | 0.0722 |
| 0 ng vs. 150 ng | 0.5286 | 0.1775 to 0.8798 | ** | 0.004 |
| 20 ng vs. 50 ng | 0.2319 | -0.1192 to 0.5831 | ns | 0.2637 |
| 20 ng vs. 150 ng | 0.4348 | 0.08363 to 0.7859 | * | 0.0149 |
| 50 ng vs. 150 ng | 0.2028 | -0.1483 to 0.5540 | ns | 0.3754 |

#### Figure 6E: RxCas13d mRNA (FFLuc guide-4)

| Tukey's multiple comparisons test | Mean Diff. | 95.00% CI of diff. | Summary | Adjusted P Value |
|---|---|---|---|---|
| 0 ng vs. 20 ng | 0.3321 | 0.09078 to 0.5734 | ** | 0.0075 |
| 0 ng vs. 50 ng | 0.6583 | 0.4170 to 0.8996 | **** | <0.0001 |
| 0 ng vs. 150 ng | 0.8895 | 0.6482 to 1.131 | **** | <0.0001 |
| 20 ng vs. 50 ng | 0.3263 | 0.08498 to 0.5676 | ** | 0.0085 |
| 20 ng vs. 150 ng | 0.5575 | 0.3162 to 0.7988 | *** | 0.0001 |
| 50 ng vs. 150 ng | 0.2312 | -0.01009 to 0.4725 | ns | 0.0619 |

**SUPPLEMENTARY METHODS**

The following *Drosophila* expression plasmids were each generated from the previously published **pUb 3xFLAG MCS** plasmid (Chen et al. (2012) *RNA*, **18**, 2148-2156.) using the three steps outlined below.

**pUb RxCas13d + RxCas13d guide RNA (Addgene #176303)**

**pUb dRxCas13d + RxCas13d guide RNA (Addgene #176304)**

**pUb PspCas13b + PspCas13b guide RNA (Addgene #176305)**

**pUb dPspCas13b + PspCas13b guide RNA (Addgene #176306)**

**pUb LwaCas13a + LwaCas13a guide RNA (Addgene #176307)**

The full plasmid sequence of **pUb 3xFLAG MCS** is as follows:

**GACGAAAGGGCCTCGTGATACGCCTATTTTTATAGGTTAA**TGTCATGATAATAATGGTTTCTTAGACGTCAGGTGGCACTTTTCGGGGAAATGTGCG
CGGAACCCCTATTTGTTTATTTTTCTAAATACATTCAAATATGTATCCGCTCATGAGACAATAACCCTGATAAATGCTTCAATAATATTGAAAAAGG
AAGAGTATGAGTATTCAACATTTCCGTGTCGCCCTTATTCCCTTTTTTGCGGCATTTTGCCTTCCTGTTTTTGCTCACCCAGAAACGCTGGTGAAAG
TAAAAGATGCTGAAGATCAGTTGGGTGCACGAGTGGGTTACATCGAACTGGATCTCAACAGCGGTAAGATCCTTGAGAGTTTTCGCCCCGAAGAACG
TTTTCCAATGATGAGCACTTTTAAAGTTCTGCTATGTGGCGCGGTATTATCCCGTATTGACGCCGGGCAAGAGCAACTCGGTCGCCGCATACACTAT
TCTCAGAATGACTTGGTTGAGTACTCACCAGTCACAGAAAAGCATCTTACGGATGGCATGACAGTAAGAGAATTATGCAGTGCTGCCATAACCATGA
GTGATAACACTGCGGCCAACTTACTTCTGACAACGATCGGAGGACCGAAGGAGCTAACCGCTTTTTTGCACAACATGGGGGATCATGTAACTCGCCT
TGATCGTTGGGAACCGGAGCTGAATGAAGCCATACCAAACGACGAGCGTGACACCACGATGCCTGTAGCAATGGCAACAACGTTGCGCAAACTATTA
ACTGGCGAACTACTTACTCTAGCTTCCCGGCAACAATTAATAGACTGGATGGAGGCGGATAAAGTTGCAGGACCACTTCTGCGCTCGGCCCTTCCGG
CTGGCTGGTTTATTGCTGATAAATCTGGAGCCGGTGAGCGTGGGTCTCGCGGTATCATTGCAGCACTGGGGCCAGATGGTAAGCCCTCCCGTATCGT
AGTTATCTACACGACGGGGAGTCAGGCAACTATGGATGAACGAAATAGACAGATCGCTGAGATAGGTGCCTCACTGATTAAGCATTGGTAACTGTCA
GACCAAGTTTACTCATATATACTTTAGATTGATTTAAAACTTCATTTTTAATTTAAAAGGATCTAGGTGAAGATCCTTTTTGATAATCTCATGACCA
AAATCCCTTAACGTGAGTTTTCGTTCCACTGAGCGTCAGACCCCGTAGAAAAGATCAAAGGATCTTCTTGAGATCCTTTTTTTCTGCGCGTAATCTG
CTGCTTGCAAACAAAAAAACCACCGCTACCAGCGGTGGTTTGTTTGCCGGATCAAGAGCTACCAACTCTTTTTCCGAAGGTAACTGGCTTCAGCAGA
GCGCAGATACCAAATACTGTTCTTCTAGTGTAGCCGTAGTTAGGCCACCACTTCAAGAACTCTGTAGCACCGCCTACATACCTCGCTCTGCTAATCC
TGTTACCAGTGGCTGCTGCCAGTGGCGATAAGTCGTGTCTTACCGGGTTGGACTCAAGACGATAGTTACCGGATAAGGCGCAGCGGTCGGGCTGAAC
GGGGGGTTCGTGCACACAGCCCAGCTTGGAGCGAACGACCTACACCGAACTGAGATACCTACAGCGTGAGCTATGAGAAAGCGCCACGCTTCCCGAA
GGGAGAAAGGCGGACAGGTATCCGGTAAGCGGCAGGGTCGGAACAGGAGAGCGCACGAGGGAGCTTCCAGGGGGAAACGCCTGGTATCTTTATAGTC
CTGTCGGGTTTCGCCACCTCTGACTTGAGCGTCGATTTTTGTGATGCTCGTCAGGGGGGCGGAGCCTATGGAAAAACGCCAGCAACGCGGCCTTTTT
ACGGTTCCTGGCCTTTTGCTGGCCTTTTGCTCACATGTTCTTTCCTGCGTTATCCCCTGATTCTGTGGATAACCGTATTACCGCCTTTGAGTGAGCT
GATACCGCTCGCCGCAGCCGAACGACCGAGCGCAGCGAGTCAGTGAGCGAGGAAGCGGAAGAGCGCCCAATACGCAAACCGCCTCTCCCCGCGCGTT
GGCCGATTCATTAATGCAGCTGGCACGACAGGTTTCCCGACTGGAAAGCGGGCAGTGAGCGCAACGCAATTAATGTGAGTTAGCTCACTCATTAGGC
ACCCCAGGCTTTACACTTTATGCTTCCGGCTCGTATGTTGTGTGGAATTGTGAGCGGATAACAATTTCACACAGGAAACAGCTATGACCATGATTAC
GCCAAAGCTTGTCGCCGGAACGCAGCGACAGAGATTCCAATGTGTCCGTATCTTTCAGGCTTTTGCCCTTCAGTTCCAGACGAAGCGACTGGCGATT
CGCGTGTGGGGTCTGCTTCAGGGTCTTGTGAATTAGGGCGCGCAGATCGCCGATGGGCGTGGCGCCGGAGGGCACCTTCACCTTGCCGTACGGCTTG
CTGTTCTTCGCGTTCAAAATCTCCAGCTCCATTTTGCTTTCGGTGCGCTTGCAATCAGTACTGTCCAAAATCGAAAATCGCCGAACCGTAGTGTGAC
CGTGCGGGGCTCTGCGAAAATAAACTTTTTTAGGTATATGGCCACACACGGGGAAAGCACAGTGGATTATATGTTTTAATATTATAATATGCAGGTT
TTCATTACTTATCCAGATGTAAGCCCACTTAAAGCGATTTAACAATTATTTGCCGAAAGAGTATAAACAAATTTCACTTAAAAATGGATTAAGAAAA
GCTAGCTTGTGTAAGATTATGCGCAGCGTTGCCAGATAGCTCCATTTAAAACACTTCAAAAACAATAAGTTTTGAAAATATATACATAAATAGCAGT
CGTTGCCGCAACGCTCAACACATCACACTTTTAAAACACCCTTTACCTACACAGAATTACTTTTTAAATTTCCAGTCAAGCTGCGAGTTTCAAAATT
ATAGCCGGTAGAGAAGACAGTGCTATTTCAAAAGCAAACTAACAAGGGTCTTAAATTCCAAAACACCAATCCTAACAAGCCTTGGACTTTTGTAAGT
TTAGATCAAAGGTGGCATTGCATTCAATGTCATGGTAAGAAGTAGGTCGTCTAGGTAGAAATCCTCATTCAGCCGGTCAAGTCAGTACGAGAAAGGT
CTCAATTTGAAATTGTCTTAAAAATATTTTATTGTTTTGTACTGTGGTGAGTTTAAACGAAAAACACAAAAAAAAAGTGATACACAGAAATCATAAA
AAATTTTAATACAAGGTATTCGTACGTATCAAAAACATTTCGGCACAATTTTTTTTCTCTGTACTAAAGTGTTACGAACACTACGGTATTTTTTAGT
GATTTTCAACGGACACCGAAGGTATATAAACAGCGTTCGCGAACGGTCGCCTTCAAAACCAATTGACATTTGCAGCAGCAAGTACAAGTAGAAAGTA
AAGCGCAATCAGCGAAAAATTTATACTTAATTGTTGGTGATTAAAGTACAATTAAAAGAACATTCTCGAAAGTCACAAGAAACGTAAGTTTTTAACT
CGCTGTTACCAATTAGTAATAAGAGCAACAAGACGTTGAGTAATTTCAAGAAAAACTGCATTTCAAGGTCTTTGTTCGGCCATTTTTTTTTATTCAA
CGCTCTACGTAATTACAAAATAAGAAATTGGCAGCCACGCATCTTGTTTTCCCAATGAATTGGCATCAAAACGCAAACAAATCTATAAATAAAACTT
GCGTGTTGATTTTCGCCAAGATTTATTGGCAAATTGTGAAATTCGCAGTGACGCATTTGAAAATTCGAGAAATCACGAACGCACTCGATCGAGCATT
TGTGTGCATGTTATTAGTTAGTTGTTTAGTTAATTGAAGTATTTTACCAACGAAATCCACTTATTTTTAGCTGAAATAGAGTAGGTTGCTTAAACAA
AGCCACGTCTGAAAATTTCTTATTGCTTGTAGTTGTGACGTCACCATATACACACAAAATAATGTGTATGCATGCATTTCAGCTGTGTATATATACA

TGCACACACTCGCAACACGAAAACGATGACGAAGCAACGGAACAAAGGTTTCTCAACTACCCTTTGTTCCCTGTTTCTTCGCTTTCCTTTGTTCCAA
TATTCGTAGAGGGTTAATAGGGGTTTCTCAACAAAGTTGGCGTCGATAAATAAGTTTCCCATTTTTATTCCCCAGCCAGGAAGTTAGTTTCAATAGT
TTTGTAATTTCAACGAAACTCATTTGATTTCGTACTAATTTTCCACATCTCTATTTTCTTCCCGCAGAATAATCCAAACTGCAGGTCGACTCTAGCT
AGAGGAAGCTTATGGACTACAAAGACCATGACGGTGATTATAAAGATCATGATATCGATTACAAGGATGACGATGACAAGGATCCACTAGTCCAGTG
TGGTGGAATTCTGCAGATATCCAGCACAGTGGCGGCCGCTCGAGTCTAGAGGGCCCGCGGTTCGAAGGTAAGCCTATCCCTAACCCTCTCCTCGGTC
TCGATTCTACGCGTACCGGTCATCATCACCATCACCATTGAGTTTATCTGACTAAATCTTAGTTTGTATTGTCATGTTTTAATACAATATGTTATGT
TTAAATATGTTTTTAATAAATTTTATAAAATAATTTCAACTTTTATTGTAACAACATTGTCCATTTACACACTCCTTTCAAGCGCGTGGGATCGATG
CTCACTCAAAGGCGGTAATACGGTTATCCACAGAATCAGGGGATAACGCAGGAAAGAACATGTGAGCCATATGGGCCCATGTGAGCCATATGGTGCA
CTCTCAGTACAATCTGCTCTGATGCCGCATAGTTAAGCCAGCCCCGACACCCGCCAACACCCGCTGACGCGCCCTGACGGGCTTGTCTGCTCCCGGC
ATCCGCTTACAGACAAGCTG**TGACCGTCTCCGGGAGCTGCATGTGTCAGAGGTTTTCACCGTCATCACCGAAACGCGCGA**

pUbi-p63e (pUb) promoter

CGTCTC: BsmBI site

CATATG: NdeI site

GAACGTTTTC: XmnI site

*Step 1:*

The BsmBI restriction site (yellow) and surrounding nucleotides (in bold above) were first removed to generate the **pUb 3xFLAG MCS (No BsmBI)** plasmid. This was done by cutting **pUb 3xFLAG MCS** with NdeI and XmnI and inserting the following sequence between the NdeI and XmnI sites.

GGCCCATGTGAGCCATATGGTGCACTCTCAGTACAATCTGCTCTGATGCCGCATAGTTAAGCCAGCCCCGACACCCGCCAACACCCGCTGACGCGCC
CTGACGGGCTTGTCTGCTCCCGGCATCCGCTTACAGACAAGCTGTGTCATGATAATAATGGTTTCTTAGACGTCAGGTGGCACTTTTCGGGGAAATG
TGCGCGGAACCCCTATTTGTTTATTTTTCTAAATACATTCAAATATGTATCCGCTCATGAGACAATAACCCTGATAAATGCTTCAATAATATTGAAA
AAGGAAGAGTATGAGTATTCAACATTTCCGTGTCGCCCTTATTCCCTTTTTTGCGGCATTTTGCCTTCCTGTTTTTGCTCACCCAGAAACGCTGGTG
AAAGTAAAAGATGCTGAAGATCAGTTGGGTGCACGAGTGGGTTACATCGAACTGGATCTCAACAGCGGTAAGATCCTTGAGAGTTTTCGCCCCGAA

*Step 2:*

A guide RNA sequence containing BsmBI restriction sites that is driven by the snRNA:U6:96Ab promoter was inserted into the NdeI site.

**pUb 3xFLAG MCS (No BsmBI) + RxCas13d guide RNA** was made by inserting the following sequence into the NdeI site in **pUb 3xFLAG MCS (No BsmBI)**:

gttcgacttgcagcctgaaatacggcacgagtaggaaaagccgagtcaaatgccgaatgcagagtctcattacagcacaatcaactcaagaaaaact
cgacactttttaccatttgcacttaaatcctttttattcgttatgtatacttttttggtccctaaccaaaacaaaaccaaactctcttagtcgt
gcctctatatttaaaactatcaatttattatagtcaataaatcgaactgtgttttcaacaaacgaacaataggacactttgattctaaaggaaattt
tgaaaatcttaagcagagggttcttaagaccatttgccaattcttataattctcaactgctctttcctgatgttgatcatttatataggtatgtttt
cctcaatacttcgaacccctaccaactggtcggggtttgaaacgagacggttgtatttgtacgtctctttttt

snRNA:U6:96Ab Promoter
Terminator
RxCas13d guide RNA scaffold
BsmBI sites

**pUb 3xFLAG MCS (No BsmBI) + PspCas13b guide RNA** was made by inserting the following sequence into the NdeI site in **pUb 3xFLAG MCS (No BsmBI)**:

Gttcgacttgcagcctgaaatacggcacgagtaggaaaagccgagtcaaatgccgaatgcagagtctcattacagcacaatcaactcaagaaaaact
cgacactttttaccatttgcacttaaatcctttttattcgttatgtatacttttttggtccctaaccaaaacaaaaccaaactctcttagtcgt
gcctctatatttaaaactatcaatttattatagtcaataaatcgaactgtgttttcaacaaacgaacaataggacactttgattctaaaggaaattt
tgaaaatcttaagcagagggttcttaagaccatttgccaattcttataattctcaactgctctttcctgatgttgatcatttatataggtatgtttt
cctcaatacttcggagacggttgtatttgtagttgtacgtctcAgttgtggaaggtccagttttgaggggctattacaacttttttt

snRNA:U6:96Ab Promoter
Terminator
PspCas13b guide RNA scaffold
BsmBI sites

**pUb 3xFLAG MCS (No BsmBI) + LwaCas13a guide RNA** was made by inserting the following sequence into the NdeI site in **pUb 3xFLAG MCS (No BsmBI)**:

Gttcgacttgcagcctgaaatacggcacgagtaggaaaagccgagtcaaatgccgaatgcagagtctcattacagcacaatcaactcaagaaaaact
cgacactttttttaccatttgcacttaaatcctttttttattcgttatgtatacttttttttggtccctaaccaaaacaaaaccaaactctcttagtcgt
gcctctatatttaaaactatcaatttattatagtcaataaatcgaactgtgttttcaacaaacgaacaataggacactttgattctaaaggaaattt
tgaaaatcttaagcagagggttcttaagaccatttgccaattcttataattctcaactgctctttcctgatgttgatcatttatataggtatgtttt
cctcaatacttcgGATTTAGACTACCCCAAAAACGAAGGGGACTAAAACgagacggttgtatttgtagttgcgtctcTTTTTTT

snRNA:U6:96Ab Promoter
Terminator
LwaCas13a guide RNA scaffold
BsmBI sites

*Step 3:*

The indicated Cas13 ORF was inserted downstream of the Ubi-p63e promoter.

**pUb RxCas13d + RxCas13d guide RNA** was made by inserting the RxCas13d ORF between SalI and AgeI in **pUb 3xFLAG MCS (No BsmBI) + RxCas13d guide RNA**:

aagaccatgattgagaagaagaaatcctttgccaaagggatgggcgtgaagagcactctggtgagcggcagcaaggtgtacatgaccacctttgcag
aggggagtgatgcccggctggagaagatcgtggaaggggacagcatccgctctgtgaatgaaggagaagctttcagtgcagaaatggctgacaagaa
tgctgggtacaaaatcgggaacgccaagttcagccaccccaagggatatgcagtggttgctaacaatcctctctacacaggacctgtgcagcaggac
atgctggggctgaaggagactttggagaagcggtatttcggggagtctgcagatggaaatgacaatatctgcattcaggtgattcataatattcttg
atattgaaaagatcctggctgagtacatcacaaatgctgcatatgctgttaataacatctcaggcttggacaaggatattattgggtttgggaagtt
ttccacagtgtatacctacgatgaatttaaagaccccgagcaccacagagctgccttcaacaataatgacaaattgatcaatgccatcaaggctcag
tatgatgaatttgacaacttcctggacaacccaaggctgggctattttggccaggccttcttcagcaaagagggccgcaactacatcattaattacg
ggaatgaatgttatgatatcctcgccctgctctctggcctgaggcactgggtggtccacaacaacgaggaagaaagcaggatttctaggacctggct
gtacaatcttgacaagaatctggacaatgaatatatctccaccctcaactatttgtatgaccggatcaccaacgaactcacaaacagcttttctaag
aactcagccgcaaatgtgaactacattgcagagacactgggcatcaaccccgccgagtttgctgaacagtacttccggttctccattatgaaggagc
agaagaatttgggttttaacattactaaacttagggaggtcatgctcgacaggaaagatatgtctgagatccggaagaaccacaaggtcttcgattc
catcaggacaaaagtctacaccatgatggactttgttatttaccgctattacattgaagaagatgctaaagtggccgccgctaataagagcctgcca
gataatgagaaatctctcagcgagaaggacatctttgtgatcaacctgcggggctccttcaatgatgatcagaaggatgcactgtactacgatgaag
ccaaccgcatctggagaaagctggagaacatcatgcacaatataaaggaattccgcggcaataagacacgggagtacaagaagaaagacgccccacg
cctgcccagaattctgcctgctggccgggatgtctctgccttttcaaaactcatgtatgccttgacaatgtttctggatggcaaagaaatcaatgat
ttgctgacaaccctgatcaacaaatttgacaacatccagagcttcctcaaggtgatgcccctgattggagtaaatgcaaaatttgtcgaggagtatg
ccttcttcaaggactctgccaagattgccgatgagctgcgcctcattaagtcattcgcccgcatgggagagcccatcgcagacgcaagaagagccat
gtacatcgatgccattcgcattttgggcaccaatctctcctatgacgagctcaaagctctggccgacaccttctctttggatgaaaatggcaacaag
cttaagaaggggaaacatggcatgagaaatttcatcatcaacaacgtcatttccaacaagcgcttccactatctcatcagatatggagaccctgccc
acctgcatgaaattgctaagaatgaggctgtggtgaaattcgtgctgggaaggatcgccgacattcaaaagaagcaggggcagaatggaaagaacca
gatcgaccgctactatgaaacctgtatcggaaaagataagggcaagtctgtcagtgagaaagtggacgccctcaccaagatcatcactggaatgaac
tatgaccagttcgacaagaaaagaagtgtgatcgaggacacaggccgggagaatgccgagagagagaagttcaagaaaattatcagcctctatctga
ccgtcatctaccacatcctgaagaacattgtcaacattaatgccaggtacgtgattggctttcactgcgttgaaagagatgcccagctttataagga
gaaaggctacgacattaacttgaagaaactggaggagaaaggatttagcagcgtgaccaaactgtgtgccggcattgacgagaccgcccctgataaa
cggaaagatgtggagaaagagatggccgagcgggccaaggagagcatcgattctctggaaagtgccaaccctaagctctacgccaattatattaaat
attctgatgagaagaaagcagaagagttcacccggcagattaacagggaaaaggccaaaacagccctgaacgcctacctgagaaacaccaagtggaa
tgtcatcattagagaagacctgcttcggattgataacaagacctgcaccctcttcaggaacaaagccgtgcacttggaagtggctcgctatgtgcat
gcctatatcaatgacatcgctgaggtgaactcctacttccagctgtatcactacatcatgcagagaatcattatgaatgaaaggtatgagaaatcat
ctggtaaagtatctgaatactttgatgctgtcaatgatgagaagaagtacaacgacagactcctgaagctgctgtgcgtgccctttggatactgcat
ccccaggtttaagaacctcagtattgaggccctctttgataggaatgaggcagcaaagtttgataaagaaaagaagaaagtttctggaaacagtgga
agtggcgcagctgcttacccctacgacgtgcctgactacgcctga

**pUb dRxCas13d + RxCas13d guide RNA** was made by inserting the dRxCas13d ORF (catalytic dead mutations are noted in red) between SalI and AgeI in **pUb 3xFLAG MCS (No BsmBI) + RxCas13d guide RNA**:

aagaccatgattgagaagaagaaatcctttgccaaagggatgggcgtgaagagcactctggtgagcggcagcaaggtgtacatgaccacctttgcag
aggggagtgatgcccggctggagaagatcgtggaaggggacagcatccgctctgtgaatgaaggagaagctttcagtgcagaaatggctgacaagaa
tgctgggtacaaaatcgggaacgccaagttcagccaccccaagggatatgcagtggttgctaacaatcctctctacacaggacctgtgcagcaggac
atgctggggctgaaggagactttggagaagcggtatttcggggagtctgcagatggaaatgacaatatctgcattcaggtgattcataatattcttg
atattgaaaagatcctggctgagtacatcacaaatgctgcatatgctgttaataacatctcaggcttggacaaggatattattgggtttgggaagtt
ttccacagtgtatacctacgatgaatttaaagaccccgagcaccacagagctgccttcaacaataatgacaaattgatcaatgccatcaaggctcag
tatgatgaatttgacaacttcctggacaacccaaggctgggctattttggccaggccttcttcagcaaagagggccgcaactacatcattaattacg
ggaatgaatgttatgatatcctcgccctgctctctggcctg**gcc**cactgggtggtc**gcc**aacaacgaggaagaaagcaggatttctaggacctggct
gtacaatcttgacaagaatctggacaatgaatatatctccaccctcaactatttgtatgaccggatcaccaacgaactcacaaacagcttttctaag
aactcagccgcaaatgtgaactacattgcagagacactgggcatcaaccccgccgagtttgctgaacagtacttccggttctccattatgaaggagc
agaagaatttgggttttaacattactaaacttagggaggtcatgctcgacaggaaagatatgtctgagatccggaagaaccacaaggtcttcgattc
catcaggacaaaagtctacaccatgatggactttgttatttaccgctattacattgaagaagatgctaaagtggccgccgctaataagagcctgcca
gataatgagaaatctctcagcgagaaggacatctttgtgatcaacctgcggggctccttcaatgatgatcagaaggatgcactgtactacgatgaag
ccaaccgcatctggagaaagctggagaacatcatgcacaatataaaggaattccgcggcaataagacacgggagtacaagaagaaagacgccccacg
cctgcccagaattctgcctgctggccgggatgtctctgccttttcaaaactcatgtatgccttgacaatgtttctggatggcaaagaaatcaatgat
ttgctgacaaccctgatcaacaaatttgacaacatccagagcttcctcaaggtgatgcccctgattggagtaaatgcaaaatttgtcgaggagtatg
ccttcttcaaggactctgccaagattgccgatgagctgcgcctcattaagtcattcgcccgcatgggagagcccatcgcagacgcaagaagagccat
gtacatcgatgccattcgcattttgggcaccaatctctcctatgacgagctcaaagctctggccgacaccttctctttggatgaaaatggcaacaag
cttaagaaggggaaacatggcatgagaaatttcatcatcaacaacgtcatttccaacaagcgcttccactatctcatcagatatggagaccctgccc
acctgcatgaaattgctaagaatgaggctgtggtgaaattcgtgctgggaaggatcgccgacattcaaaagaagcaggggcagaatggaaagaacca
gatcgaccgctactatgaaacctgtatcggaaaagataagggcaagtctgtcagtgagaaagtggacgccctcaccaagatcatcactggaatgaac
tatgaccagttcgacaagaaaagaagtgtgatcgaggacacaggccgggagaatgccgagagagagagaagttcaagaaaattatcagcctctatctga
ccgtcatctaccacatcctgaagaacattgtcaacattaatgccaggtacgtgattggctttcactgcgttgaaagagatgcccagctttataagga
gaaaggctacgacattaacttgaagaaactggaggagaaaggatttagcagcgtgaccaaactgtgtgccggcattgacgagaccgcccctgataaa
cggaaagatgtggagaaagagatggccgagcgggccaaggagagcatcgattctctggaaagtgccaaccctaagctctacgccaattatattaaat
attctgatgagaagaaagcagaagagttcacccggcagattaacagggaaaaggccaaaacagccctgaacgcctacctgagaaacaccaagtggaa
tgtcatcattagagaagacctgcttcggattgataacaagacctgcaccctcttc**gcc**aacaaagccgt**ggc**cttggaagtggctcgctatgtgcat
gcctatatcaatgacatcgctgaggtgaactcctacttccagctgtatcactacatcatgcagagaatcattatgaatgaaaggtatgagaaatcat
ctggtaaagtatctgaatactttgatgctgtcaatgatgagaagaagtacaacgacagactcctgaagctgctgtgcgtgccctttggatactgcat
ccccaggtttaagaacctcagtattgaggccctctttgataggaatgaggcagcaaagtttgataaagaaaagaagaaagtttctggaaacagtgga
agtggcgcagctgcttacccctacgacgtgcctgactacgcctga

**pUb PspCas13b + PspCas13b guide RNA** was made by inserting the PspCas13b ORF between SalI and MluI in **pUb 3xFLAG MCS (No BsmBI) + PspCas13b guide RNA**:

aagaccatggattacaaagacgatgacgataagGTTAACGGTACCGAGCTCCCCGGGTTAATTAAAggtggatctaacatccccgctctggtggaaa
accagaagaagtactttggcacctacagcgtgatggccatgctgaacgctcagaccgtgctggaccacatccagaaggtggccgatattgagggcga
gcagaacgagaacaacgagaatctgtggtttcaccccgtgatgagccacctgtacaacgccaagaacggctacgacaagcagcccgagaaaaccatg
ttcatcatcgagcggctgcagagctacttcccattcctgaagatcatggccgagaaccagagagagtacagcaacggcaagtacaagcagaaccgcg
tggaagtgaacagcaacgacatcttcgaggtgctgaagcgcgccttcggcgtgctgaagatgtacagggacctgaccaaccactacaagacctacga
ggaaaagctgaacgacggctgcgagttcctgaccagcacagagcaacctctgagcggcatgatcaacaactactacacagtggccctgcggaacatg
aacgagagatacggctacaagacagaggacctggccttcatccaggacaagcggttcaagttcgtgaaggacgcctacggcaagaaaaagtcccaag
tgaataccggattcttcctgagcctgcaggactacaacggcgacacacagaagaagctgcacctgagcggagtgggaatcgccctgctgatctgcct
gttcctggacaagcagtacatcaacatctttctgagcaggctgcccatcttctccagctacaatgcccagagcgaggaacggcggatcatcatcaga
tccttcggcatcaacagcatcaagctgcccaaggaccggatccacagcgagaagtccaacaagagcgtggccatggatatgctcaacgaagtgaagc
ggtgccccgacgagctgttcacaacactgtctgccgagaagcagtcccggttcagaatcatcagcgacgaccacaatgaagtgctgatgaagcggag
cagcgacagattcgtgcctctgctgctgcagtatatcgattacggcaagctgttcgaccacatcaggttccacgtgaacatgggcaagctgagatac
ctgctgaaggccgacaagacctgcatcgacggccagaccagagtcagagtgatcgagcagcccctgaacggcttcggcagactggaagaggccgaga
caatgcggaagcaagagaacggcaccttcggcaacagcggcatccggatcagagacttcgagaacatgaagcgggacgacgccaatcctgccaacta
tccctacatcgtggacacctacacacactacatcctggaaaacaacaaggtcgagatgtttatcaacgacaaagaggacagcgccccactgctgccc
gtgatcgaggatgatagatacgtggtcaagacaatccccagctgccggatgagcaccctggaaattccagccatggccttccacatgtttctgttcg
gcagcaagaaaaccgagaagctgatcgtggacgtgcacaaccggtacaagagactgttccaggccatgcagaaagaagaagtgaccgccgagaatat
cgccagcttcggaatcgccgagagcgacctgcctcagaagatcctggatctgatcagcggcaatgcccacggcaaggatgtggacgccttcatcaga
ctgaccgtggacgacatgctgaccgacaccgagcggagaatcaagagattcaaggacgaccggaagtccattcggagcgccgacaacaagatgggaa
agagaggcttcaagcagatctccacaggcaagctggccgacttcctggccaaggacatcgtgctgtttcagcccagcgtgaacgatggcgagaacaa
gatcaccggcctgaactaccggatcatgcagagcgccattgccgtgtacgatagcggcgacgattacgaggccaagcagcagttcaagctgatgttc
gagaaggcccggctgatcggcaagggcacaacagagcctcatccatttctgtacaaggtgttcgcccgcagcatccccgccaatgccgtcgagttct
acgagcgctacctgatcgagcggaagttctacctgaccggcctgtccaacgagatcaagaaaggcaacagagtggatgtgcccttcatccggcggga
ccagaacaagtggaaaacacccgccatgaagaccctgggcagaatctacagcgaggatctgcccgtggaactgcccagacagatgttcgacaatgag

atcaagtcccacctgaagtccctgccacagatggaaggcatcgacttcaacaatgccaacgtgacctatctgatcgccgagtacatgaagagagtgc
tggacgacgacttccagaccttctaccagtggaaccgcaactaccggtacatggacatgcttaagggcgagtacgacagaaagggctccctgcagca
ctgcttcaccagcgtggaagagagagaaggcctctggaaagagcgggcctccagaacagagcggtacagaaagcaggccagcaacaagatccgcagc
aaccggcagatgagaaacgccagcagcgaagagatcgagacaatcctggataagcggctgagcaacagccggaacgagtaccagaaaagcgagaaag
tgatccggcgctacagagtgcaggatgccctgctgtttctgctggccaaaaagaccctgaccgaactggccgatttcgacggcgagaggttcaaact
gaaagaaatcatgcccgacgccgagaagggaatcctgagcgagatcatgcccatgagcttcaccttcgagaaaggcggcaagaagtacaccatcacc
agcgagggcatgaagctgaagaactacggcgacttctttgtgctggctagcgacaagaggatcggcaacctgctggaactcgtgggcagcgacatcg
tgtccaaagaggatatcatggaagagttcaacaaatacgaccagtgcaggcccgagatcagctccatcgtgttcaacctggaaaagtgggccttcga
cacataccccgagctgtctgccagagtggaccgggaagagaaggtggacttcaagagcatcctgaaaatcctgctgaacaacaagaacatcaacaaa
gagcagagcgacatcctgcggaagatccggaacgccttcgatcacaacaattaccccgacaaaggcgtggtggaaatcaaggccctgcctgagatcg
ccatgagcatcaagaaggcctttggggagtacgccatcatgaagggaagtggctacccctacgacgtgcctgactacgcctga

**pUb dPspCas13b + PspCas13b guide RNA** was made by inserting the dPspCas13b ORF (catalytic dead mutations are noted in red) between SalI and MluI in **pUb 3xFLAG MCS (No BsmBI) + PspCas13b guide RNA**:

aagaccatggattacaaagacgatgacgataagGTTAACGGTACCGAGCTCCCCGGGTTAATTAAAggtggatctAACATCCCCGCTCTGGTGGAAA
ACCAGAAGAAGTACTTTGGCACCTACAGCGTGATGGCCATGCTGAACGCTCAGACCGTGCTGGACCACATCCAGAAGGTGGCCGATATTGAGGGCGA
GCAGAACGAGAACAACGAGAATCTGTGGTTTCACCCCGTGATGAGCCACCTGTACAACGCCAAGAACGGCTACGACAAGCAGCCCGAGAAAACCATG
TTCATCATCGAGCGGCTGCAGAGCTACTTCCCATTCCTGAAGATCATGGCCGAGAACCAGAGAGAGTACAGCAACGGCAAGTACAAGCAGAACCGCG
TGGAAGTGAACAGCAACGACATCTTCGAGGTGCTGAAGCGCGCCTTCGGCGTGCTGAAGATGTACAGGGACCTGACCAAC**GCA**TACAAGACCTACGA
GGAAAAGCTGAACGACGGCTGCGAGTTCCTGACCAGCACAGAGCAACCTCTGAGCGGCATGATCAACAACTACTACACAGTGGCCCTGCGGAACATG
AACGAGAGATACGGCTACAAGACAGAGGACCTGGCCTTCATCCAGGACAAGCGGTTCAAGTTCGTGAAGGACGCCTACGGCAAGAAAAAGTCCCAAG
TGAATACCGGATTCTTCCTGAGCCTGCAGGACTACAACGGCGACACACAGAAGAAGCTGCACCTGAGCGGAGTGGGAATCGCCCTGCTGATCTGCCT
GTTCCTGGACAAGCAGTACATCAACATCTTTCTGAGCAGGCTGCCCATCTTCTCCAGCTACAATGCCCAGAGCGAGGAACGGCGGATCATCATCAGA
TCCTTCGGCATCAACAGCATCAAGCTGCCCAAGGACCGGATCCACAGCGAGAAGTCCAACAAGAGCGTGGCCATGGATATGCTCAACGAAGTGAAGC
GGTGCCCCGACGAGCTGTTCACAACACTGTCTGCCGAGAAGCAGTCCCGGTTCAGAATCATCAGCGACGACCACAATGAAGTGCTGATGAAGCGGAG
CAGCGACAGATTCGTGCCTCTGCTGCTGCAGTATATCGATTACGGCAAGCTGTTCGACCACATCAGGTTCCACGTGAACATGGGCAAGCTGAGATAC
CTGCTGAAGGCCGACAAGACCTGCATCGACGGCCAGACCAGAGTCAGAGTGATCGAGCAGCCCCTGAACGGCTTCGGCAGACTGGAAGAGGCCGAGA
CAATGCGGAAGCAAGAGAACGGCACCTTCGGCAACAGCGGCATCCGGATCAGAGACTTCGAGAACATGAAGCGGGACGACGCCAATCCTGCCAACTA
TCCCTACATCGTGGACACCTACACACACTACATCCTGGAAAACAACAAGGTCGAGATGTTTATCAACGACAAAGAGGACAGCGCCCCACTGCTGCCC
GTGATCGAGGATGATAGATACGTGGTCAAGACAATCCCCAGCTGCCGGATGAGCACCCTGGAAATTCCAGCCATGGCCTTCCACATGTTTCTGTTCG
GCAGCAAGAAAACCGAGAAGCTGATCGTGGACGTGCACAACCGGTACAAGAGACTGTTCCAGGCCATGCAGAAAGAAGAAGTGACCGCCGAGAATAT
CGCCAGCTTCGGAATCGCCGAGAGCGACCTGCCTCAGAAGATCCTGGATCTGATCAGCGGCAATGCCCACGGCAAGGATGTGGACGCCTTCATCAGA
CTGACCGTGGACGACATGCTGACCGACACCGAGCGGAGAATCAAGAGATTCAAGGACGACCGGAAGTCCATTCGGAGCGCCGACAACAAGATGGGAA
AGAGAGGCTTCAAGCAGATCTCCACAGGCAAGCTGGCCGACTTCCTGGCCAAGGACATCGTGCTGTTTCAGCCCAGCGTGAACGATGGCGAGAACAA
GATCACCGGCCTGAACTACCGGATCATGCAGAGCGCCATTGCCGTGTACGATAGCGGCGACGATTACGAGGCCAAGCAGCAGTTCAAGCTGATGTTC
GAGAAGGCCCGGCTGATCGGCAAGGGCACAACAGAGCCTCATCCATTTCTGTACAAGGTGTTCGCCCGCAGCATCCCCGCCAATGCCGTCGAGTTCT
ACGAGCGCTACCTGATCGAGCGGAAGTTCTACCTGACCGGCCTGTCCAACGAGATCAAGAAAGGCAACAGAGTGGATGTGCCCTTCATCCGGCGGGA
CCAGAACAAGTGGAAAACACCCGCCATGAAGACCCTGGGCAGAATCTACAGCGAGGATCTGCCCGTGGAACTGCCCAGACAGATGTTCGACAATGAG
ATCAAGTCCCACCTGAAGTCCCTGCCACAGATGGAAGGCATCGACTTCAACAATGCCAACGTGACCTATCTGATCGCCGAGTACATGAAGAGAGTGC
TGGACGACGACTTCCAGACCTTCTACCAGTGGAACCGCAACTACCGGTACATGGACATGCTTAAGGGCGAGTACGACAGAAAGGGCTCCCTGCAGCA
CTGCTTCACCAGCGTGGAAGAGAGAGAAGGCCTCTGGAAAGAGCGGGCCTCCAGAACAGAGCGGTACAGAAAGCAGGCCAGCAACAAGATCCGCAGC
AACCGGCAGATGAGAAACGCCAGCAGCGAAGAGATCGAGACAATCCTGGATAAGCGGCTGAGCAACAGCCGGAACGAGTACCAGAAAAGCGAGAAAG
TGATCCGGCGCTACAGAGTGCAGGATGCCCTGCTGTTTCTGCTGGCCAAAAAGACCCTGACCGAACTGGCCGATTTCGACGGCGAGAGGTTCAAACT
GAAAGAAATCATGCCCGACGCCGAGAAGGGAATCCTGAGCGAGATCATGCCCATGAGCTTCACCTTCGAGAAAGGCGGCAAGAAGTACACCATCACC
AGCGAGGGCATGAAGCTGAAGAACTACGGCGACTTCTTTGTGCTGGCTAGCGACAAGAGGATCGGCAACCTGCTGGAACTCGTGGGCAGCGACATCG
TGTCCAAAGAGGATATCATGGAAGAGTTCAACAAATACGACCAGTGCAGGCCCGAGATCAGCTCCATCGTGTTCAACCTGGAAAAGTGGGCCTTCGA
CACATACCCCGAGCTGTCTGCCAGAGTGGACCGGGAAGAGAAGGTGGACTTCAAGAGCATCCTGAAAATCCTGCTGAACAACAAGAACATCAACAAA
GAGCAGAGCGACATCCTGCGGAAGatccggaacgccttcgat**cac**aacaattaccccgacaaaggcgtggtggaaatcaaggccctgcctgagatcg
ccatgagcatcaagaaggcctttggggagtacgccatcatgaagggaagtggctacccctacgacgtgcctgactacgcctga

**pUb LwaCas13a + LwaCas13a guide RNA** was made by inserting the LwaCas13a ORF sequence between SpeI and AgeI in **pUb 3xFLAG MCS (No BsmBI) + LwaCas13a guide RNA**:

caaagtgaccaaggtcgacggcatcagccacaagaagtacatcgaagagggcaagctcgtgaagtccaccagcgaggaaaaccggaccagcgagaga
ctgagcgagctgctgagcatccggctggacatctacatcaagaaccccgacaacgcctccgaggaagagaaccggatcagaagagagaacctgaaga
agttctttagcaacaaggtgctgcacctgaaggacagcgtgctgtatctgaagaaccggaaagaaaagaacgccgtgcaggacaagaactatagcga
agaggacatcagcgagtacgacctgaaaaacaagaacagcttctccgtgctgaagaagatcctgctgaacgaggacgtgaactctgaggaactggaa
atctttcggaaggacgtggaagccaagctgaacaagatcaacagcctgaagtacagcttcgaagagaacaaggccaactaccagaagatcaacgaga

acaacgtggaaaaagtgggcggcaagagcaagcggaacatcatctacgactactacagagagagcgccaagcgcaacgactacatcaacaacgtgca
ggaagccttcgacaagctgtataagaaagaggatatcgagaaactgtttttcctgatcgagaacagcaagaagcacgagaagtacaagatccgcgag
tactatcacaagatcatcggccggaagaacgacaaagagaacttcgccaagattatctacgaagagatccagaacgtgaacaacatcaaagagctga
ttgagaagatccccgacatgtctgagctgaagaaaagccaggtgttctacaagtactacctggacaaagaggaactgaacgacaagaatattaagta
cgccttctgccacttcgtggaaatcgagatgtcccagctgctgaaaaactacgtgtacaagcggctgagcaacatcagcaacgataagatcaagcgg
atcttcgagtaccagaatctgaaaaagctgatcgaaaacaaactgctgaacaagctggacacctacgtgcggaactgcggcaagtacaactactatc
tgcaagtgggcgagatcgccacctccgactttatcgcccggaaccggcagaacgaggccttcctgagaaacatcatcggcgtgtccagcgtggccta
cttcagcctgaggaacatcctggaaaccgagaacgagaacggtatcaccggccggatgcggggcaagaccgtgaagaacaacaagggcgaagagaaa
tacgtgtccggcgaggtggacaagatctacaatgagaacaagcagaacgaagtgaaagaaaatctgaagatgttctacagctacgacttcaacatgg
acaacaagaacgagatcgaggacttcttcgccaacatcgacgaggccatcagcagcatcagacacggcatcgtgcacttcaacctggaactggaagg
caaggacatcttcgccttcaagaatatcgcccccagcgagatctccaagaagatgtttcagaacgaaatcaacgaaaagaagctgaagctgaaaatc
ttcaagcagctgaacagcgccaacgtgttcaactactacgagaaggatgtgatcatcaagtacctgaagaataccaagttcaacttcgtgaacaaaa
acatccccttcgtgcccagcttcaccaagctgtacaacaagattgaggacctgcggaataccctgaagtttttttggagcgtgcccaaggacaaaga
agagaaggacgcccagatctacctgctgaagaatatctactacggcgagttcctgaacaagttcgtgaaaaactccaaggtgttctttaagatcacc
aatgaagtgatcaagattaacaagcagcggaaccagaaaaccggccactacaagtatcagaagttcgagaacatcgagaaaaccgtgcccgtggaat
acctggccatcatccagagcagagagatgatcaacaaccaggacaaagaggaaaagaatacctacatcgactttattcagcagattttcctgaaggg
cttcatcgactacctgaacaagaacaatctgaagtatatcgagagcaacaacaacaatgacaacaacgacatcttctccaagatcaagatcaaaaag
gataacaaagagaagtacgacaagatcctgaagaactatgagaagcacaatcggaacaaagaaatccctcacgagatcaatgagttcgtgcgcgaga
tcaagctggggaagattctgaagtacaccgagaatctgaacatgttttacctgatcctgaagctgctgaaccacaaagagctgaccaacctgaaggg
cagcctggaaaagtaccagtccgccaacaaagaagaaaccttcagcgacgagttggaactgatcaacctgctgaacctggacaacaacagagtgacc
gaggacttcgagctggaagccaacgagatcggcaagttcctggacttcaacgaaaacaaaatcaaggaccggaaagagctgaaaaagttcgacacca
acaagatctatttcgacggcgagaacatcatcaagcaccgggccttctacaatatcaagaaatacggcatgctgaatctgctggaaaagatcgccga
taaggccaagtataagatcagcctgaaagaactgaaagagtacagcaacaagaagaatgagattgaaaagaactacaccatgcagcagaacctgcac
cggaagtacgccagacccaagaaggacgaaaagttcaacgacgaggactacaaagagtatgagaaggccatcggcaacatccagaagtacacccacc
tgaagaacaaggtggaattcaatgagctgaacctgctgcagggcctgctgctgaagatcctgcaccggctcgtgggctacaccagcatctgggagcg
ggacctgagattccggctgaagggcgagtttcccgagaaccactacatcgaggaaattttcaatttcgacaactccaagaatgtgaagtacaaaagc
ggccagatcgtggaaaagtatatcaacttctacaaagaactgtacaaggacaatgtggaaaagcggagcatctactccgacaagaaagtgaagaaac
tgaagcaggaaaaaaaggacctgtacatccggaactacattgcccacttcaactacatcccccacgccgagattagcctgctggaagtgctggaaaa
cctgcggaagctgctgtcctacgaccggaagctgaagaacgccatcatgaagtccatcgtggacattctgaaagaatacggcttcgtggccaccttc
aagatcggcgctgacaagaagatcgaaatccagaccctggaatcagagaagatcgtgcacctgaagaatctgaagaaaaagaaactgatgaccgacc
ggaacagcgaggaactgtgcgaactcgtgaaagtcatgttcgagtacaaggccctggaaggaagtggctacccctacgacgtgcctgactacgcctg
a

Guide RNA sequences were then cloned into the corresponding plasmids described above. All guide RNAs used in the study were ordered as paired forward/reverse oligos and the sequences are provided in **Supplementary Tables S1 and S2**. Oligos were phosphorylated and annealed using the following 10 μL reactions:

| | |
|---|---|
| 1 μL | 100 mM Forward oligo |
| 1 μL | 100 mM Reverse oligo |
| 1 μL | 10 mM ATP |
| 1 μL | 10x T4 Polynucleotide Kinase Buffer |
| 0.5 μL | T4 Polynucleotide Kinase (NEB) |
| 5.5 μL | Water |

Reactions were incubated in a PCR machine at 37°C for 30 min, followed by 95°C for 5 min, followed by a ramp down to 25°C (0.1°C/sec). Ligations were then performed using purified plasmids cut with BsmBI.

**Reporter plasmids generated:**

**Hy_pMtnA mCherry SV40** (Addgene #176302) was made by inserting the mCherry ORF between the EcoRV and NotI sites in **Hy_pMT EGFP SV40 pA Sense** (Addgene #69911):

```
ACCATGGTGAGCAAGGGCGAGGAggataacatggccatcatcaaggagttcatgcgcttcaaggtgcacatggagggctccgtgaacggccacgagt
tcgagatcgagggcgagggcgagggccgcccctacgagggcacccagaccgccaagctgaaggtgaccaagggtggccccctgcccttcgcctggga
catcctgtcccctcagttcatgtacggctccaaggcctacgtgaagcaccccgccgacatccccgactacttgaagctgtccttccccgagggcttc
aagtgggagcgcgtgatgaacttcgaggacggcggcgtggtgaccgtgacccaggactcctccctgcaggacggcgagttcatctacaaggtgaagc
tgcgcggcaccaacttcccctccgacggccccgtaatgcagaagaagaccatgggctgggaggcctcctccgagcggatgtaccccgaggacggcgc
cctgaagggcgagatcaagcagaggctgaagctgaaggacggcggccactacgacgctgaggtcaagaccacctacaaggccaagaagcccgtgcag
ctgcccggcgcctacaacgtcaacatcaagttggacatcacctcccacaacgaggactacaccatcgtggaacagtacgaacgcgccgagggccgcc
actccaccggcggCATGGACGAGCTGTACAAGTAG
```

**Hy_pMtnA FFLuc SV40** (Addgene #176299) was made by inserting the firefly luciferase (FFLuc) ORF between the XhoI and NotI sites in **Hy_pMT EGFP SV40 pA Sense** (Addgene #69911):

```
accatggaagatgccaaaaacattaagaagggcccagcgccattctacccactcgaagacgggaccgccggcgagcagctgcacaaagccatgaagc
gctacgccctggtgcccggcaccatcgcctttaccgacgcacatatcgaggtggacattacctacgccgagtacttcgagatgagcgttcggctggc
agaagctatgaagcgctatgggctgaatacaaaccatcggatcgtggtgtgcagcgagaatagcttgcagttcttcatgcccgtgttgggtgccctg
ttcatcggtgtggctgtggccccagctaacgacatctacaacgagcgcgagctgctgaacagcatgggcatcagccagcccaccgtcgtattcgtga
gcaagaaagggctgcaaaagatcctcaacgtgcaaaagaagctaccgatcatacaaaagatcatcatcatggatagcaagaccgactaccagggctt
ccaaagcatgtacaccttcgtgacttcccatttgccacccggcttcaacgagtacgacttcgtgcccgagagcttcgaccgggacaaaaccatcgcc
ctgatcatgaacagtagtggcagtaccggattgcccaagggcgtagccctaccgcaccgcaccgcttgtgtccgattcagtcatgcccgcgacccca
tcttcggcaaccagatcatccccgacaccgctatcctcagcgtggtgccatttcaccacggcttcggcatgttcaccacgctgggctacttgatctg
cggctttcgggtcgtgctcatgtaccgcttcgaggaggagctattcttgcgcagcttgcaagactataagattcaatctgccctgctggtgcccaca
ctatttagcttcttcgctaagagcactctcatcgacaagtacgacctaagcaacttgcacgagatcgccagcggcggggcgccgctcagcaaggagg
taggtgaggccgtggccaaacgcttccacctaccaggcatccgccagggctacggcctgacagaaacaaccagcgccattctgatcacccccgaagg
ggacgacaagcctggcgcagtaggcaaggtggtgcccttcttcgaggctaaggtggtggacttggacaccggtaagacactgggtgtgaaccagcgc
ggcgagctgtgcgtccgtggccccatgatcatgagcggctacgttaacaaccccgaggctacaaacgctctcatcgacaaggacggctggctgcaca
gcggcgacatcgcctactgggacgaggacgagcacttcttcatcgtggaccggctgaagagcctgatcaaatacaagggctaccaggtagccccagc
cgaactggagagcatcctgctgcaacaccccaacatcttcgacgccggggtcgccggcctgcccgacgacgatgccggcgagctgcccgccgcagtc
gtcgtgctggaacacggtaaaaccatgaccgagaaggagatcgtggactatgtggccagccaggttacaaccgccaagaagctgcgcggtggtgttg
tgttcgtggacgaggtgcctaaaggactgaccggcaagttggacgcccgcaagatccgcgagattctcattaaggccaagaagggcggcaagatcgc
cgtggactacaaagaccatgacggtgattataaagatcatgacatcgattacaaggatgacgatgacaagtaa
```

**Hy_pUbi-p63e mCherry SV40** (Addgene #176300) was made by inserting the mCherry ORF between the XbaI and NotI sites in **Hy_pUbi-p63e eGFP SV40** (Addgene #132650):

```
ATGGTGAGCAAGGGCGAGGAggataacatggccatcatcaaggagttcatgcgcttcaaggtgcacatggagggctccgtgaacggccacgagttcg
agatcgagggcgagggcgagggccgcccctacgagggcacccagaccgccaagctgaaggtgaccaagggtggccccctgcccttcgcctgggacat
cctgtcccctcagttcatgtacggctccaaggcctacgtgaagcaccccgccgacatccccgactacttgaagctgtccttccccgagggcttcaag
tgggagcgcgtgatgaacttcgaggacggcggcgtggtgaccgtgacccaggactcctccctgcaggacggcgagttcatctacaaggtgaagctgc
gcggcaccaacttcccctccgacggccccgtaatgcagaagaagaccatgggctgggaggcctcctccgagcggatgtaccccgaggacggcgccct
gaagggcgagatcaagcagaggctgaagctgaaggacggcggccactacgacgctgaggtcaagaccacctacaaggccaagaagcccgtgcagctg
cccggcgcctacaacgtcaacatcaagttggacatcacctcccacaacgaggactacaccatcgtggaacagtacgaacgcgccgagggccgccact
ccaccggcggCATGGACGAGCTGTACAAGTGA
```

**pcDNA3.1(+) mCherry** (Addgene #176301) was made by inserting the mCherry ORF between the HindIII and XbaI sites in **pcDNA3.1(+) eGFP** (Addgene #129020):

```
GccaccATGGTGAGCAAGGGCGAGGAggataacatggccatcatcaaggagttcatgcgcttcaaggtgcacatggagggctccgtgaacggccacg
agttcgagatcgagggcgagggcgagggccgcccctacgagggcacccagaccgccaagctgaaggtgaccaagggtggccccctgcccttcgcctg
ggacatcctgtcccctcagttcatgtacggctccaaggcctacgtgaagcaccccgccgacatccccgactacttgaagctgtccttccccgagggc
```

ttcaagtgggagcgcgtgatgaacttcgaggacggcggcgtggtgaccgtgacccaggactcctccctgcaggacggcgagttcatctacaaggtga
agctgcgcggcaccaacttcccctccgacggccccgtaatgcagaagaagaccatgggctgggaggcctcctccgagcggatgtaccccgaggacgg
cgccctgaagggcgagatcaagcagaggctgaagctgaaggacggcggccactacgacgctgaggtcaagaccacctacaaggccaagaagcccgtg
cagctgcccggcgcctacaacgtcaacatcaagttggacatcacctcccacaacgaggactacaccatcgtggaacagtacgaacgcgccgagggcc
gccactccaccggcggCATGGACGAGCTGTACAAGTGA
